# Quantitative Annotations of T-Cell Repertoire Specificity

**DOI:** 10.1101/2023.01.29.526145

**Authors:** Jiaqi Luo, Xueying Wang, Yiping Zou, Lingxi Chen, Wei Liu, Wei Zhang, Shuai Cheng Li

## Abstract

The specificity of a T-cell receptor (TCR) repertoire determines personalized immune capacity. Existing methods have modelled the qualitative aspects of TCR specificity, while the quantitative aspects remained unaddressed. We developed a package, TCRanno, to quantify the specificity of TCR repertoires. Applying TCRanno to 4,195 TCR repertoires revealed quantitative changes in repertoire specificity upon infections, autoimmunity and cancers. Specifically, TCRanno found cytomegalovirus-specific TCRs in seronegative healthy individuals, supporting the possibility of abortive infections. TCRanno discovered age-accumulated fraction of SARS-CoV2-specific TCRs in pre-pandemic samples, which may explain the aggressive symptoms and age-related severity of COVID-19. TCRanno also identified the encounter of Hepatitis B antigens as a potential trigger of systemic lupus erythematosus. TCRanno annotations showed capability in distinguishing TCR repertoires of healthy and cancers including melanoma, lung and breast cancers. TCRanno may also facilitate single-cell TCRseq+gene expression data analyses by isolating T-cells with the specificity of interest.

## Introduction

Specificity is the hallmark of the adaptive immune response. Each T-cell is identified by its T-cell receptor (TCR), which specifically recognizes an antigen-derived epitope peptide presented by certain major histocompatibility complex (MHC) molecules. Millions of T-cells with their TCRs constitute the T-cell pool i.e., the T-cell repertoire or the TCR repertoire, which protects the host from millions of foreign microorganisms. Advances in single-cell and immune receptor sequencing techniques have enabled the high-throughput, high-resolution determination of TCR sequences, where we began to appreciate the huge diversity of TCR sequences in TCR repertoires (1–3). Since the discovery of TCR sequence diversity, the specificity of different TCR clonotypes has become a fundamental problem. Studying the specificity composition of a T-cell repertoire, i.e., the respective fractions of TCRs specific to different epitopes, antigens and organisms, is essential to understanding how the past and current immune responses shape and maintain a T-cell pool.

Recent advances in *ex vivo* tetramer T-cell labeling technique have allowed high-throughput capture of TCR sequences specific to a given epitope of interest (1, 2), resulting in the accumulation of more than 200k TCR sequences with experimentally verified epitope specificity. However, the speed of verification goes far behind the speed of sequence data generation - currently, over three billion immune receptor sequences have been deposited to the immuneACCESS database. The accumulation of verified paired TCR-epitope sequences has allowed the development of qualitative annotation methods probing the specificity of individual TCRs, such as TCR clustering algorithms (4, 5), TCR distance- or similarity-based matching (6, 7) and TCR-peptide-MHC binding affinity prediction methods (8, 9). However, the quantitative aspects of TCR repertoire specificity, i.e., the specificity composition mentioned above, have been largely untouched. Since T-cell clonal expansion is a common feature in immune responses, a quantitative annotation of the T-cell pool is necessary. Currently there is a lack of tools to address the quantity of TCR repertoire specificity, that is, to generate a quantitative profile of TCR specificity to various epitopes, antigens and organisms.

Here, we developed TCRanno, a package for simultaneous qualitative and quantitative annotations of the specificity composition of TCR repertoires. TCRanno demonstrated reliable epitope prediction performance by outperforming Needleman-Wunsch Hamming distance (10), Levenshtein/edit distance, tcrdist (7) and TCRMatch (6) on four benchmark datasets. We then applied TCRanno to peripheral CD8 TCR repertoires from healthy individuals (n=786), SARS-CoV2 exposed/infected individuals (n=1485), pre- or post-vaccination individuals (n=361), systemic lupus erythematosus patients (SLE, n=877) and cancer patients (n=686). TCRanno annotations revealed distinct clonal size distribution and specificity composition in healthy versus diseased repertoires. Within the healthy population, we discovered a significant overlap of known and predicted cytomegalovirus (CMV)-specific TCRs in the CMV^+^ and CMV^−^ sub-populations, indicating CMV exposure but rapid clearance of the virus before seroconversion in some CMV^−^ individuals. In the comparative analyses on repertoires before and during the COVID-19 pandemic, we found age-accumulated fractions of TCRs likely reactive to SARS-CoV2 in pre-pandemic samples. Moreover, the estimated SARS-CoV2-specific TCR fraction was abnormally high in the deceased and the elderly. These findings corresponded to the lymphopenia, cytokine storm and age-related severity of COVID-19. In comparing the specificity composition between healthy versus SLE repertoires, we identified the only significant upregulation of TCR fraction percentage specific to Hepatitis B antigens, suggesting a possible provoking effect on SLE.

Finally, we demonstrated two examples of advanced usage. We showed that a seven-feature representation of TCRanno quantitative annotations distinguishes TCR repertoires of healthy versus infections, autoimmunity and cancers. We also demonstrated that TCRanno qualitative annotations facilitate single-cell TCRseq+gene expression data analyses by isolating cells with the specificity of interest (e.g., SARS-CoV2-specific CD8 T-cells) for precise downstream comparative analyses.

Note that all data used within this study and TCRanno takes as input the CD8 (i.e., MHCI-restricted), TCR *β* CDR3 amino acid sequence. For ease of expression, hereinafter we use TCR sequence and TCR repertoire to denote this specified TCR subtype and the entirety.

## Methods

### Overview of TCRanno Methodology

The major components of TCRanno methodology include the deep-learning model that generates epitope-aware embeddings for TCR sequences, the algorithm for TCR specificity prediction, the process of aggregating TCR clonotype frequencies of a repertoire to obtain the profile of specificity composition, and the reference database. A summary of each component is provided below. See Supplementary Methods for further details.

### Deep-learning model

#### Discriminative feature learning

We designed an encoder-classifier structure (Figure 1a) to allow vector representation of any TCR sequences, and at the same time introduce supervised cognate epitope information during model training. The encoder generates a latent vector (embedding) that captures the discriminative features of the input TCR sequence; the classifier takes the embedding as input and predicts the epitope class of the input TCR sequence. By minimizing the softmax loss for classification, the encoder can learn to generate similar embeddings for similar TCR sequences (i.e., TCR sequences of the same epitope class), such that cosine similarity of the embeddings can be used to measure TCR similarity. We also adapted the center loss (11) to minimize the intraclass variations and maximize the interclass differences of the embedding. The center loss function *L_C_* is given as follow,

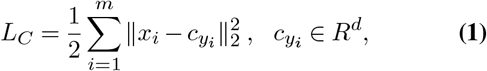

where *m* is the size of the mini-batch and 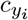 is the feature center of the *y_i_* th epitope class. We set the center loss weight *λ* = 1, meaning an equal weight to the softmax loss. The model was trained for 50 epochs. TCRanno only relies on the encoder to generate the embeddings. The classifier was disposable and no longer required once the training was finished.

**Figure 1.**
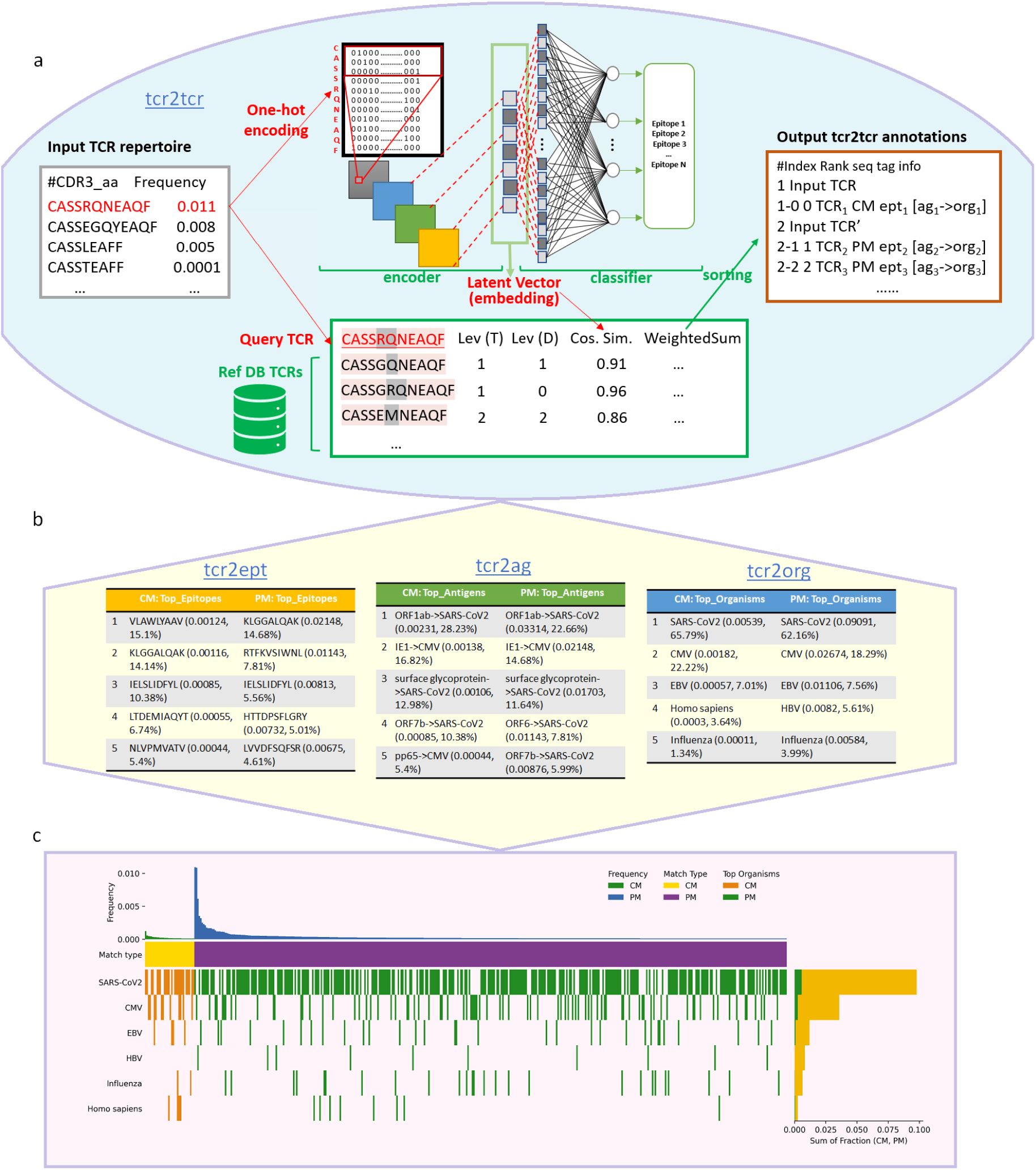
Schematic overview of the workflow of TCRanno: qualitative annotations, quantitative annotations and plot visualization. **(a)** TCRanno qualitative annotations predict the specificity of individual TCR sequences. Each TCR sequence within the input repertoire file is subjected to matching against the reference database. A small portion of the input TCR sequences can find a complete match (CM) sequence within the reference database; TCRanno directly returns the matched sequence and the corresponding epitope, antigen and organism information. Most input TCR sequences cannot find an identical sequence within the database; TCRanno finds and returns the most similar database sequence(s) (predicted match, PM) with the epitope-antigen-organism information. To measure TCR pairwise similarity, TCRanno uses an ensemble similarity score, which is the weighted sum of three similarity scores derived from i) Levenshtein distance of trimmed TCR sequences, ii) Levenshtein distance of TCR D segments and iii) cosine similarity of latent vectors (embeddings) that captures the discriminative features of TCR sequences. **(b)** TCRanno quantitative annotations aggregate TCR clonotype frequencies to calculate TCR fractions specific to epitopes, antigens and organisms. The completely matched (CM) and predicted match (PM) TCRs are calculated and presented separately. **(c)** Graphical summary of the input repertoire’s specificity composition against top organisms. Each column is a TCR clonotype; each row is an organism. The top panel shows the frequency for each clonotype. Clonotypes are ordered according to match type (CM, PM) and clonotype frequency; organisms are ordered according to specific TCR fractions. The right panel shows the specific TCR fractions (CM, PM) for each organism (row).

#### Data Augmentation

Although we have around 200k labeled TCR sequences, the classes are heavily imbalanced - some epitope classes have more than 10k TCR sequences while some have only one or two. To this end, we first performed data-cleaning by removing epitope classes with less than 10 TCR sequences, resulting in a total of 198,842 unique TCR sequences against 382 epitope classes as the training data. Then we performed data augmentation. Briefly, the original training TCR sequences were mutated, inserted, and deleted at one or two positions except for the three amino acids at both ends, to generate all possible sequence combinations that were 1-2 positions different from the original sequences. Then the newly generated sequences were randomly chosen to make up to 50,000 sequences (including the original sequences) for each epitope class.

### TCR specificity prediction

TCRanno predicts the specificity of a query (input) TCR sequence by finding its most similar counterpart within a reference database (Figure 1a). The reference database contains known TCR sequences with verified cognate epitopes. The most similar reference TCR sequence is assumed to share epitope specificity with the query TCR sequence. Hence the prediction algorithm consists of two parts: calculating the pairwise similarity score of the query TCR sequence to each of the reference TCR sequences and inferring the specificity of the query TCR sequence from the top-scored reference TCR sequence.

#### Measuring TCR similarity

We developed an ensemble similarity metric called tcr2tcr score, to measure the similarity of the query TCR sequences to each of the reference TCR sequences. Tcr2tcr score is a self-tuned weighted sum of three similarity scores s_*T*_, s_*D*_ and s_*E*_ as specified below that incorporates supervised deep-learning features and unsupervised distance metrics. For ease of expression, we use *τ_q_* to denote a query TCR sequence and *τ_i_* (*i ∈ R*) to denote a TCR sequence within the reference database.

s_*T*_ measures the similarity of trimmed TCR sequences by Levenshtein distance. Trimming of heads and tails of the TCR sequences is performed to focus on center positions (Supplementary Methods). The pairwise score of the query TCR sequence *τ_q_* to each reference TCR sequence *τ_i_*, denoted as 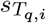, is specified as

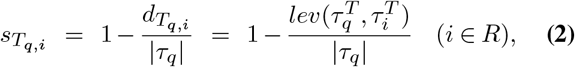

where 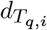 is the Levenshtein distance of the trimmed query and reference TCR sequences 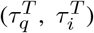, and |*τ*| is the length of the query TCR sequence. If the length of the reference TCR sequence is more than twice of the query TCR sequence, 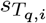 will be less than 0; otherwise, 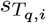 is ranged 0 to 1. The smaller the Levenshtein distance the larger the 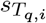 score. Reference TCR sequences with the top *n* 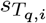 scores to the query are pooled as candidates, in order to ensure the elected reference TCR sequence(s) is similar to the query at the sequence level. We tested a series of *n* values (Supplementary Figure S1) and found the best performance and speed were obtained at *n* = 100. The rest of the calculations described below considers the top-100 pool of reference TCR sequences (denoted as *R_pool_*).

s_*D*_ measures the similarity of D segments within the TCR sequences by Levenshtein distance. A TCR sequence consists of V, D, and J segments citegiudicelli2005imgt. The V and J segments can be obtained by mapping to a map of known human V/J segments (Supplementary Methods). The D segment can then be obtained by subtracting the V and J segments. The pairwise score of *τ_q_* to *τ_i_*, denoted as 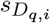, is specified as

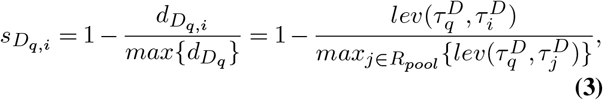

where 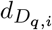 is the Levenshtein distance of the D segments in the query and each reference TCR sequence 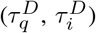, and 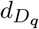 is the array of all 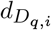. The range of 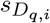 is 0 to 1.

s_*E*_ is the cosine similarity of the embedding vectors generated by the encoder (denoted as *ϵ*). For two TCR sequences with the same specificity, the encoder is expected to generate similar embedding vectors with high cosine similarity. The pairwise score of *τ_q_* to *τ_i_*, denoted as 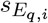, is specified as

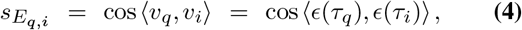

where *v_q_* is the embedding of the query TCR sequence *τ_q_*, and *v_i_* is the embedding of the reference TCR sequence *τ_i_*. The range of 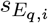 is −1 to 1.

Finally, the tcr2tcr score *S_q,i_* calculates the weighted sum of the three similarity scores.

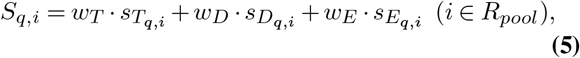

where the weights (*w_T_*, *w_D_*, *w_E_*) of the three scores are their reversed proportion of variances (Supplementary Methods). The self-adapted weights can keep terms from getting too large.

#### Inferring epitope specificity

W ithin t he t op-100 p ool of reference TCR sequences, the one with the top *S_q,i_* score will be selected and its cognate epitope will be considered as the cognate epitope of the query TCR sequence. If the query TCR sequence is 100% identical to the selected reference TCR sequence, we call it a complete match (CM); otherwise, we call it a predicted match (PM). The inferred epitope specificity for each query TCR sequence will be used in the downstream process of aggregating TCR clonotype frequencies.

Alternatively, if sometimes users only desire qualitative annotations, i.e., predicting the specificity of individual TCR sequences, the reference TCR sequences with top K (user-defined) *S_q,i_* scores will be selected for output.

### Aggregating TCR clonotype frequencies

After predicting specificity for input TCR sequences, TCRanno aggregates clonotype frequencies of TCRs that have been predicted to share the same cognate epitope. Since an epitope belongs to an antigen and an organism, clonotype frequencies of TCRs can be further aggregated at antigen and organism levels. Eventually, we obtain the full profile (*P*) of TCR repertoire specificity at the three levels: epitope (*P_E_*), antigen (*P_A_*) and organism (*P_O_*).

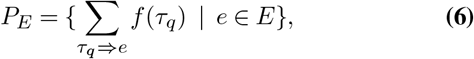

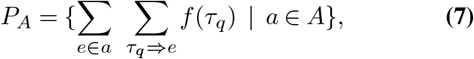

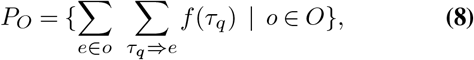

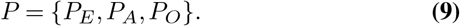

Here we use f(*τ_q_*) to denote the clonotype frequency of the query TCR *τ_q_*. We use *τ_q_* ⇒ *e* to denote that the epitope *e* is cognate to the query TCR sequence, *e ∈ a* to denote that the epitope *e* belongs to the antigen *a*, and *e ∈ o* to denote that the epitope *e* belongs to the organism *o*. We denote *E*, *A* and *O* as the universe of epitopes, antigens and organisms, respectively.

Since the epitope specificity of the CM sequences has been experimentally verified while that of the PM sequences has not, the former has a higher reliability. Hence the CM and PM sequences are aggregated and presented separately to preserve information at varied confidence levels.

### Reference database

The full reference database of TCRanno contains 200,769 known MHCI-restricted, experimentally verified TCR-epitope amino acid sequence pairs extracted from four databases IEDB (12), VDJDB (13), McPAS (14), ImmuneCODE (15) and the peptide-MHC multimer dataset by 10X Genomics (2) (Table 1). As mentioned above, most TCR sequences within the full reference database were used for model training. However, SARS-CoV2-specific TCR sequences constituted more than 89% of the total sequences in the full reference database (Supplementary Figure S2) which could strongly affect the specificity prediction. Therefore, the full reference database is not used as TCRanno’s default reference database. Instead, TCRanno’s default is the IEDB benchmark reference database (see Supplementary Methods for details) where the performance has been benchmarked. We used it in the subsequent application analyses on the 4,195 TCR repertoires.

**Table 1.**
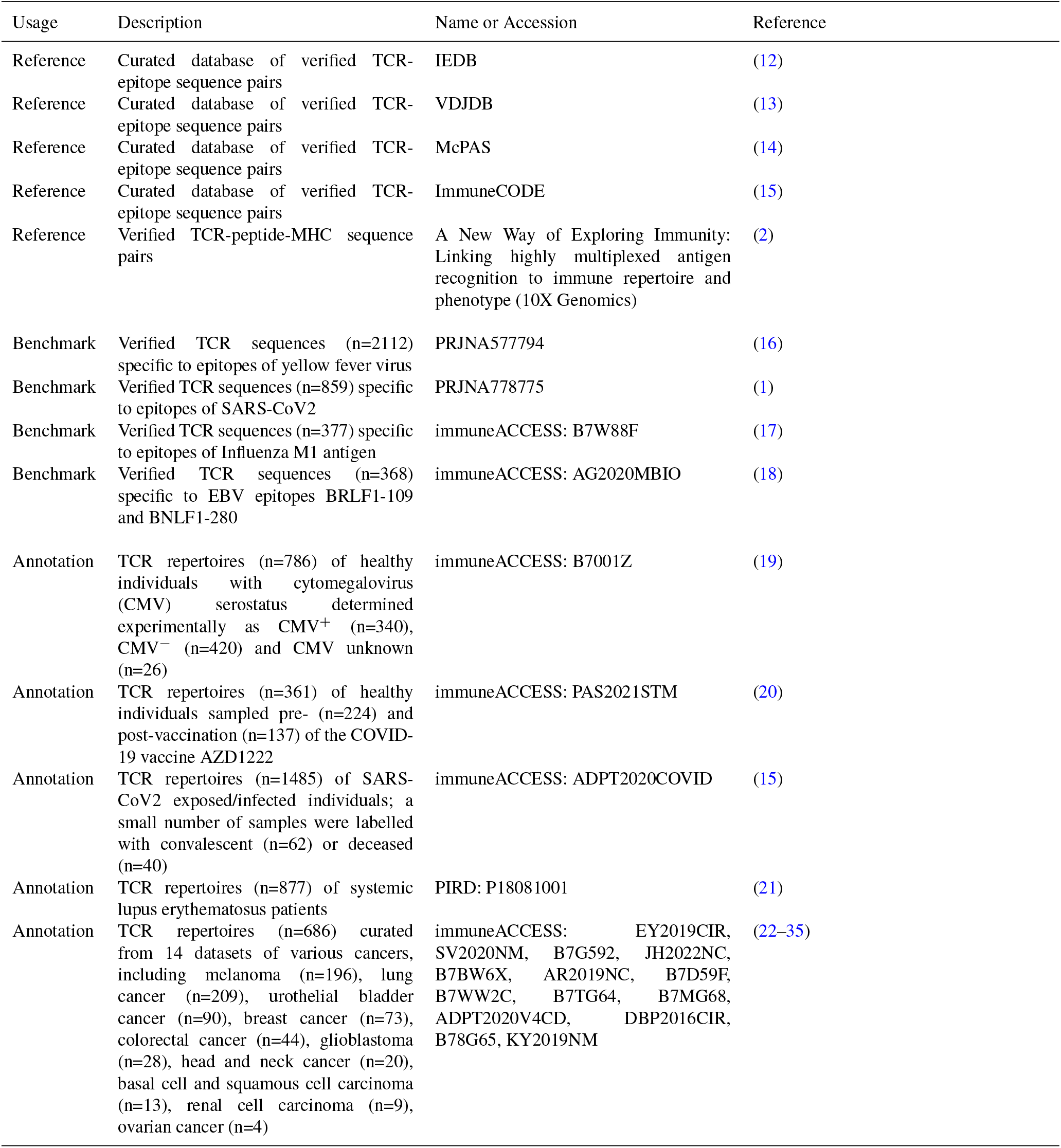
Databases and datasets used in this study.

| Usage | Description | Name or Accession | Reference |
| --- | --- | --- | --- |
| Reference | Curated database of verified TCR-epitope sequence pairs | IEDB | (12) |
| Reference | Curated database of verified TCR-epitope sequence pairs | VDJDB | (13) |
| Reference | Curated database of verified TCR-epitope sequence pairs | McPAS | (14) |
| Reference | Curated database of verified TCR-epitope sequence pairs | ImmuneCODE | (15) |
| Reference | Verified TCR-peptide-MHC sequence pairs | A New Way of Exploring Immunity: Linking highly multiplexed antigen recognition to immune repertoire and phenotype (10X Genomics) | (2) |
| Benchmark | Verified TCR sequences (n=2112) specific to epitopes of yellow fever virus | PRJNA577794 | (16) |
| Benchmark | Verified TCR sequences (n=859) specific to epitopes of SARS-CoV2 | PRJNA778775 | (1) |
| Benchmark | Verified TCR sequences (n=377) specific to epitopes of Influenza M1 antigen | immuneACCESS: B7W88F | (17) |
| Benchmark | Verified TCR sequences (n=368) specific to EBV epitopes BRLF1-109 and BNLF1-280 | immuneACCESS: AG2020MBIO | (18) |
| Annotation | TCR repertoires (n=786) of healthy individuals with cytomegalovirus (CMV) serostatus determined experimentally as CMV<sup>+</sup> (n=340), CMV<sup>-</sup> (n=420) and CMV unknown (n=26) | immuneACCESS: B7001Z | (19) |
| Annotation | TCR repertoires (n=361) of healthy individuals sampled pre- (n=224) and post-vaccination (n=137) of the COVID-19 vaccine AZD1222 | immuneACCESS: PAS2021STM | (20) |
| Annotation | TCR repertoires (n=1485) of SARS-CoV2 exposed/infected individuals; a small number of samples were labelled with convalescent (n=62) or deceased (n=40) | immuneACCESS: ADPT2020COVID | (15) |
| Annotation | TCR repertoires (n=877) of systemic lupus erythematosus patients | PIRD: P18081001 | (21) |
| Annotation | TCR repertoires (n=686) curated from 14 datasets of various cancers, including melanoma (n=196), lung cancer (n=209), urothelial bladder cancer (n=90), breast cancer (n=73), colorectal cancer (n=44), glioblastoma (n=28), head and neck cancer (n=20), basal cell and squamous cell carcinoma (n=13), renal cell carcinoma (n=9), ovarian cancer (n=4) | immuneACCESS: EY2019CIR, SV2020NM, B7G592, JH2022NC, B7BW6X, AR2019NC, B7D59F, B7WW2C, B7TG64, B7MG68, ADPT2020V4CD, DBP2016CIR, B78G65, KY2019NM | (22–35) |

### Datasets

#### Benchmark datasets

We collected four experimentally verified datasets (Table 1; see Supplementary Methods for data preparation details) of viral epitope-specific TCR sequences to evaluate the epitope prediction performance of TCRanno and four state-of-the-art approaches: TCRMatch (6), Needleman-Wunsch Hamming distance (10), tcrdist and Levenshtein/edit distance (7). The four benchmark datasets contain TCR sequences specific to epitopes of the yellow fever virus (YFV, n=2112), SARS-CoV2 (n=859), Influenza (flu, n=377) and Epstein-Barr virus (EBV, n=368).

#### Annotation datasets

We performed TCR repertoire annotations and multi-group comparisons on a total of 4,195 repertoires curated from 18 studies (Table 1; see Supplementary Methods for data preparation details). The repertoires were all sampled from periphery blood mononuclear cells (PBMC) and classified into five groups of different conditions. Group 1: healthy individuals with different CMV serostatus (n=786). Group 2: healthy individuals sampled pre- and post-vaccination of the COVID-19 vaccine AZD1222 (n=361). Group 3: SARS-CoV2 exposed/infected individuals (n=1485). Group 4: SLE patients (n=877). Group 5: patients of various cancer types (n=686).

### Benchmarking performance of TCR specificity prediction

We benchmarked TCRanno’s performance on specificity along with four state-of-the-art methods: TCRMatch (6), Needleman-Wunsch Hamming distance (hereinafter denoted as nwhamming) (10), tcrdist and Levenshtein/edit distance (7), on the four benchmark datasets. We downloaded and installed TCRMatch from the IEDB Analysis Resources, and we set the filtering level as ‘Low’ (0.84) to allow as many matches as possible for performance optimization. The tcrdist, nwhamming and Levenshtein metrics were imported from the pwseqdist package. Pair-wise distance matrix for each metric was computed using the rep_funcs module in tcrdist3. We then tested each method on finding the top K (K=1, 2, 3, 4, 5, 6, 7, 8, 9, 10) most similar TCR sequence(s) within the benchmark reference database for each test TCR sequence. We determined the hit rate as the rate of correctly finding TCR sequence(s) recognizing the same epitope as with the test TCR sequence.

For runtime evaluation, we generated TCR repertoires of size 100, 500, 1,000, 5,000, 10,000, 50,000 and 100,000 by random sampling from a large pool of TCR sequences. We generated the large TCR pool by extracting sequences (taking the top 10,000 sequences per repertoire) from 587 TCR repertoires of healthy individuals (36). We evaluated runtime for all methods using 8-CPUs (i7-9700) on each tested repertoire size for 5 repeats, except for tcrdist, nwhamming and Levenshtein on sizes 50,000 and 100,000 which only performed once due to long runtime.

### Multi-group comparative analyses of repertoire specificity composition

We first performed TCRanno qualitative and quantitative annotations to each of the 4,195 TCR repertoires. For epitopes, antigens and organisms of interest, we visualized the distribution of specific TCR fractions as boxplots for each population and/or sub-population. We performed the Mann-Whitney U test to validate the differences in TCR fraction distribution between groups of interest. We further calculated the mean specificity composition (i.e., the mean TCR fraction specific to epitopes, antigens and organisms) for each population and/or sub-population. For the SLE vs healthy comparative study, we computed bar plots of the mean TCR fraction and the mean TCR fraction percentage specific to major organisms and antigens.

### Distinguishing repertoires of different conditions and visualization on 2D space

To distinguish repertoires of healthy versus pathological (e.g. cancers, autoimmunity, infection) conditions, we selected a panel of seven numeric features from TCRanno’s quantitative annotation output: the fractions of TCR specific to the top five microorganisms: SARS-CoV2, CMV, Influenza, EBV and Hepatitis B virus (HBV), as well as the medium-frequency clonotype fraction and the high-frequency clonotype fraction which sum up to the effective fraction. The effective fraction is defined as the sum of clonotype frequencies of TCRs with clonotype frequency no less than 10^−4^; the medium-frequency clonotype fraction is defined as the sum of clonotype frequencies of TCRs with clonotype frequency no less than 10^−4^ and less than 10^−2^; the high-frequency clonotype fraction is defined as the sum of clonotype frequencies of TCRs with clonotype frequency no less than 10^−2^. For each individual repertoire, we computed the seven numeric features as a seven-dimension vector from their TCRanno annotation output. We applied SEAT (Structure Entropy hierArchy Detection) (37) and UMAP (Uniform Manifold Approximation and Projection) (38) for dimensionality reduction, and visualized the 2D separation of repertoires under different conditions using scatter plot of the first two dimensions of the SEAT or UMAP embeddings.

### TCRanno-aided single-cell TCRseq+gene expression data analyses

Seurat objects for a pediatric COVID-19 (pcovid) dataset (39) and an adult COVID-19 (acovid) dataset (40) were downloaded from the original studies. The acovid data consists of experimentally enriched SARS-CoV2-specific CD8 T-cells from PBMCs, while the pcovid data contains non-enriched CD8 T-cells. To isolate SARS-CoV2-specific T-cells in the pcovid data, we performed TCRanno qualitative annotation to predict the specificity of each TCR sequence, then we selected those T-cells with TCR sequences specific to SARS-CoV2. Then we used the DEenrichRPlot function in Seurat (41) with the ’GO Biological Process 2018’ database, to perform differential expression and enrichment analysis on the TCRanno-isolated T-cells of the pcovid data and the experimentally-isolated T-cells in the acovid data.

## Results

We developed TCRanno, a toolkit for annotating, quantifying and visualizing the specificity composition of TCR repertoires at epitope, antigen and organism levels (Figure 1). TCRanno consists of two steps: i) qualitative annotations - predicting the specificity of individual TCR sequences (Figure 1a), and ii) quantitative annotations - aggregating TCR clonotype frequencies to obtain a profile of TCR repertoire specificity at the three levels (Figure 1b). For each TCR sequence within the input repertoire, TCRanno predicts specificity by searching a large reference database and returning an identical (complete match, CM) or the most similar known TCR sequence (predicted match, PM), which is assumed to share specificity with the input TCR sequence.

The subsequent aggregating process provides an estimation of TCR fractions specific to different epitopes, antigens and organisms (Figure 1b). The specificity landscape plots powered by Comut (42) offer graphical summaries of the input TCR repertoire’s specificity composition at the three levels (Figure 1c; Supplementary Figure S3). TCRanno also supports multi-repertoire annotations to allow group comparisons.

### TCRanno outperformed state-of-the-art approaches on TCR specificity prediction

Performance of TCR specificity prediction by TCRanno, TCRMatch, tcrdist, Levenshtein distance and Needleman-Wunsch Hamming distance (nwhamming) were compared on four independent benchmark datasets (Table 1). The datasets contained TCR sequences specific to epitopes derived from i) yellow fever virus (YFV, n=2112); ii) SARS-CoV2 (n=859); iii) Influenza (flu, n=377) and iv) Epstein-Barr virus (EBV, n=368). Performance was determined by the hit rate, which is the rate of correctly finding the most similar TCR sequence(s) recognizing the same epitope as the test TCR sequence. The hit rates over top K choices (see Methods) by each method on the four datasets were plotted and compared (Supplementary Figure S4). TCRanno’s core algorithm, tcr2tcr (see Methods), ranked at the top for all the benchmark datasets, showing versatile generalizability. Importantly, tcr2tcr demonstrated advantages at K=1, which gives an indication of TCRanno’s annotation accuracy since the quantitative annotations only consider a single best match for each input TCR sequence. We further verified that the encoder-generated embeddings for the benchmark TCR sequences showed significant intraclass aggregation at the latent vector space (Supplementary Figure S5). Supplementary Table S1 shows the runtime of each method using 8-CPUs (i7-9700) on various input repertoire sizes ranging from 100 to 100,000 (5 repeats per method per size). Tcr2tcr handled small input sizes (100-500) in less than 30 seconds and ran the fastest on medium and large input sizes, finishing annotations of 100,000 TCR sequences in less than 18 minutes.

### Clonotype frequency distributions differed between healthy and pathological conditions

We then applied TCRanno to a total of 4,195 peripheral CD8 TCR repertoires from multiple datasets (Table 1) representing populations under five different conditions: i) healthy individuals with different CMV serostatus (n=786, denoted as ‘Healthy group’), ii) SARS-CoV2 exposed/infected individuals (n=1485, denoted as ‘SARS-CoV2 group’), iii) COVID-19 AZD1222 pre- and post-vaccinated individuals (n=361, denoted as ‘Vaccine group’), iv) SLE patients (n=877, denoted as ‘SLE group’) and v) cancer patients (n=686, denoted as ‘Cancer group’). We first examined the distribution of repertoire clonal size (i.e., clonotype frequency) and productive fraction (i.e., sum of clonotype frequencies of in-frame TCRs) among different populations. Significant differences were observed in the clonotype frequency distributions (Figure 2a). While the two healthy populations (Healthy, Vaccine) were dominated by low-frequency clonotypes (with clonotype frequency less than 10^−4^), the three disease populations (SARS-CoV2, Cancer, SLE) exhibited a shift to medium (10^−4^ ≤ frequency < 10^−2^) or high-frequency clonotypes (frequency ≥ 10^−2^), with SLE being extremely scarce on low-frequency clonotypes. All populations showed similar levels of productive fraction.

**Figure 2.**
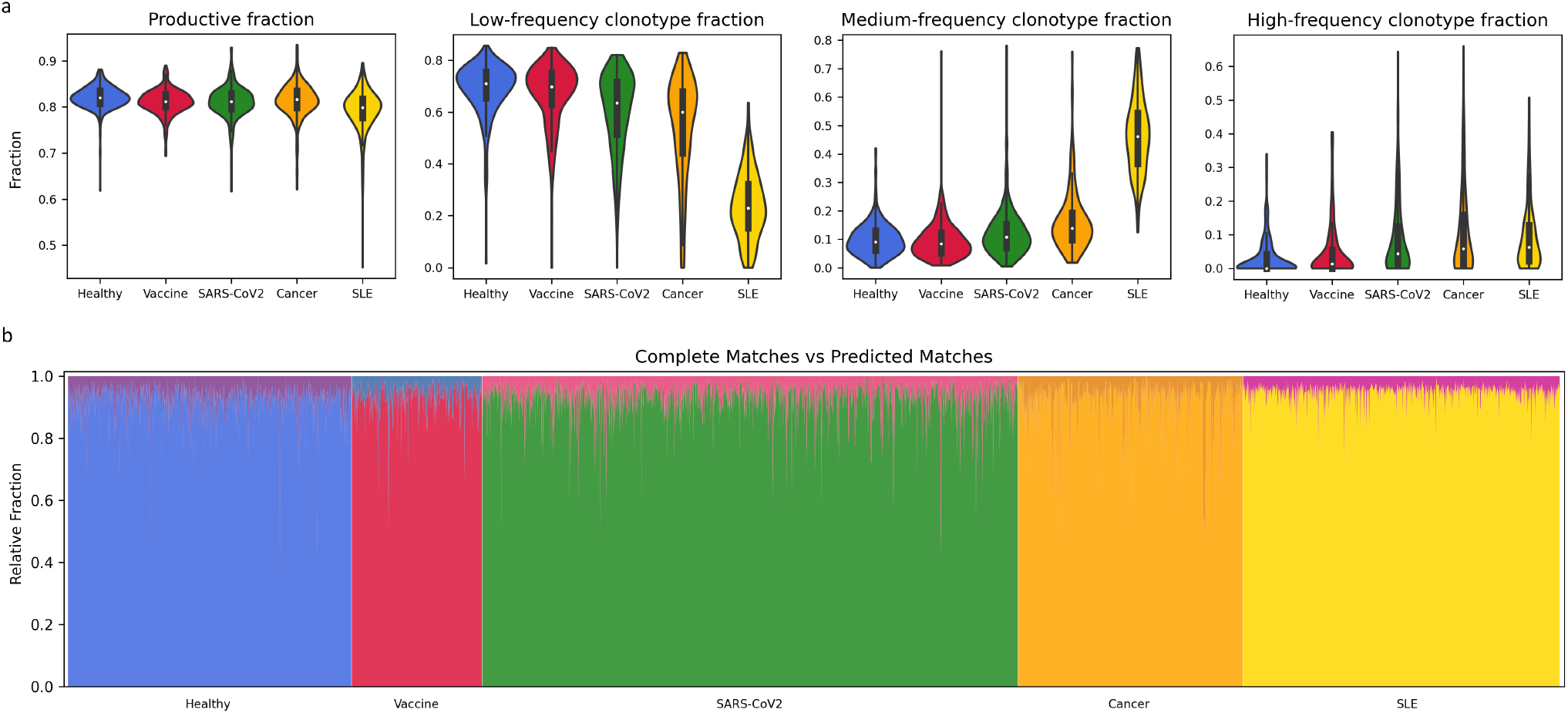
Distributions of clonal size and productive fraction in different populations. **(a)** Violin plots of productive fraction and fractions of low-, medium- and high-frequency clonotypes for the five conditions: i) healthy individuals with different CMV serostatus (n=786), ii) SARS-CoV2 exposed/infected individuals (n=1485), iii) COVID-19 AZD1222 pre- and post-vaccinated individuals (n=361), iv) SLE patients (n=877) and v) cancer patients (n=686). **(b)** Relative fraction of TCR sequences being complete matches (CM) versus predicted matches (PM) within the effective fraction. The CM and PM relative fractions sum to 1. The calculated mean CM relative fraction was 0.070 (7.0%).

Since almost all naïve TCRs are less than 10^−4^ frequency (3), we excluded the low-frequency clonotypes in this study to focus on the likely memory portion because most naïve T-cells will not recognize an epitope till die. We defined the effective fraction of a repertoire as the sum of frequencies of all productive TCR clonotypes with ≥ 10^−4^ frequency.

Within the effective fraction, known TCR sequences were detected in 4,187 out of the 4,195 (99.8%) repertoires, i.e., completely matches (CM) to the reference database. However, the fraction of unknown TCR sequences, i.e., predicted matches (PM) dominated all repertoires examined, accounting for 93.0% of the effective fraction on average within a repertoire (Figure 2b).

### CMV-specific TCRs identified in CMV^−^ repertoires

To investigate how CMV affects the host’s T-cell pool, we analyzed quantitative differences in repertoire specificity composition between CMV^+^ (n=340) and CMV^−^ (n=420) individuals within the Healthy group. We compared the estimated TCR fractions specific to CMV and the two major CMV antigens: IE1 and pp65, between the two sub-populations (Figure 3a; Supplementary Table S2-3). We observed a significant increase in the estimated TCR fractions specific to the virus and the two antigens in CMV^+^ individuals compared to CMV^−^ individuals. However, we also found substantial known and predicted CMV-specific TCRs in CMV^−^ individuals, with a median CM fraction over 0.001 and a median PM fraction over 0.01. Considering an estimate of 5-10 × 10^9^ T-cells in peripheral blood (43) and a normal 2:1 CD4:CD8 ratio, a 0.001 CM fraction means at least 10^6^ known CMV-reactive CD8 T-cells. This indicated at least in some CMV^−^ individuals, the possibility of past exposure but rapid clearance of the virus before seroconversion by pre-existing CMV-reactive T-cells, a phenomenon similar to what has been observed in SARS-CoV2-exposed seronegative individuals (44). The results also showed that IE1-specific TCRs dominated over pp65-specific TCRs in both sub-populations, especially in CMV^−^ individuals.

**Figure 3.**
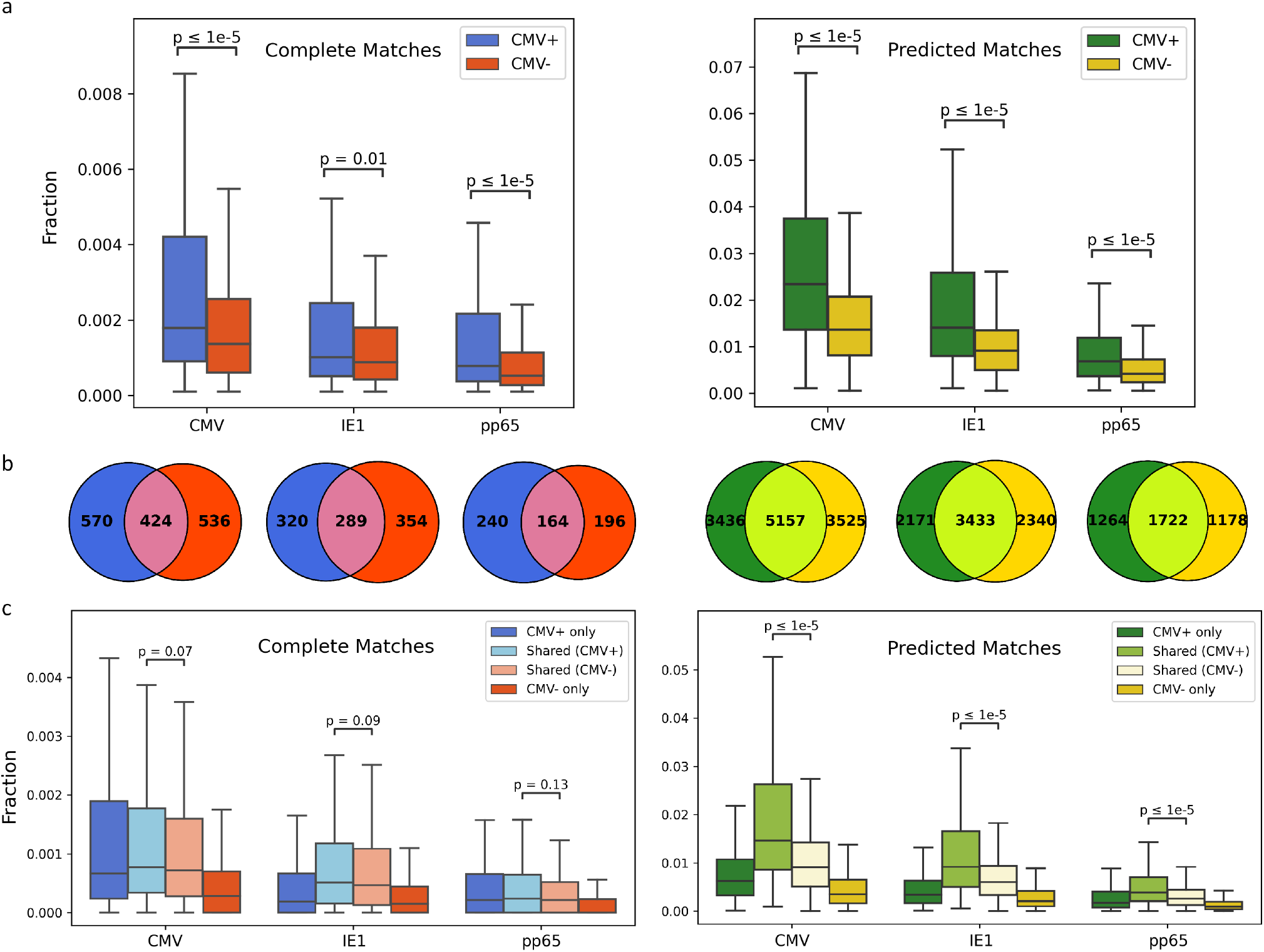
Quantitative differences and overlap in CMV-specific TCRs between CMV^+^ and CMV^−^ individuals. **(a)** Fractions of TCRs specific to CMV and its major antigens IE1 and pp65, in CMV^+^ and CMV^−^ individuals, for both complete matches (left) and predicted matches (right). **(b)** Venn plots showing the number of CMV-specific TCR sequences shared between CMV^+^ and CMV^−^ individuals, for both complete matches (left) and predicted matches (right). **(c)** Fractions of CMV-specific TCRs as being CMV^+^ exclusive, shared (overlapped), CMV^−^ exclusive, for both complete matches (left) and predicted matches (right).

To further examine the differences in T-cell responses between CMV^+^ and CMV^−^ individuals, we extracted the CMV-reactive TCR sequences that were shared or exclusive for both sub-populations and calculated the corresponding TCR fractions (Figure 3b, 3c). Surprisingly, the fraction of shared TCRs dominated over the fraction of exclusive TCRs in both CMV^+^ and CMV^−^ repertoires, suggesting dominating roles of the shared TCRs in the immune responses against CMV in both sub-populations. Moreover, the shared TCRs seem to have constituted a higher fraction in the CMV^+^ group than in the CMV^−^ group. Based on currently available information we could not elucidate whether this higher fraction of shared TCRs in CMV^+^ individuals was formed after the establishment of infection, or whether there existed other factors affecting the functions of these otherwise pre-existing TCRs in CMV^+^ individuals, making them unable to clear the virus just as some of the CMV^−^ individuals presumably did.

### Age-accumulated fraction of SARS-CoV2-specific TCRs widely found in pre-COVID-19 repertoires

Next, we analyzed the 1,485 repertoires from the SARS-CoV2 group and compared them with the 786 repertoires from the Healthy group sampled at least 2 years before the COVID-19 pandemic. Consistent with previous studies (1), the estimated SARS-CoV2-specific TCR fraction was significantly higher in exposed/infected individuals than in pre-pandemic individuals (Figure 4a; Supplementary Figure S6; Supplementary Table S5-9), despite an overall lower T-cell count. Note that due to the sudden rise of SARS-CoV2 research, the reference database was dominated by TCR sequences recognizing SARS-CoV2 antigens (Supplementary Figure S2), making the estimated fractions specific to other viral antigens prone to be underestimated and those to SARS-CoV2 prone to be overestimated. Despite this, SARS-CoV2-specific TCRs were widely and substantially found in the pre-pandemic repertoires with a mean CM fraction of 0.007 and a mean PM fraction of 0.080 (Figure 4a; Supplementary Table S4). These results corresponded to recent findings (44, 45) that pre-existing memory T-cells elicit broad cross-reactivity to SARS-CoV2. Moreover, the SARS-CoV2-specific fraction seemed to accumulate with age - we observed an gradual increase in the estimated SARS-CoV2-specific fraction towards older age groups (Figure 4b) in both the pre-pandemic healthy population and the SARS-CoV2 exposed/infected population. The exceptionally high SARS-CoV2-specific fraction shown in the exposed/infected 0-9 years group was probably because of the group only contained two samples of severely ill (ICU admitted) cases likely representing the rare, multisystem inflammatory syndrome in children, while most pediatric COVID-19 cases were mild and self-limiting (39). The pre-pandemic repertoires of age ≥ 60 were estimated with a median SARS-CoV2-specific fraction (CM+PM) over 0.1, meaning more than 10% of the T-cells could be activated by the virus. If such a high portion of potentially reactive T-cells reside in the peripheral blood, it would be no surprise to see a systemic inflammatory response that leads to cytokine storm, lymphopenia and other aggressive symptoms as observed in COVID-19. Hence these likely pre-existing, age-accumulated SARS-CoV2-reactive TCRs in pre-pandemic individuals may account for the age-related differences in COVID-19 disease outcomes. Furthermore, the estimated fraction of SARS-CoV2-specific TCRs in deceased individuals significantly outweighed that of the convalescents (Figure 4a), in addition to a 3- to 5-fold change (upregulation) of estimated TCR fraction against self (Homo sapiens) and self-antigens such as APAK13, NUF2, MLANA and ABCD3 in deceased individuals, as compared to healthy individuals or the convalescents (Supplementary Table S4, S6-7). These results suggested a deleterious effect of excessive SARS-CoV2-specific T-Cells by inducing autoimmunity.

**Figure 4.**
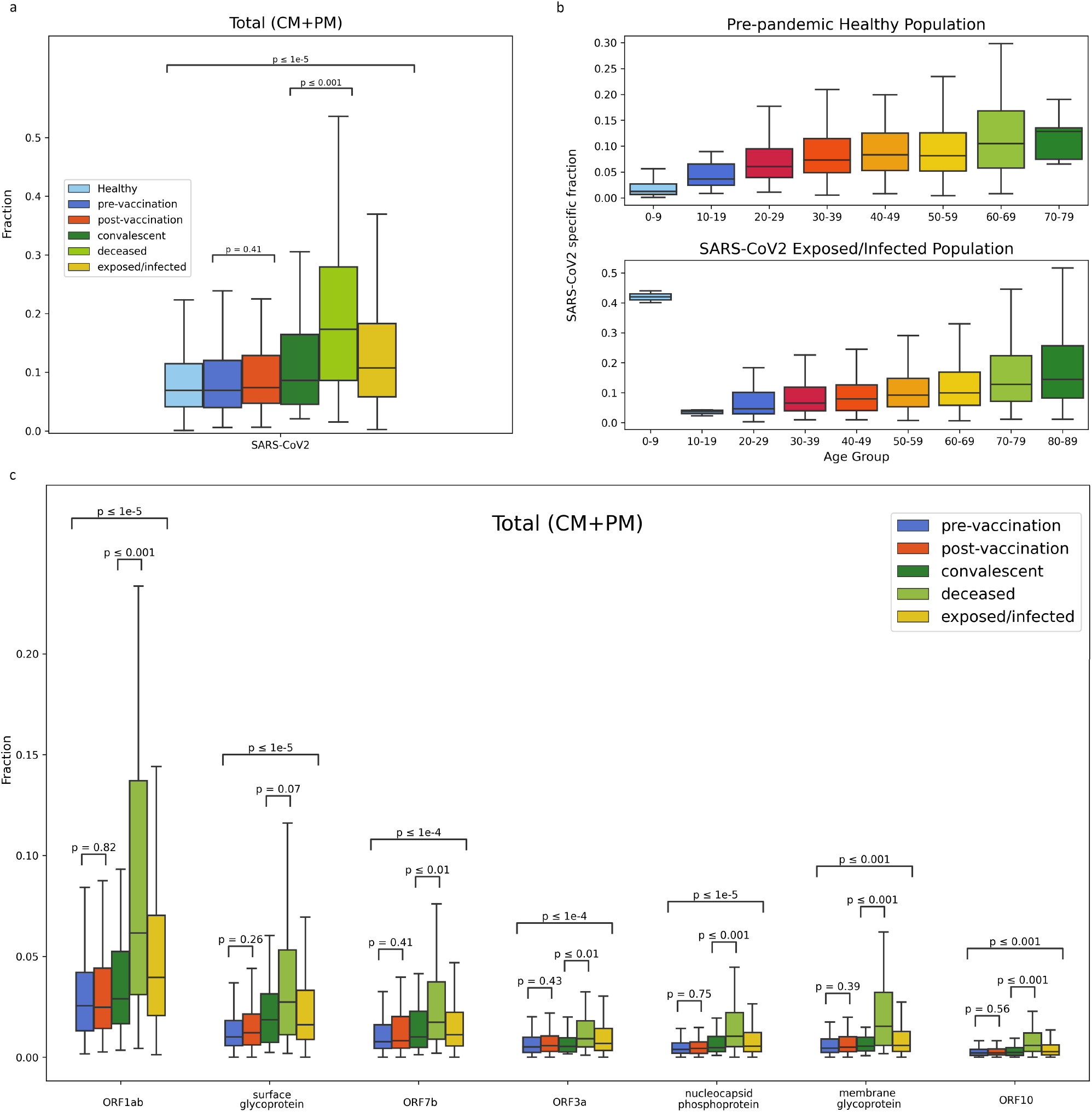
Total fractions (complete matches + predicted matches) of SARS-CoV2-specific TCRs in different populations. **(a)** SARS-CoV2-specific TCR fractions in healthy (n=786), pre-vaccination (n=224), post-vaccination (n=137), SARS-CoV2 convalescent (n=62), deceased (n=40) and exposed/infected (n=1485) individuals. **(b)** SARS-CoV2-specific TCR fractions in different age groups of the healthy group (top) and the exposed/infected group (bottom). **(c)** Fractions of TCRs specific to major antigens of SARS-CoV2 in the pre-vaccination, post-vaccination, SARS-CoV2 convalescent, deceased and exposed/infected sub-populations.

We then examined the estimated fractions of TCRs recognizing major antigens of SARS-CoV2 in different sub-populations. Figure 4c shows SARS-CoV2 antigens with specific TCR fractions (CM+PM) no less than 3% of effective fraction in exposed/infected individuals. Interestingly, TCRs recognizing the conserved ORF1ab antigen dominated all five sub-populations examined (pre-vaccination, post-vaccination, convalescent, deceased and exposed/infected), followed by TCRs specific to surface glycoprotein (spike). This was not likely to be caused by dominating frequency of ORF1ab-specific TCR sequences in the reference database, because the top-3 antigens in the reference database were ORF7b (29.49%), surface glycoprotein (12.70%) and ORF3a (10.79%) (Supplementary Figure S2, bottom right). Although the sampling timepoint of the post-vaccination repertoires (at day 28, right after the second dose) may be too early to capture the boosted T cell responses, we did observe a trend of increase in estimated TCR fraction against surface glycoprotein (spike), the target antigen of the AZD1222 vaccine (Figure 4c).

### Encounter of HBV antigens identified as a possible trigger of SLE onset

Next, we tried to determine what caused the abnormally high effective fraction reflecting a high memory-like portion in SLE (Figure 2a). We calculated the mean TCR fraction specific to major organisms and antigens for the SLE group (n=877) and the Healthy group (n=786). Here we included only the CM TCRs for which specificity have been verified experimentally to maximize the reliability of the analysis. We further converted the respective TCR fractions into percentages by dividing the total CM fraction (Figure 5; Supplementary Table S4, S10). We found that except for HBV which showed a 3-fold change (upregulation) in SLE patients (Figure 5a, top; highlighted in red boxes), the fraction percentage specific to other organisms did not increase and the overall percentage compositions remained similar, despite an apparent increase in the actual fraction against these organisms (Figure 5a, bottom). The significant increase in HBV-specific TCRs came mainly from TCRs specific to HBV antigens derived from the precore/core protein and the DNA polymerase (Figure 5b, highlighted with red boxes), an indication of past or current encounters of HBV antigens. These results correlate with previous findings of the associations between HBV immunization and risks of SLE onset, as well as the higher HBV-carrier incidence (46) and the high risk of HBV reactivation (47) in SLE patients. A sharp increase in effective fraction accompanied by a largely unchanged percentage composition suggested the clonal expansion in SLE as a non-specific, uniform expansion, a typical manifestation of bystander T cell activation.

**Figure 5.**
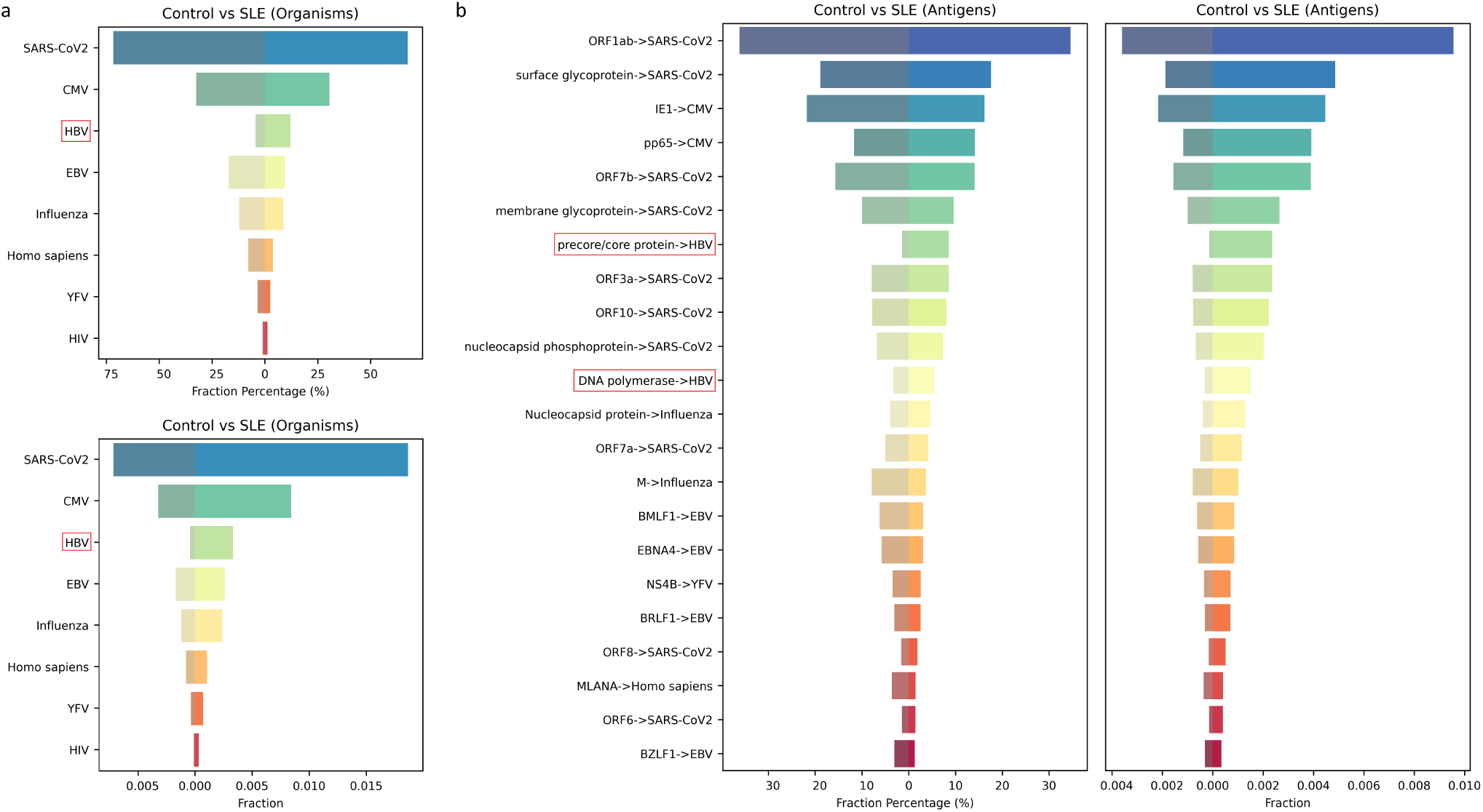
A comparison of TCR repertoire specificity composition represented by mean fraction percentage and mean fraction, between the SLE (n=877) and the Healthy (n=786) group, at **(a)** organism level and **(b)** antigen level. Only completely matched TCR sequences were considered. HBV and HBV antigens (highlighted with red boxes) were the only entries showing significant upregulation in mean fraction percentage.

### Quantitative annotations distinguish repertoires of healthy versus diseases

Next, we tested the possibility of distinguishing repertoires of different conditions (healthy, infection, autoimmunity, cancer, etc.) using the quantitative differences in TCR repertoire specificity. We observed a positive correlation between the effective fraction and the intensity of immune response, which makes effective fraction a strong biomarker for distinguishing not only populations (Figure 2a) but also sub-populations within each population group (Figure 6a; Supplementary Figure S7). For example, the CMV^+^ sub-population held a higher mean effective fraction than the CMV^−^ sub-population within the Healthy group (Figure 6a; Supplementary Table S2-3). The melanoma sub-population, known for being the most immunogenic, was shown to have the highest effective fraction among different cancers (Figure 6a; Supplementary Table S11-15). Strikingly, the urothelial bladder cancer sub-population kept a low-level effective fraction very similar to the healthy populations (Figure 6a; Supplementary Table S4, S14), reflecting the immunosuppressive nature of the type of cancer (48). The autoimmune SLE, known for widespread inflammation, showed the highest effective fraction among all sub-populations studied (Figure 6a; Supplementary Table S10). We also noticed some cancer-related upregulation of TCR fractions specific to certain antigens and organisms. For example, we observed a 7-fold change (upregulation) in TCR fraction (CM) against the self-antigens MLANA and ABCD3 (Supplementary Table S4, S11) in melanoma, and a 7-fold change (upregulation) of Influenza-specific TCR fraction (CM) in breast cancer (Supplementary Table S4, S13). In addition to the anti-tumor effects of viral infection shown in previous studies (49–51), our results in turn suggested that the expanded viral-specific TCRs in repertoires of cancer patients could be the result of an intrinsic anti-tumor response.

**Figure 6.**
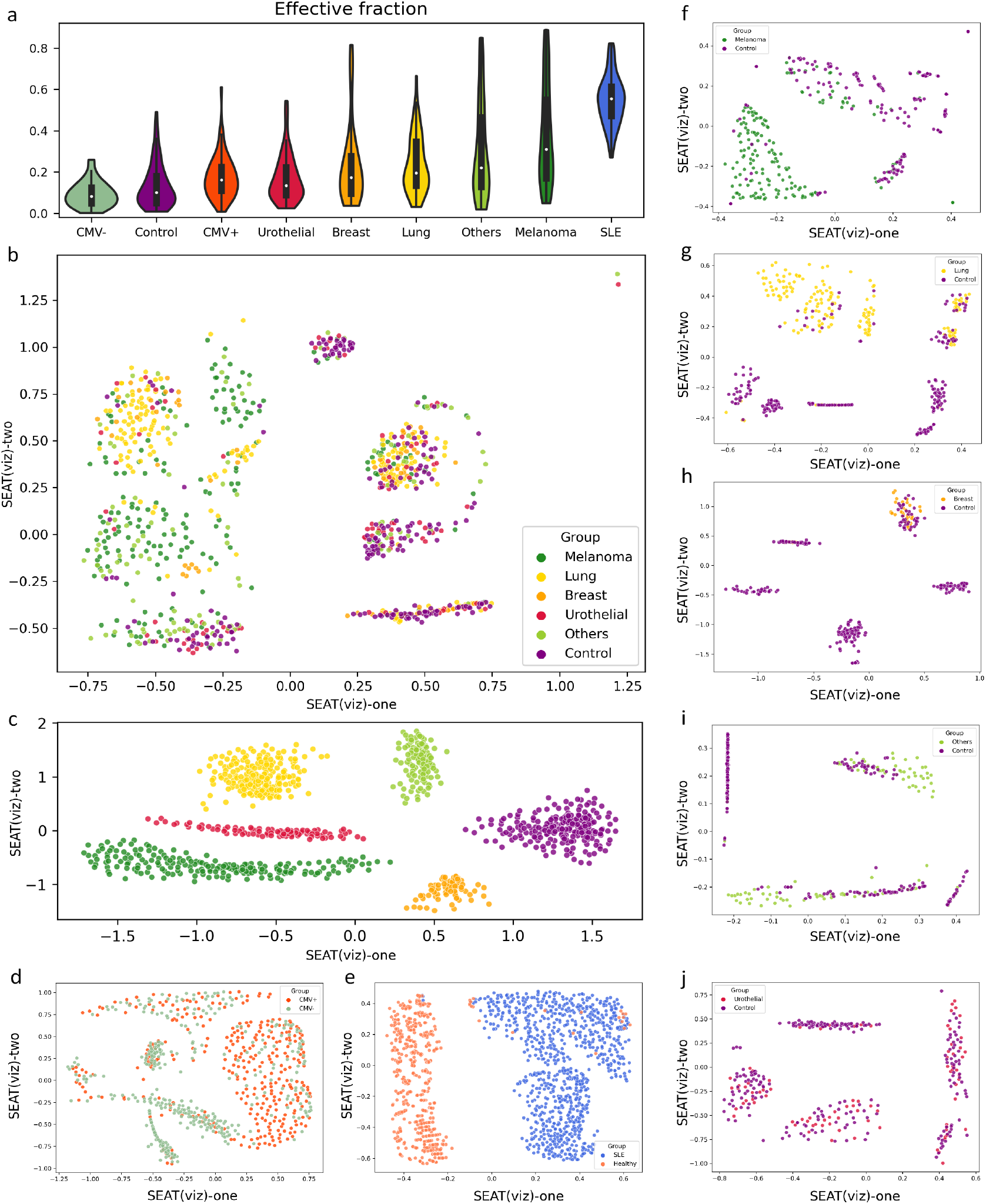
Distinguishing healthy versus pathological repertoires with a seven-feature vector representation of TCRanno annotation output. **(a)** Effective fraction correlates with immune response intensity. In healthy conditions, the CMV^+^ group showed a higher effective fraction than the CMV^−^ group; among cancers, melanoma showed the highest effective fraction and has been known for being the most immunogenic, while urothelial bladder cancer showed a low effective fraction similar to healthy populations; SLE showed an extremely high effective fraction (much higher than that of melanoma) consistent with the widespread inflammation commonly observed. **(b)** scatter plot of the first two dimensions of the *unsupervised* SEAT hierarchical embedding of the seven-feature vector representation of repertoires from various cancers (n=686) vs healthy control (n=224, pre-vaccination). Each point represented an individual repertoire. Different colors were used to specify different conditions. **(c)** scatter plot of the first two dimensions of the *supervised* SEAT hierarchical embedding of the same seven-feature vectors used in **(b)**. **(d-j)** scatter plots of the first two dimensions of the *unsupervised* SEAT hierarchical embedding of the seven-feature vector representation of repertoires from **(d)** CMV^+^ (n=420) vs CMV^−^ (n=340), **(e)** SLE (n=877) vs Healthy (n=786), **(f)** melanoma (n=196), **(g)** lung cancer (n=209), **(h)** breast cancer (n=73), **(i)** other cancers (n=118), **(j)** urothelial bladder cancer (n=90) vs healthy control (n=224), respectively.

Therefore, we hypothesized that repertoires of different sub-populations have different distributions of effective fraction and organism-specific fraction, which can be separated in high-dimensional space. We selected a total of seven numeric features from TCRanno’s quantitative annotation output to represent a repertoire (Supplementary Table S16): the medium-frequency clonotype fraction and the high-frequency clonotype fraction that constitute the effective fraction, plus the TCR fraction specific to each of the major five microorganisms: SARS-CoV2, CMV, Influenza, EBV and HBV. Our results demonstrated that the studied sub-populations were largely separable on dimensionality-reduced 2D space using unsupervised SEAT hierarchical embedding (Figure 6b, 6d-j) or UMAP (Supplementary Figures S8-15), especially in melanoma (n=196) vs control (n=224), lung cancer (n=209) vs control (n=224) and SLE (n=877) vs Healthy (n=786). Consistent with the low effective fraction inseparable from healthy populations (Figure 6a), urothelial bladder cancer was hardly separable in the seven-dimensional vector space (Figure 6j).

### Qualitative annotations isolate cells of interest for single-cell data analyses

Single-cell gene expression coupled with TCRseq allows an integrated study of the transcriptomes associated with individual TCR sequences. However, the subset of T-cells recognizing the organism of interest (e.g., SARS-CoV2-specific T-cells) cannot be easily selected unless extra experimental enrichment. With TCRanno’s specificity prediction, we can tag TCRs with their predicted cognate epitopes, antigens and organisms, then we can isolate T-cells of interest for downstream comparative analyses.

To specifically investigate the roles of SARS-CoV2-specific CD8 T-cells in COVID-19 severe illnesses of both children and adults, we re-analyzed two single-cell TCRseq+gene expression datasets, one studied (39) the multisystem inflammatory syndrome in children (MIS-C) associated with pediatric COVID-19 (pcovid) and the other studied adult COVID-19 (acovid) with different severity (40). While the acovid dataset consists of experimentally isolated SARS-CoV2-reactive CD8 T-cells from PBMCs, the pcovid dataset contains non-enriched CD8 T-cells from PBMCs. We used TCRanno’s qualitative annotations to isolate T-cells likely specific to SARS-CoV2 in the pcovid dataset. Then we identified differentially expressed genes and enriched biological processes (41) associated with pcovid versus MIS-C from the TCRanno-isolated T-cells and mild acovid versus severe acovid from the experimentally-isolated T-cells, respectively. Consistent with previous findings (39), the predicted SARS-CoV2-reactive CD8 T-cells in MIS-C resembled a toxic shock syndrome (Supplementary Figure S16, top panel), manifested with significantly increased respiration (52), cell death, mRNA degradation and strong inflammatory responses via Wnt/*β*-catenin-TCF signaling pathway (53, 54), which produce pro-inflammatory cytokines such as granulocyte-macrophage colony-stimulating factor and tumor necrosis factor. A similar profile was observed in those of the severe acovid patients, with a significant increase in glycolysis, apoptosis and inflammation (Supplementary Figure S17, top panel). We also found that the predicted SARS-CoV2-specific CD8 T-cells in pcovid resembled those in the mild acovid by showing a regulated viral immune response, as evidenced by proper T-cell activation via antigen presentation and intact type I interferon signaling (Supplementary Figure S16-17, bottom panels). Our results demonstrated that the TCRanno-isolated pcovid T-cells were transcriptionally similar to the experiment-isolated acovid T-cells.

## Discussion

We presented TCRanno, a toolkit for qualitative and quantitative annotations of TCR repertoire specificity. TCRanno gives annotations at epitope, antigen and organism levels based on validated knowledge or predictions according to the extent of mapping (CM, PM) to database TCR sequences. We verified TCRanno’s performance on specificity prediction with four independent datasets and demonstrated advantages in performance and speed over state-of-the-art approaches. We then applied TCRanno to analyses of bulk TCRseq or single-cell TCRseq+gene expression data of peripheral TCR repertoires under five different conditions (healthy, pre-/post-vaccination, SARS-CoV2 exposed/infected, SLE patients and cancers) and showed how TCRanno helps discover meaningful insights.

TCRanno has demonstrated improvements in methodology for specificity prediction. TCRanno’s prediction algorithm is an effective combination of machine-learning and non-machine-learning methods with self-adapted parameter tuning (see Methods). We cannot guarantee a TCR’s cognate epitope exists in the reference database. The number of unknown epitope classes outweighs the number of known epitope classes. Therefore, instead of training a classification model to limited classes, we focused on generating epitope-aware, discriminative embeddings with generalizability to unseen, unlimited classes. Conventional variational autoencoders with the classic, unsupervised encoder-decoder structure can generate similar embeddings for similar TCR sequences (55). We designed an encoder-classifier structure that provides supervised epitope information to the encoder during training, so that similar embeddings now represent TCR sequences likely of the same epitope specificity. Besides performance and speed, ease of adaptability to future data is another important advantage of TCRanno. Since TCRanno’s specificity prediction is not a classification task, we expect the algorithm to work on TCR sequences specific to previously unseen epitopes as long as new reference data of these epitopes are added to the database. Unlike traditional classification models which require reconstruction and training of a new model to fit a new epitope class, the adaptability cost for TCRanno (i.e., updating the database) is ignorable. Alternatively, the current pre-trained encoder model can be retrained when novel large-scale labeled data becomes available or even be replaced with customized, more efficient model structures.

TCRanno estimates a T-cell repertoire’s specificity composition, i.e. the respective fractions specific to different epitopes, antigens and organisms. The specificity composition reflects the past or current immune responses and therefore may vary upon conditions. For example, we observed a significant increase in the estimated fractions of TCRs specific to CMV and its major antigens within CMV^+^ individuals, and a significant increase in the estimated SARS-CoV2 specific TCR fraction within SARS-CoV2 exposed/infected individuals compared to healthy controls. Note that a high TCR fraction does not necessarily mean a physically large number of cells, as shown in previous studies that SARS-CoV2 infected individuals have a higher fraction of SARS-CoV2-specific CD8 T-cells but a lower number of SARS-CoV2-specific CD8 T-cells probably due to lymphopenia (1). Comparative analyses on specificity composition help us to understand how a condition or event (infection, disease, vaccination, etc.) changes the landscape of a T-cell pool and identify biomarkers associated with a particular condition.

Some interesting findings were note-worthy. Consistent with previous observations (19), we found an appreciable amount of known and predicted CMV-specific TCRs in CMV^−^ individuals. Moreover, we found a large overlap of CMV-specific TCRs between CMV^+^ and CMV^−^ individuals which constituted a dominating fraction in both sub-populations. Considering the extremely high CMV prevalence worldwide, our results suggested that effective defense against the virus instead of non-exposure could account for the seronegativity in at least some CMV^−^ individuals. Indeed, an early study (56) revealed that CMV mRNA transcripts were detected in 44% of CMV seronegative healthy individuals, though the underlying mechanisms for CMV abortive infection in healthy individuals remained to be elucidate. Perspective studies comparing the presence and functions of CMV-specific T-cells pre- and post-infection in CMV^+^ individuals, and that between the two sub-populations, are awaited to elucidate why CMV^+^ individuals fail to clear the virus despite sharing abundant CMV-specific TCRs with CMV^−^ individuals.

We identified a substantial amount of TCRs likely reactive to SARS-CoV2 in the pre-pandemic population, in line with recent evidence demonstrating *in vitro* cross-reactivity of SARS-CoV2-specific T-cells from pre-pandemic samples against antigens of commensal bacteria (45). Together with our observations: i) the estimated SARS-CoV2 specific TCR fraction increased over age; ii) the deceased sub-population possessed significantly higher SARS-CoV2-reactive and self-reactive TCR fractions than the convalescents, the results suggested that the aggressive and systemic symptoms of COVID-19 may come from the cross-reactivity by a broad spectrum of pre-existing memory T-cells. The exposure to SARS-CoV2 can result in a high-magnitude immune response that causes systemic inflammation; in severe cases, excessive immunity may further overwhelm the tolerance mechanisms, attacking self-antigens or non-harmful microbiota present in multiple organs even after the clearance of the virus.

Our quantitative annotations in healthy versus SLE repertoires identified HBV antigens as a potential trigger of SLE onset. Interestingly, hepatitis B vaccination has been previously reported to cause or accelerate SLE onset in mouse models and human participants receiving the vaccine (57–59). Identifying and characterizing the autoimmunity-inducing HBV antigens and removing them from the vaccine could be a possible solution. Our results suggested the precore/core protein and the DNA polymerase as possible autoimmunity-inducing antigens. Moreover, our observations indicated an unusually high memory-like fraction in SLE compared to all other populations, and we further demonstrated the enlarged memory pool was likely due to non-specific, uniform clonal expansions via bystander T-cell activation. Mechanisms behind autoimmunity-associated bystander activation and future prospects for therapeutic development have been discussed elsewhere in the literature (60).

We also demonstrated the feasibility of using the quantitative differences in TCR repertoire specificity composition to distinguish repertoires of healthy versus pathological conditions including viral infections, autoimmunity and cancers (especially melanoma and lung cancer). Features extracted from TCRanno’s quantitative annotations can help develop novel non-invasive methods for disease screening and monitoring. More importantly, all these results revealed that the peripheral T-cell memory pool is dominated by TCRs recognizing common infectious agents (SARS-CoV2, CMV, Influenza, EBV, HBV), and in some cancer types the fraction could be even higher, suggesting possible cross-reactivity of these viral-specific TCRs against tumor antigens. Unlike the small number of tumor-specific TCRs, these large fractions of pre-existing TCRs may serve as an attractive weapon that can be retargeted toward cancer. Our results provided supporting evidence for the plausibility of using infectious disease vaccines in the treatment of cancers (49–51, 61).

This study has several limitations. First, the distribution of TCR sequences within the reference database affects the computed repertoire specificity composition. Extensive studies on T-cell responses to SARS-CoV2 have greatly enriched our knowledge of TCR:peptide-MHC recognition and enlarged the size of the reference database, while inevitably introducing huge differences in the magnitude of sample sizes for different epitopes, antigens and organisms. Therefore, the fraction (percentage) composition should be anticipated with caution that the non-SARS-CoV2 counterparts may be underestimated and hard to compare directly. Nevertheless, the CM fraction can at least serve as an independent lower bound as to infer the amount of TCRs specific to the epitope/antigen/organism of interest. Accumulating more non-SARS-CoV2 data (e.g., CMV, EBV, Influenza) will help solve the data-imbalance problem. Second, the total size of known data (about 200k) is still too small, compared to the expected space of 107 unique TCR sequences per individual (19). This resulted in only 7% of the effective fraction being completely matched to the reference database, affecting the confidence of annotations. Similarly, the generation and accumulation of more verified data will increase the size of the reference database and improve the fraction of completely matched input sequences. Third, compared to viral antigens, current knowledge of self or tumor antigens (neoantigens) and their specific TCRs remained poor, affecting annotations for cancer and autoimmune diseases. Finally, TCRanno currently only considers MHC-I restricted (i.e., CD8) TCR***β*** CDR3 sequences, due to even fewer verified and labeled CD4/TCRa/CDR1/CDR2 data for model training.

## Data and Software Availability

All the data used within this study can be retrieved from public databases. TCRanno is an open-source Python package with source code freely available at https://github.com/deepomicslab/TCRanno. A simplified web implementation is available under the TIMEDB (62) platform: https://timedb.deepomics.org/submit/tcrAnalyses.

## Acknowledgements

Not applicable.

## References

1. Joshua M Francis, Del Leistritz-Edwards, Augustine Dunn, Christina Tarr, Jesse Lehman, Conor Dempsey, Andrew Hamel, Violeta Rayon, Gang Liu, Yuntong Wang, Marcos Wille, Melissa Durkin, Kane Hadley, Aswathy Sheena, Benjamin Roscoe, Mark Ng, Graham Rockwell, Margaret Manto, Elizabeth Gienger, Joshua Nickerson, Amir Moarefi, Michael Noble, Thomas Malia, Philip D Bardwell, William Gordon, Joanna Swain, Mojca Skoberne, Karsten Sauer, Tim Harris, Ananda W Goldrath, Alex K Shalek, Anthony J Coyle, Christophe Benoist, and Daniel C Pregibon. Allelic variation in class I HLA determines CD8 + T cell repertoire shape and cross-reactive memory responses to SARS-CoV-2 MGH COVID-19 Collection and Processing Team. Sci. Immunol, 2022.

2. 10x Genomics. A new way of exploring immunity–linking highly multiplexed antigen recognition to immune repertoire and phenotype. Tech. rep, 2019.

3. Theres Oakes, James M. Heather, Katharine Best, Rachel Byng-Maddick, Connor Husovsky, Mazlina Ismail, Kroopa Joshi, Gavin Maxwell, Mahdad Noursadeghi, Natalie Riddell, Tabea Ruehl, Carolin T. Turner, Imran Uddin, and Benny Chain. Quantitative characterization of the T cell receptor repertoire of naïve and memory subsets using an integrated experimental and computational pipeline which is robust, economical, and versatile. Frontiers in Immunology, 2017.

4. Jacob Glanville, Huang Huang, Allison Nau, Olivia Hatton, Lisa E. Wagar, Florian Rubelt, Xuhuai Ji, Arnold Han, Sheri M. Krams, Christina Pettus, Nikhil Haas, Cecilia S.Lindestam Arlehamn, Alessandro Sette, Scott D. Boyd, Thomas J. Scriba, Olivia M. Martinez, and Mark M. Davis. Identifying specificity groups in the T cell receptor repertoire. Nature, 2017.

5. Huang Huang, Chunlin Wang, Florian Rubelt, Thomas J. Scriba, and Mark M. Davis. Analyzing the Mycobacterium tuberculosis immune response by T-cell receptor clustering with GLIPH2 and genome-wide antigen screening. Nature Biotechnology 2020 38:10, 2020.

6. William D. Chronister, Austin Crinklaw, Swapnil Mahajan, Randi Vita, Zeynep Koşaloğ lu-Yalçın, Zhen Yan, Jason A. Greenbaum, Leon E. Jessen, Morten Nielsen, Scott Christley, Lindsay G. Cowell, Alessandro Sette, and Bjoern Peters. TCRMatch: Predicting T-Cell Receptor Specificity Based on Sequence Similarity to Previously Characterized Receptors. Frontiers in Immunology, 2021.

7. Koshlan Mayer-Blackwell, Stefan Schattgen, Liel Cohen-Lavi, Jeremy C Crawford, Aisha Souquette, Jessica A Gaevert, Tomer Hertz, Paul G Thomas, Philip Bradley, and Andrew Fiore-Gartland. Tcr meta-clonotypes for biomarker discovery with tcrdist3 enabled identification of public, hla-restricted clusters of sars-cov-2 tcrs. Elife, 10:e68605, 2021.

8. Sofie Gielis, Pieter Moris, Wout Bittremieux, Nicolas De Neuter, Benson Ogunjimi, Kris Laukens, and Pieter Meysman. Detection of enriched t cell epitope specificity in full t cell receptor sequence repertoires. Frontiers in immunology, 10:2820, 2019.

9. Alessandro Montemurro, Viktoria Schuster, Helle Rus Povlsen, Amalie Kai Bentzen, Vanessa Jurtz, William D Chronister, Austin Crinklaw, Sine R Hadrup, Ole Winther, Bjoern Peters, et al. Nettcr-2.0 enables accurate prediction of tcr-peptide binding by using paired tcr *α* and *β* sequence data. Communications biology, 4(1):1–13, 2021.

10. Jeff Daily. Parasail: SIMD C library for global, semi-global, and local pairwise sequence alignments. BMC Bioinformatics, 2016.

11. Yandong Wen, Kaipeng Zhang, Zhifeng Li, and Yu Qiao. A discriminative feature learning approach for deep face recognition. In European conference on computer vision, pages 499–515. Springer, 2016.

12. Randi Vita, Swapnil Mahajan, James A. Overton, Sandeep Kumar Dhanda, Sheridan Martini, Jason R. Cantrell, Daniel K. Wheeler, Alessandro Sette, and Bjoern Peters. The Immune Epitope Database (IEDB): 2018 update. Nucleic acids research, 2019.

13. Dmitry V. Bagaev, Renske M.A. Vroomans, Jerome Samir, Ulrik Stervbo, Cristina Rius, Garry Dolton, Alexander Greenshields-Watson, Meriem Attaf, Evgeny S. Egorov, Ivan V. Zvyagin, Nina Babel, David K. Cole, Andrew J. Godkin, Andrew K. Sewell, Can Kesmir, Dmitriy M. Chudakov, Fabio Luciani, and Mikhail Shugay. VDJdb in 2019: database extension, new analysis infrastructure and a T-cell receptor motif compendium. Nucleic Acids Research, 2020.

14. Nili Tickotsky, Tal Sagiv, Jaime Prilusky, Eric Shifrut, and Nir Friedman. McPAS-TCR: a manually curated catalogue of pathology-associated T cell receptor sequences. Bioinformatics (Oxford, England), 2017.

15. Sean Nolan, Marissa Vignali, Mark Klinger, Jennifer N Dines, Ian M Kaplan, Emily Svejnoha, Tracy Craft, Katie Boland, Mitch Pesesky, Rachel M Gittelman, Thomas M Snyder, Christopher J Gooley, Simona Semprini, Claudio Cerchione, Massimiliano Mazza, Ottavia M Delmonte, Kerry Dobbs, Gonzalo Carreño-Tarragona, Santiago Barrio, Vittorio Sambri, Giovanni Martinelli, Jason D Goldman, James R Heath, Luigi D Notarangelo, Jonathan M Carlson, Joaquin Martinez-Lopez, and Harlan S Robins. A large-scale database of T-cell receptor beta (TCR *β*) sequences and binding associations from natural and synthetic exposure to SARS-CoV-2. Research square, 2020.

16. Anastasia A. Minervina, Mikhail V. Pogorelyy, Ekaterina A. Komech, Vadim K. Karnaukhov, Petra Bacher, Elisa Rosati, Andre Franke, Dmitriy M. Chudakov, Ilgar Z. Mamedov, Yury B. Lebedev, Thierry Mora, and Aleksandra M. Walczak. Primary and secondary anti-viral response captured by the dynamics and phenotype of individual T cell clones. eLife, 2020.

17. In Young Song, Anna Gil, Rabinarayan Mishra, Dario Ghersi, Liisa K. Selin, and Lawrence J. Stern. Broad TCR repertoire and diverse structural solutions for recognition of an immunodominant CD8+ T cell epitope. Nature Structural and Molecular Biology 2017 24:4, 2017.

18. Anna Gil, Larisa Kamga, Ramakanth Chirravuri-Venkata, Nuray Aslan, Fransenio Clark, Dario Ghersi, Katherine Luzuriaga, and Liisa K. Selin. Epstein-Barr Virus Epitope–Major Histocompatibility Complex Interaction Combined with Convergent Recombination Drives Selection of Diverse T Cell Receptor *α* and *β* Repertoires. mBio, 2020.

19. Ryan O. Emerson, William S. DeWitt, Marissa Vignali, Jenna Gravley, Joyce K. Hu, Edward J. Osborne, Cindy Desmarais, Mark Klinger, Christopher S. Carlson, John A. Hansen, Mark Rieder, and Harlan S. Robins. Immunosequencing identifies signatures of cytomegalovirus exposure history and HLA-mediated effects on the T cell repertoire. Nature Genetics 2017 49:5, 2017.

20. Phillip A. Swanson, Marcelino Padilla, Wesley Hoyland, Kelly McGlinchey, Paul A. Fields, Sagida Bibi, Saul N. Faust, Adrian B. McDermott, Teresa Lambe, Andrew J. Pollard, Nicholas M. Durham, and Elizabeth J. Kelly. AZD1222/ChAdOx1 nCoV-19 vaccination induces a polyfunctional spike protein-specific TH1 response with a diverse TCR repertoire. Science Translational Medicine, 2021.

21. Xiao Liu, Wei Zhang, Ming Zhao, Longfei Fu, Limin Liu, Jinghua Wu, Shuangyan Luo, Longlong Wang, Zijun Wang, Liya Lin, Yan Liu, Shiyu Wang, Yang Yang, Lihua Luo, Juqing Jiang, Xie Wang, Yixin Tan, Tao Li, Bochen Zhu, Yi Zhao, Xiaofei Gao, Ziyun Wan, Cancan Huang, Mingyan Fang, Qianwen Li, Huanhuan Peng, Xiangping Liao, Jinwei Chen, Fen Li, Guanghui Ling, Hongjun Zhao, Hui Luo, Zhongyuan Xiang, Jieyue Liao, Yu Liu, Heng Yin, Hai Long, Haijing Wu, Huanming Yang, Jian Wang, and Qianjin Lu. T cell receptor *β* repertoires as novel diagnostic markers for systemic lupus erythematosus and rheumatoid arthritis. Annals of the rheumatic diseases, 2019.

22. Erik Yusko, Marissa Vignali, Richard K Wilson, Elaine R Mardis, F Stephen Hodi, Christine Horak, Han Chang, David M Woods, Harlan Robins, and Jeffrey Weber. Association of tumor microenvironment t-cell repertoire and mutational load with clinical outcome after sequential checkpoint blockade in melanomatumor microenvironment and t-cell biomarkers in melanoma. Cancer immunology research, 7(3):458–465, 2019.

23. Sara Valpione, Elena Galvani, Joshua Tweedy, Piyushkumar A Mundra, Antonia Banyard, Philippa Middlehurst, Jeff Barry, Sarah Mills, Zena Salih, John Weightman, et al. Immune awakening revealed by peripheral t cell dynamics after one cycle of immunotherapy. Nature cancer, 1(2):210–221, 2020.

24. Lidia Robert, Jennifer Tsoi, Xiaoyan Wang, Ryan Emerson, Blanca Homet, Thinle Chodon, Stephen Mok, Rong Rong Huang, Alistair J Cochran, Begoña Comin-Anduix, et al. Ctla4 blockade broadens the peripheral t-cell receptor repertoirectla4 blockade broadens the peripheral tcr repertoire. Clinical Cancer Research, 20(9):2424–2432, 2014.

25. Jani Huuhtanen, Liang Chen, Emmi Jokinen, Henna Kasanen, Tapio Lönnberg, Anna Kreutzman, Katriina Peltola, Micaela Hernberg, Chunlin Wang, Cassian Yee, et al. Evolution and modulation of antigen-specific t cell responses in melanoma patients. Nature communications, 13(1):1–14, 2022.

26. Silvia C Formenti, Nils-Petter Rudqvist, Encouse Golden, Benjamin Cooper, Erik Wennerberg, Claire Lhuillier, Claire Vanpouille-Box, Kent Friedman, Lucas Ferrari de Andrade, Kai W Wucherpfennig, et al. Radiotherapy induces responses of lung cancer to ctla-4 blockade. Nature medicine, 24(12):1845–1851, 2018.

27. Alexandre Reuben, Jiexin Zhang, Shin-Heng Chiou, Rachel M Gittelman, Jun Li, Won-Chul Lee, Junya Fujimoto, Carmen Behrens, Xiaoke Liu, Feng Wang, et al. Comprehensive t cell repertoire characterization of non-small cell lung cancer. Nature communications, 11(1): 1–13, 2020.

28. Melody S Hsu, Shaina Sedighim, Tina Wang, Joseph P Antonios, Richard G Everson, Alexander M Tucker, Lin Du, Ryan Emerson, Erik Yusko, Catherine Sanders, et al. Tcr sequencing can identify and track glioma-infiltrating t cells after dc vaccinationtcr usage in glioblastoma patients before and after dc vaccination. Cancer immunology research, 4(5): 412–418, 2016.

29. Dung T Le, Jennifer N Durham, Kellie N Smith, Hao Wang, Bjarne R Bartlett, Laveet K Aulakh, Steve Lu, Holly Kemberling, Cara Wilt, Brandon S Luber, et al. Mismatch repair deficiency predicts response of solid tumors to pd-1 blockade. Science, 357(6349):409–413, 2017.

30. Ryan O Emerson, Anna M Sherwood, Mark J Rieder, Jamie Guenthoer, David W Williamson, Christopher S Carlson, Charles W Drescher, Muneesh Tewari, Jason H Bielas, and Harlan S Robins. High-throughput sequencing of t-cell receptors reveals a homogeneous repertoire of tumour-infiltrating lymphocytes in ovarian cancer. The Journal of pathology, 231(4):433–440, 2013.

31. Alexandra Snyder, Tavi Nathanson, Samuel A Funt, Arun Ahuja, Jacqueline Buros Novik, Matthew D Hellmann, Eliza Chang, Bulent Arman Aksoy, Hikmat Al-Ahmadie, Erik Yusko, et al. Contribution of systemic and somatic factors to clinical response and resistance to pd-l1 blockade in urothelial cancer: an exploratory multi-omic analysis. PLoS medicine, 14 (5):e1002309, 2017.

32. David E Hamm. immunoseq hstcrb-v4b control data. immuneACCESS, 10:21417, 2020.

33. David B Page, Jianda Yuan, David Redmond, Y Hanna Wen, Jeremy C Durack, Ryan Emerson, Stephen Solomon, Zhiwan Dong, Phillip Wong, Christopher Comstock, et al. Deep sequencing of t-cell receptor dna as a biomarker of clonally expanded tils in breast cancer after immunotherapyt-cell receptor dna deep sequencing in breast cancer. Cancer immunology research, 4(10):835–844, 2016.

34. Jeffrey J Wallin, Johanna C Bendell, Roel Funke, Mario Sznol, Konstanty Korski, Suzanne Jones, Genevive Hernandez, James Mier, Xian He, F Stephen Hodi, et al. Atezolizumab in combination with bevacizumab enhances antigen-specific t-cell migration in metastatic renal cell carcinoma. Nature communications, 7(1):1–8, 2016.

35. Kathryn E Yost, Ansuman T Satpathy, Daniel K Wells, Yanyan Qi, Chunlin Wang, Robin Kageyama, Katherine L McNamara, Jeffrey M Granja, Kavita Y Sarin, Ryanne A Brown, et al. Clonal replacement of tumor-specific t cells following pd-1 blockade. Nature medicine, 25(8):1251–1259, 2019.

36. Jared Dean, Ryan O Emerson, Marissa Vignali, Anna M Sherwood, Mark J Rieder, Christopher S Carlson, and Harlan S Robins. Annotation of pseudogenic gene segments by massively parallel sequencing of rearranged lymphocyte receptor loci. Genome medicine, 7(1):1–8, 2015.

37. L Chen and SC Li. Incorporating cell hierarchy to decipher the functional diversity of single cells. Nucleic Acids Research, pages gkac1044–gkac1044, 2022.

38. Leland McInnes, John Healy, and James Melville. Umap: Uniform manifold approximation and projection for dimension reduction. arXiv preprint arXiv:1802.03426, 2018.

39. Keith Sacco, Riccardo Castagnoli, Svetlana Vakkilainen, Can Liu, Ottavia M Delmonte, Cihan Oguz, Ian M Kaplan, Sara Alehashemi, Peter D Burbelo, Farzana Bhuyan, et al. Immunopathological signatures in multisystem inflammatory syndrome in children and pediatric covid-19. Nature Medicine, 28(5):1050–1062, 2022.

40. Anthony Kusnadi, Ciro Ramírez-Suástegui, Vicente Fajardo, Serena J Chee, Benjamin J Meckiff, Hayley Simon, Emanuela Pelosi, Grégory Seumois, Ferhat Ay, Pandurangan Vijayanand, et al. Severely ill patients with covid-19 display impaired exhaustion features in sars-cov-2–reactive cd8+ t cells. Science immunology, 6(55):eabe4782, 2021.

41. Yuhan Hao, Stephanie Hao, Erica Andersen-Nissen, William M Mauck III, Shiwei Zheng, Andrew Butler, Maddie J Lee, Aaron J Wilk, Charlotte Darby, Michael Zager, et al. Integrated analysis of multimodal single-cell data. Cell, 184(13):3573–3587, 2021.

42. Jett Crowdis, Meng Xiao He, Brendan Reardon, and Eliezer M Van Allen. Comut: visualizing integrated molecular information with comutation plots. Bioinformatics, 36(15):4348–4349, 2020.

43. Donna L Farber, Naomi A Yudanin, and Nicholas P Restifo. Human memory t cells: generation, compartmentalization and homeostasis. Nature Reviews Immunology, 14(1): 24–35, 2014.

44. Leo Swadling, Mariana O Diniz, Nathalie M Schmidt, Oliver E Amin, Aneesh Chandran, Emily Shaw, Corinna Pade, Joseph M Gibbons, Nina Le Bert, Anthony T Tan, et al. Pre-existing polymerase-specific t cells expand in abortive seronegative sars-cov-2. Nature, 601 (7891):110–117, 2022.

45. Laurent Bartolo, Sumbul Afroz, Yi-Gen Pan, Ruozhang Xu, Lea Williams, Chin-Fang Lin, Ceylan Tanes, Kyle Bittinger, Elliot S. Friedman, Phyllis A. Gimotty, Gary D. Wu, and Laura F. Su. SARS-CoV-2-specific T cells in unexposed adults display broad trafficking potential and cross-react with commensal antigens. Science Immunology, 2022.

46. Omer Gendelman, Naim Mahroum, Doron Comaneshter, Pnina Rotman-Pikielny, Arnon D. Cohen, Howard Amital, and Michael Sherf. Hepatitis B carrier state among SLE patients: case–control study. Immunologic Research 2016 65:1, 2016.

47. Ming Han Chen, Chien Sheng Wu, Ming Huang Chen, Chang Youh Tsai, Fa Yauh Lee, and Yi Hsiang Huang. High risk of viral reactivation in hepatitis b patients with systemic lupus erythematosus. International Journal of Molecular Sciences, 2021.

48. Paul L Crispen and Sergei Kusmartsev. Mechanisms of immune evasion in bladder cancer. Cancer Immunology, Immunotherapy, 69(1):3–14, 2020.

49. Uzoma K Iheagwara, Pamela L Beatty, Phu T Van, Ted M Ross, Jonathan S Minden, and Olivera J Finn. Influenza virus infection elicits protective antibodies and t cells specific for host cell antigens also expressed as tumor-associated antigens: A new view of cancer immunosurveillanceviral infections elicit immunity against tumor antigens. Cancer immunology research, 2(3):263–273, 2014.

50. Jenna H Newman, C Brent Chesson, Nora L Herzog, Praveen K Bommareddy, Salvatore M Aspromonte, Russell Pepe, Ricardo Estupinian, Mones M Aboelatta, Stuti Buddhadev, Saeed Tarabichi, et al. Intratumoral injection of the seasonal flu shot converts immunologically cold tumors to hot and serves as an immunotherapy for cancer. Proceedings of the National Academy of Sciences, 117(2):1119–1128, 2020.

51. Eileena F Giurini, Michael Williams, Adam Morin, Andrew Zloza, and Kajal H Gupta. Inactivated sars-cov-2 reprograms the tumor immune microenvironment and improves murine cancer outcomes. bioRxiv, 2022.

52. Adalberto Fernandes Santos, Pedro Póvoa, Paulo Paixão, António Mendonça, and Luís Taborda-Barata. Changes in glycolytic pathway in sars-cov 2 infection and their importance in understanding the severity of covid-19. Frontiers in chemistry, 9:685196, 2021.

53. Alexandre Vallee, Yves Lecarpentier, and Jean-Noel Vallee. Interplay of opposing effects of the wnt/*β*-catenin pathway and ppar *γ* and implications for sars-cov2 treatment. Frontiers in immunology, 12:666693, 2021.

54. Johanna K Ljungberg, Jessica C Kling, Thao Thanh Tran, and Antje Blumenthal. Functions of the wnt signaling network in shaping host responses to infection. Frontiers in Immunology, 10:2521, 2019.

55. Davidsen Kristian, Branden J Olson, William S DeWitt III, Jean Feng, Elias Harkins, Philip Bradley, and Frederick A Matsen IV. Deep generative models for t cell receptor protein sequences. eLife, 8, 2019.

56. Lin J Zhang, Philip Hanff, Cynthia Rutherford, WH Churchill, and Clyde S Crumpacker. Detection of human cytomegalovirus dna, rna, and antibody in normal donor blood. Journal of Infectious Diseases, 171(4):1002–1006, 1995.

57. N Agmon-Levin, Y Zafrir, Z Paz, T Shilton, G Zandman-Goddard, and Y Shoenfeld. Ten cases of systemic lupus erythematosus related to hepatitis b vaccine. Lupus, 18(13):1192–1197, 2009.

58. Y Zafrir, N Agmon-Levin, Z Paz, T Shilton, and Y Shoenfeld. Autoimmunity following hepatitis b vaccine as part of the spectrum of ‘autoimmune (auto-inflammatory) syndrome induced by adjuvants’(asia): analysis of 93 cases. Lupus, 21(2):146–152, 2012.

59. Nancy Agmon-Levin, María-Teresa Arango, Shaye Kivity, Aviva Katzav, Boris Gilburd, Miri Blank, Nir Tomer, Alex Volkov, Iris Barshack, Joab Chapman, et al. Immunization with hepatitis b vaccine accelerates sle-like disease in a murine model. Journal of autoimmunity, 54:21–32, 2014.

60. Yovana Pacheco, Yeny Acosta-Ampudia, Diana M Monsalve, Christopher Chang, M Eric Gershwin, and Juan-Manuel Anaya. Bystander activation and autoimmunity. Journal of autoimmunity, 103:102301, 2019.

61. Liese Vandeborne, Pan Pantziarka, An MT Van Nuffel, and Gauthier Bouche. Repurposing infectious diseases vaccines against cancer. Frontiers in Oncology, page 1767, 2021.

62. X Wang, L Chen, W Liu, Y Zhang, D Liu, C Zhou, S Shi, J Dong, Z Lai, B Zhao, W Zhang, H Cheng, and S Li. Timedb: tumor immune micro-environment cell composition database with automatic analysis and interactive visualization. Nucleic Acids Research, pages gkac1006–gkac1006, 2022.

