## Supplementary Materials for "Quantitative Annotations of T-Cell Repertoire Specificity"

# -

### Supplementary Materials

Jiaqi Luo, Xueying Wang, Yiping Zou, Lingxi Chen, Wei Liu, Wei Zhang,

and Shuai Cheng Li*

S2. Mean fraction (percentage) composition of TCR repertoire specificity of CMV+ individuals.. 22

S3. Mean fraction (percentage) composition of TCR repertoire specificity of CMV- individuals... 23

S4. Mean fraction (percentage) composition of TCR repertoire specificity of healthy individuals. 24
S5. Mean fraction (percentage) composition of TCR repertoire specificity of SARS-CoV2 exposed/infected individuals. 25
S6. Mean fraction (percentage) composition of TCR repertoire specificity of COVID-19 deceased individuals. 26
S7. Mean fraction (percentage) composition of TCR repertoire specificity of COVID-19 convalescents. 27
S8. Mean fraction (percentage) composition of TCR repertoire specificity of COVID-19 AZD1222 pre-vaccination individuals. 28
S9. Mean fraction (percentage) composition of TCR repertoire specificity of COVID-19 AZD1222 post-vaccination individuals. 29
S10. Mean fraction (percentage) composition of TCR repertoire specificity of SLE patients. 30
S11. Mean fraction (percentage) composition of TCR repertoire specificity of melanoma patients. 31
S12. Mean fraction (percentage) composition of TCR repertoire specificity of lung cancer patients. 32
S13. Mean fraction (percentage) composition of TCR repertoire specificity of breast cancer patients. 33
S14. Mean fraction (percentage) composition of TCR repertoire specificity of urothelial bladder cancer patients. 34
S15. Mean fraction (percentage) composition of TCR repertoire specificity of all cancer patients. 35
S16. ANOVA test p-values for the 7 numeric features on different comparisons of sub-populations. 36

### Supplementary Methods

#### TCR specificity prediction

We developed an ensemble similarity measure called tcr2tcr score, which is a self-tuned weighted sum of three similarity scores $s_T$, $s_D$ and $s_E$, to measure the similarity of the query TCR sequence to each of the reference TCR sequences. Here we use $\tau_q$ to denote a query TCR sequence, $| \tau_q |$ to denote the length of the query TCR sequence and $\tau_i$ ($i \in R$) to denote a reference TCR sequence.

The first similarity score $s_T$ is a real number measuring the similarity of the trimmed query and reference TCR sequences by Levenshtein distance. Trimming of heads and tails of the TCR sequences is performed before Levenshtein distance calculation to focus on centre positions. Depending on the length of the query TCR sequence, the number of amino acids trimmed on both ends (T) is determined by the formula specified below.

$$\mathrm{T} = \left\lceil \frac{|\tau_q| - 10}{4} \right\rceil$$

Then we calculate the Levenshtein distance of the trimmed query TCR sequence $\tau_q^T$ toward each of all the trimmed TCR sequences $\tau_i^T$, denoted as $d_{T_{q,i}}$ .

$$d_{T_{q,i}} = \mathrm{lev}(\tau_q^T, \tau_i^T), \ \mathrm{i} \in \mathrm{R}$$

The pairwise score $s_{T_{q,i}}$ is the scaled form of $d_{T_{q,i}}$ so that an $s_{T_{q,i}}$ score of 1 map to zero Levenshtein distance (i.e., identical sequences) while a small score map to distinct sequences. The score can even be negative when the reference TCR sequence is much longer than the query TCR sequence.

$$s_{T_{q,i}} = 1 - \frac{d_{T_{q,i}}}{|\tau_q|} = 1 - \frac{\mathrm{lev}(\tau_q^T, \tau_i^T)}{|\tau_q|}, \ \mathrm{i} \in \mathrm{R}$$

Here the denominator is the length of the query instead of the maximum $d_T$ value to allow calculations independent of the reference database.

As shown in Supplementary Figure S2, restricting the selection to a smaller pool of reference TCR sequences with the top n=100 $s_{T_{q,i}}$ scores to the query increases the prediction performance and reduces the runtime. Therefore, after calculating $s_{T_{q,i}}$ for all reference TCR sequences, the top-100 pool (denoted as $\mathrm{R}_{pool}$) will be selected for downstream calculations.

The second similarity score $s_D$ is a real number measuring the similarity of D segments within the TCR sequences by Levenshtein distance. A TCR CDR3 amino acid (aa) sequence consists of V, D, and J segments. The V and J segments were obtained by mapping the TCR CDR3 aa sequence to a map of known human V/J segments. The known V/J map was generated by first mapping more than 20 million TCR DNA sequences extracted from 587 healthy donors (1) to the IMGT (2) V and J gene databases, then the mapped V and J segments of DNA sequence were translated back to aa sequences. The D segment (aa sequence) can then be obtained by subtracting the V and J segments from the full TCR CDR3 aa sequence. The pairwise distance $d_{D_{q,i}}$ and score $s_{D_{q,i}}$ are specified as follow.

$$d_{D_{q,i}} = \mathrm{lev}(\tau_q^D, \tau_i^D), \ \mathrm{i} \in \mathrm{R}_{pool}$$

$$s_{D_{q,i}} = 1 - \frac{\mathrm{lev}(\tau_q^D, \tau_i^D)}{max\{d_{D_q}\}} = 1 - \frac{\mathrm{lev}(\tau_q^D, \tau_i^D)}{\max_{j \in R_{pool}} \{\mathrm{lev}(\tau_q^D, \tau_j^D)\}}$$

Similar to $s_{T_{q,i}}$, the $s_{D_{q,i}}$ score is obtained by scaling $d_{D_{q,i}}$. Since the D segment length can be zero, instead of dividing the query TCR's D segment length, here we use the maximum of all $d_{D_{q,i}}$ values as the denominator, and $s_{D_{q,i}}$ has the range of 0-1.

The third similarity score $s_E$ is the cosine similarity (ranged -1 to 1) of the embedding vectors generated by the encoder (denoted as ε). The embedding vectors ($v_q$, $v_i$) representing the query TCR sequence $\tau_q$ and the reference TCR sequence $\tau_i$ is:

$$v_q = \epsilon(\tau_q), v_i = \epsilon(\tau_i),$$

and the pairwise score $S_{E_{q,i}}$ is the cosine similarity of the two vectors:

$$S_{E_{q,i}} = \cos\langle v_q, v_i \rangle = \cos\langle \epsilon(\tau_q), \epsilon(\tau_i) \rangle.$$

Finally, the tcr2tcr score $S_{q,i}$ is the weighted sum of the above three similarity scores. The weights of the three similarity scores ($w_T$, $w_D$, $w_E$) are their reversed proportion of variances:

$$\sigma_T^2 = \frac{1}{n} \sum_{i=1}^{n} (x_i - \mu_{s_T})^2, x_i \in s_T$$

$$\sigma_D^2 = \frac{1}{n} \sum_{i=1}^{n} (x_i - \mu_{s_D})^2, x_i \in s_D$$

$$\sigma_E^2 = \frac{1}{n} \sum_{i=1}^{n} (x_i - \mu_{s_E})^2, x_i \in s_E$$

$$S_{var} = \sigma_T^2 + \sigma_D^2 + \sigma_E^2$$

$$w_T = \frac{S_{var}}{\sigma_T^2}, w_D = \frac{S_{var}}{\sigma_D^2}, w_E = \frac{S_{var}}{\sigma_E^2}$$

where $\sigma_T^2$, $\mu_{s_T}$ is the variance and mean of all $S_{T_{q,i}}$, respectively; $\sigma_D^2$, $\mu_{s_D}$ is the variance and mean of all $S_{D_{q,i}}$, respectively; $\sigma_E^2$, $\mu_{s_E}$ is the variance and mean of all $S_{E_{q,i}}$, respectively. Then the pairwise tcr2tcr score is the weighted sum of the three similarity scores:

$$S_{q,i} = w_T \cdot S_{T_{q,i}} + w_D \cdot S_{D_{q,i}} + w_E \cdot S_{E_{q,i}},\ i \in R_{pool}.$$

The reference TCR sequence with the highest $S_{q,i}$ score is deemed the most similar TCR sequence to the query TCR sequence.

#### Preparing Benchmark datasets and benchmark reference database

We collected four experimentally verified datasets of viral epitope-specific TCR sequences to evaluate the epitope prediction performance of TCRanno and four state-of-the-art approaches: TCRMatch (3), Needleman-Wunsch Hamming distance (4), tcrdist and Levenshtein/edit distance (5).

i) PRJNA577794 (6), a multi-omic study of TCR response upon yellow fever vaccinations. We selected the yellow fever-specific CD8+ T cells (CD3+CD8+A02-NS4b221-227+) and downloaded the corresponding raw TCRseq data (fastq.gz). TCR sequences were extracted using Mixcr (7) with default parameters. Sequences with length less than 4 aa or more than 30 aa were excluded. Sequences identical to the TCR sequences within the reference database were also excluded, resulting in 2112 unique TCR sequences and their cognate epitopes.

ii) PRJNA778775 (8), an ex vivo characterization of CD8 T cell responses to SARS-CoV2 epitopes presented by four major HLA class I alleles. We downloaded the experimentally verified single-cell TCR sequences against various SARS-CoV2 epitopes (some other viral epitopes as control were also included) from the original paper and extracted the TCR-epitope pairs. Sequences with length less than 4 aa or more than 30 aa were excluded. Sequences identical to the TCR sequences within the reference database were also excluded, resulting in 859 unique TCR sequences and their cognate epitopes.

iii) immuneACCESS: B7W88F (9), a study demonstrating broad recognition of influenza M1 by different TCRs in HLA-A2+ individuals. We downloaded all the TCRb repertoire data of CD8 T cells recognizing HLA-A2/M1. Sequences with length less than 4 aa or more than 30 aa were excluded. Sequences identical to the TCR sequences within the reference database were also excluded, resulting in 377 unique TCR sequences and their cognate epitopes.

iv) immuneACCESS: AG2020MBIO (10), a study of EBV-specific TCR clonotypes in acute infectious mononucleosis. We downloaded all the TCRb repertoire data of CD8 T cells recognizing HLA-A2/BRLF1-109 and HLA-A2/BMLF1-280. Sequences with length less than 4 aa or more than 30 aa were excluded. Sequences identical to the TCR sequences within the reference database were also excluded, resulting in 368 unique TCR sequences and their cognate epitopes.

We used the IEDB database (11), which was originally used as reference data by TCRMatch, as the benchmark reference database. The TCR sequences in the original TCRMatch IEDB reference data was not in the standard C-F format. We need to transfer the data back to the standard C-F format to run the benchmark test on TCRMatch without error. We therefore took the TCR sequences from the full reference database that map to the original TCRMatch IEDB reference data, resulting in 126,797 unique TCR sequences, as the benchmark reference database.

**Preparing TCR Repertoire Annotation Data**

We performed TCR repertoire annotations and multi-group comparisons on a total of 4,195 repertoires curated from 18 studies described below. The repertoires were all sampled from periphery blood mononuclear cells (PBMC) and classified into five groups of populations with different conditions.

Group 1: healthy individuals with different CMV serostatus (n=786). TCR Repertoire data was obtained from i) immuneACCESS: B7001Z (12), containing two cohorts of healthy individuals with CMV serostatus determined experimentally as CMV+ (n=340), CMV- (n=420) and CMV unknown (n=26). Metadata recording CMV serostatus was obtained from the original paper, and TCR repertoires of the 786 individuals were retrieved from the immuneACCESS database.

Group 2: healthy individuals sampled pre- and post-vaccination of the COVID-19 vaccine AZD1222 (n=361). TCR Repertoire data was obtained from ii) immuneACCESS: PAS2021STM (13). A total of 361 repertoires (pre-vaccination: n=224; post-vaccination: n=137) were retrieved from the immuneACCESS database.

Group 3: SARS-CoV2 exposed/infected individuals (n=1485). TCR Repertoire data was obtained from iii) immuneACCESS: ADPT2020COVID (14), containing two cohorts of individuals exposed to or infected with SARS-CoV2. A total of 1485 TCR repertoires were retrieved from the immuneACCESS database. A small number of samples were further labelled with the clinical outcomes as either convalescent (n=62) or deceased (n=40) in the metadata files.

Group 4: systemic lupus erythematosus (SLE) patients (n=877). TCR Repertoire data was obtained from iv) pan immune repertoire database: P18081001 (15). A total of 877 SLE repertoires were retrieved from the study.

Group 5: various type of solid-tumor patients (n=686), including melanoma (n=196), lung cancer (n=209), urothelial bladder cancer (n=90), breast cancer (n=73), colorectal cancer (n=44), glioblastoma (n=28), head and neck cancer (n=20), basal cell and squamous cell carcinoma (n=13), renal cell carcinoma (n=9), ovarian cancer (n=4). Cancer types of less than 50 samples were grouped together into `other cancers' (n=118). TCR Repertoire data was obtained from datasets specified below.

v) immuneACCESS: EY2019CIR (16), melanoma. A total of 75 TCR repertoires (filtering non-PBMC samples) were retrieved from the dataset.

vi) immuneACCESS: SV2020NM (17), melanoma. A total of 58 TCR repertoires (filtering non-PBMC samples) were retrieved from the dataset.

vii) immuneACCESS: B7G592 (18), melanoma. A total of 42 TCR repertoires were retrieved from the dataset.

viii) immuneACCESS: JH2022NC (19), melanoma. A total of 21 TCR repertoires were retrieved from the dataset.

ix) immuneACCESS: B7BW6X (20), lung cancer samples receiving radiation therapy plus CTLA-4 blockade. A total of 69 TCR repertoires were retrieved from the study.

x) immuneACCESS: AR2019NC (21), localized non-small cell lung cancer samples. A total of 121 TCR repertoires were retrieved from the dataset.

xi) immuneACCESS: B7D59F (22), glioblastoma samples pre- and post-treatment of dendritic cell vaccination. A total of 28 TCR repertoires were retrieved from the study.

xii) immuneACCESS: B7WW2C (23), colorectal cancer samples receiving anti-PD1 (Pembrolizumab) treatment. A total of 24 TCR repertoires were retrieved from the study.

xiii) immuneACCESS:B7TG64 (24), ovarian cancer samples. A total of 4 TCR repertoires (filtering non-PBMC samples) were retrieved from the dataset.

xiv) immuneACCESS: B7MG68 (25), urothelial bladder cancers. A total of 90 TCR repertoires were retrieved from the dataset.

xv) immuneACCESS: ADPT2020V4CD (26), immunoSEQ hsTCRB-V4b control data. A total of 59 TCR repertoires of various cancers (19 lung cancer, 20 head and neck cancer, 20 colorectal cancer) were retrieved from the dataset.

xvi) immuneACCESS: DBP2016CIR (27), breast cancers. A total of 73 TCR repertoires (filtering non-PBMC samples) were retrieved from the dataset.

xvii) immuneACCESS: B78G65 (28), renal cell carcinoma. A total of 9 TCR repertoires (filtering non-PBMC samples) were retrieved from the dataset.

xviii) immuneACCESS: KY2019NM (29), basal cell carcinoma and squamous cell carcinoma. A total of 13 TCR repertoires (filtering non-PBMC samples) were retrieved from the dataset.

#### References

1. Dean J, Emerson RO, Vignali M, Sherwood AM, Rieder MJ, Carlson CS, et al. Annotation of pseudogenic gene segments by massively parallel sequencing of rearranged lymphocyte receptor loci. Genome Med. 2015;7(1):1–8.

2. Giudicelli V, Chaume D, Lefranc MP. IMGT/GENE-DB: a comprehensive database for human and mouse immunoglobulin and T cell receptor genes. Nucleic Acids Res. 2005;33(Database issue).

3. Chronister WD, Crinklaw A, Mahajan S, Vita R, Koşaloğlu-Yalçın Z, Yan Z, et al. TCRMatch: Predicting T-Cell Receptor Specificity Based on Sequence Similarity to Previously Characterized Receptors. Front Immunol. 2021 Mar 11;12:673.

4. Daily J. Parasail: SIMD C library for global, semi-global, and local pairwise sequence alignments. BMC Bioinformatics. 2016;17(1):1–11.

5. Mayer-Blackwell K, Schattgen S, Cohen-Lavi L, Crawford JC, Souquette A, Gaevert JA, et al. TCR meta-clonotypes for biomarker discovery with tcrdist3 enabled identification of public, HLA-restricted clusters of SARS-CoV-2 TCRs. Elife. 2021;10.

6. Minervina AA, Pogorelyy M V., Komech EA, Karnaukhov VK, Bacher P, Rosati E, et al. Primary and secondary anti-viral response captured by the dynamics and phenotype of individual T cell clones. Elife. 2020 Feb 1;9.

7. Bolotin DA, Poslavsky S, Mitrophanov I, Shugay M, Mamedov IZ, Putintseva E V., et al. MiXCR: software for comprehensive adaptive immunity profiling. Nat Methods. 2015;12(5):380–1.

8. Francis JM, Leistritz-Edwards D, Dunn A, Tarr C, Lehman J, Dempsey C, et al. Allelic variation in class I HLA determines CD8 + T cell repertoire shape and cross-reactive memory responses to SARS-CoV-2 MGH COVID-19 Collection and Processing Team. Sci Immunol. 2022;7:3070.

9. Song IY, Gil A, Mishra R, Ghersi D, Selin LK, Stern LJ. Broad TCR repertoire and diverse structural solutions for recognition of an immunodominant CD8+ T cell epitope. Nat Struct Mol Biol 2017;24(4):395–406.

10. Gil A, Kamga L, Chirravuri-Venkata R, Aslan N, Clark F, Ghersi D, et al. Epstein-Barr Virus Epitope–Major Histocompatibility Complex Interaction Combined with Convergent Recombination Drives Selection of Diverse T Cell Receptor α and β Repertoires. MBio. 2020;11(2).

11. Vita R, Mahajan S, Overton JA, Dhanda SK, Martini S, Cantrell JR, et al. The Immune Epitope Database (IEDB): 2018 update. Nucleic Acids Res. 2019;47(D1):D339–43.

12. Emerson RO, DeWitt WS, Vignali M, Gravley J, Hu JK, Osborne EJ, et al. Immunosequencing identifies signatures of cytomegalovirus exposure history and HLA-mediated effects on the T cell repertoire. Nat Genet. 2017;49(5):659–65.

13. Swanson PA, Padilla M, Hoyland W, McGlinchey K, Fields PA, Bibi S, et al. AZD1222/ChAdOx1 nCoV-19 vaccination induces a polyfunctional spike protein-specific TH1 response with a diverse TCR repertoire. Sci Transl Med. 2021;13(620):7211.

14. Nolan S, Vignali M, Klinger M, Dines JN, Kaplan IM, Svejnoha E, et al. A large-scale database of T-cell receptor beta (TCRβ) sequences and binding associations from natural and synthetic exposure to SARS-CoV-2. Res Sq. 2020.

15. Liu X, Zhang W, Zhao M, Fu L, Liu L, Wu J, et al. T cell receptor β repertoires as novel diagnostic markers for systemic lupus erythematosus and rheumatoid arthritis. Ann Rheum Dis. 2019;78(8):1070–8.

16. Yusko E, Vignali M, Wilson RK, Mardis ER, Stephen Hodi F, Horak C, et al. Association of Tumor Microenvironment T-cell Repertoire and Mutational Load with Clinical Outcome after Sequential Checkpoint Blockade in Melanoma. Cancer Immunol Res. 2019;7(3):458–65.

17. Valpione S, Galvani E, Tweedy J, Mundra PA, Banyard A, Middlehurst P, et al. Immune-awakening revealed by peripheral T cell dynamics after one cycle of immunotherapy. Nat cancer. 2020;1(2):210–21.

18. Robert L, Tsoi J, Wang X, Emerson R, Homet B, Chodon T, et al. CTLA4 blockade broadens the peripheral T-cell receptor repertoire. Clin Cancer Res. 2014;20(9):2424–32.

19. Huuhtanen J, Chen L, Jokinen E, Kasanen H, Lönnberg T, Kreutzman A, et al. Evolution and modulation of antigen-specific T cell responses in melanoma patients. Nat Commun. 2022;13(1).

20. Formenti SC, Rudqvist NP, Golden E, Cooper B, Wennerberg E, Lhuillier C, et al. Radiotherapy induces responses of lung cancer to CTLA-4 blockade. Nat Med. 2018;24(12):1845–51.

21. Reuben A, Zhang J, Chiou SH, Gittelman RM, Li J, Lee WC, et al. Comprehensive T cell repertoire characterization of non-small cell lung cancer. Nat Commun. 2020;11(1).

22. Melody SH, Shaina S, Tina W, Joseph PA, Richard GE, Alexander MT, et al. TCR Sequencing Can Identify and Track Glioma-Infiltrating T Cells after DC Vaccination. Cancer Immunol Res. 2016;4(5):412–8.

23. Le DT, Durham JN, Smith KN, Wang H, Bartlett BR, Aulakh LK, et al. Mismatch repair deficiency predicts response of solid tumors to PD-1 blockade. Science. 2017;357(6349):409–13.

24. Emerson RO, Sherwood AM, Rieder MJ, Guenthoer J, Williamson DW, Carlson CS, et al. High-throughput sequencing of T-cell receptors reveals a homogeneous repertoire of tumour-infiltrating lymphocytes in ovarian cancer. J Pathol. 2013;231(4):433–40.

25. Snyder A, Nathanson T, Funt SA, Ahuja A, Buros Novik J, Hellmann MD, et al. Contribution of systemic and somatic factors to clinical response and resistance to PD-L1 blockade in urothelial cancer: An exploratory multi-omic analysis. PLoS Med. 2017;14(5).

26. Hamm, DE. immunoSEQ hsTCRB-V4b Control Data. immuneACCESS. Adaptive Biotechnologies, Seattle WA. https://doi.org/10.21417/ADPT2020V4CD.

27. Page DB, Yuan J, Redmond D, Wen YH, Durack JC, Emerson R, et al. Deep Sequencing of T-cell Receptor DNA as a Biomarker of Clonally Expanded TILs in Breast Cancer after Immunotherapy. Cancer Immunol Res. 2016;4(10):835–44.

28. Wallin JJ, Bendell JC, Funke R, Sznol M, Korski K, Jones S, et al. Atezolizumab in combination with bevacizumab enhances antigen-specific T-cell migration in metastatic renal cell carcinoma. Nat Commun. 2016;7.

29. Yost KE, Satpathy AT, Wells DK, Qi Y, Wang C, Kageyama R, et al. Clonal replacement of tumor-specific T cells following PD-1 blockade. Nat Med. 2019;25(8):1251–9.

### Supplementary Figures

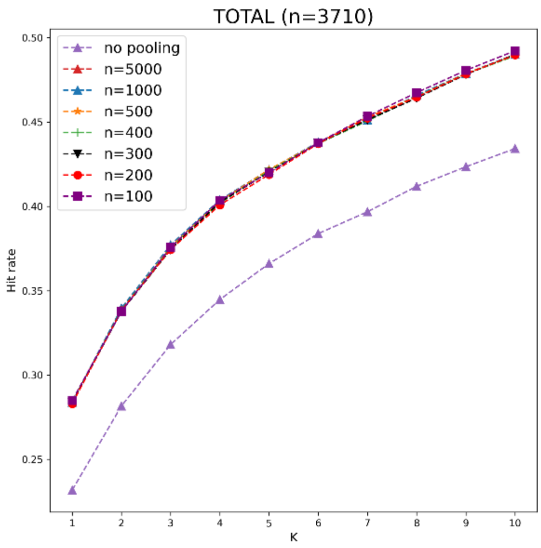

S1. Selecting the pooling parameter n.
When n=100, TCRanno showed the best performance and run fastest on the merged total of the benchmark datasets.

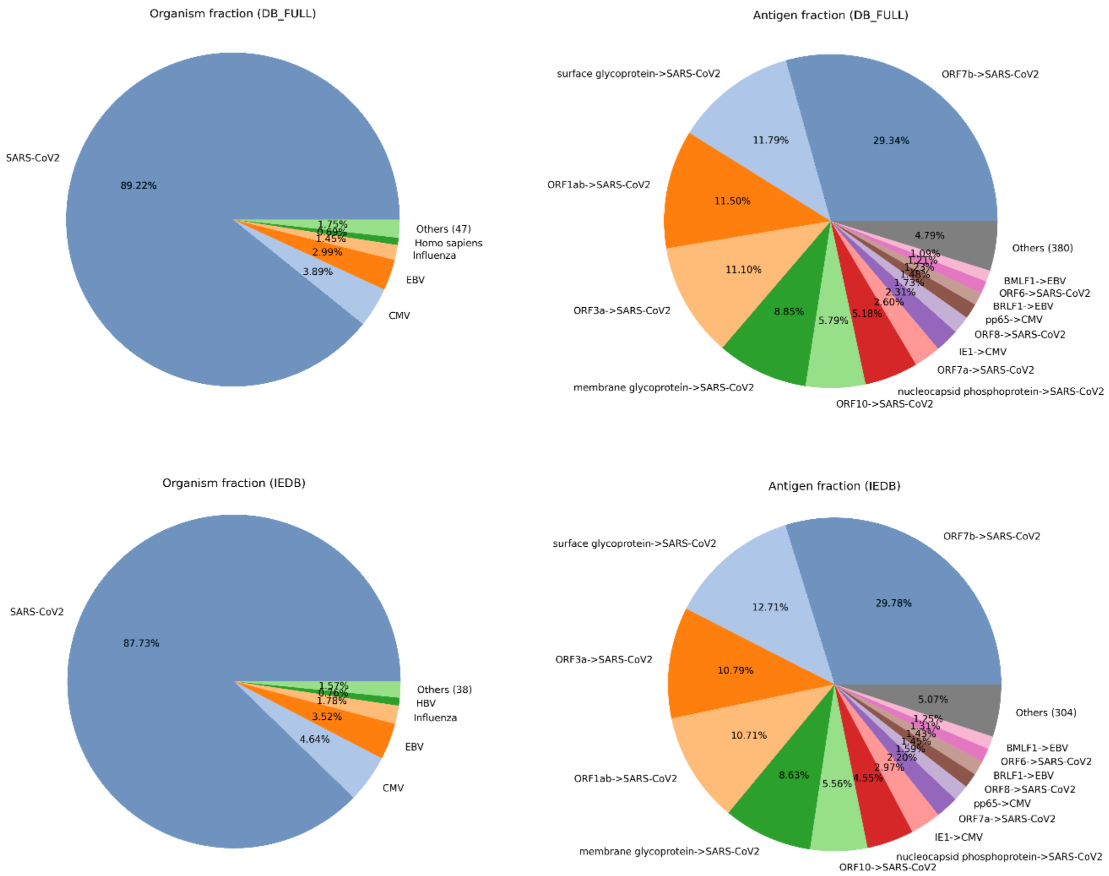

S2. Pie charts showing the frequency distribution of organisms and antigens in the full reference database and the IEDB benchmark database.

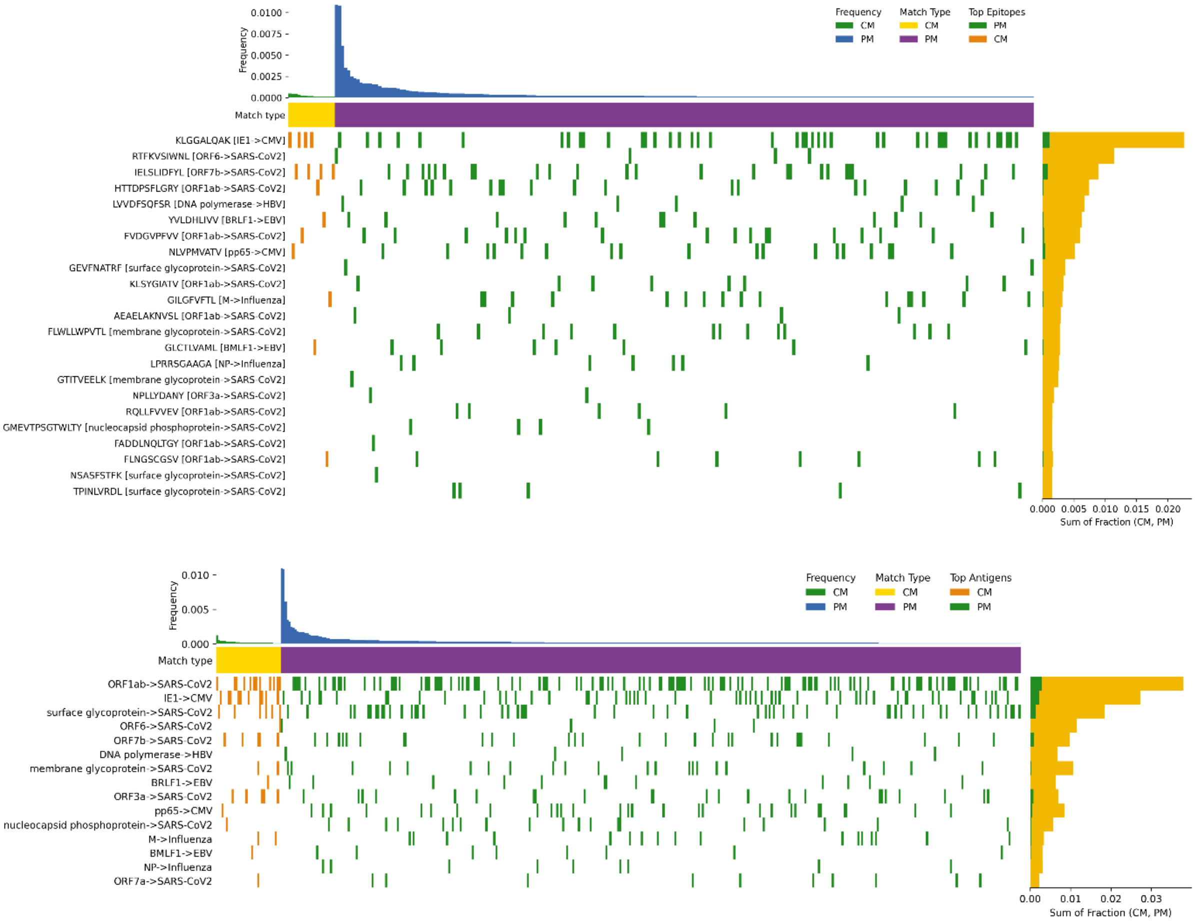

S3. Example visualization plots as a graphical summary for specificity landscape of a repertoire. Plots are generated at three levels: epitope (top), antigen (bottom) and organism (in main Figure 1c).

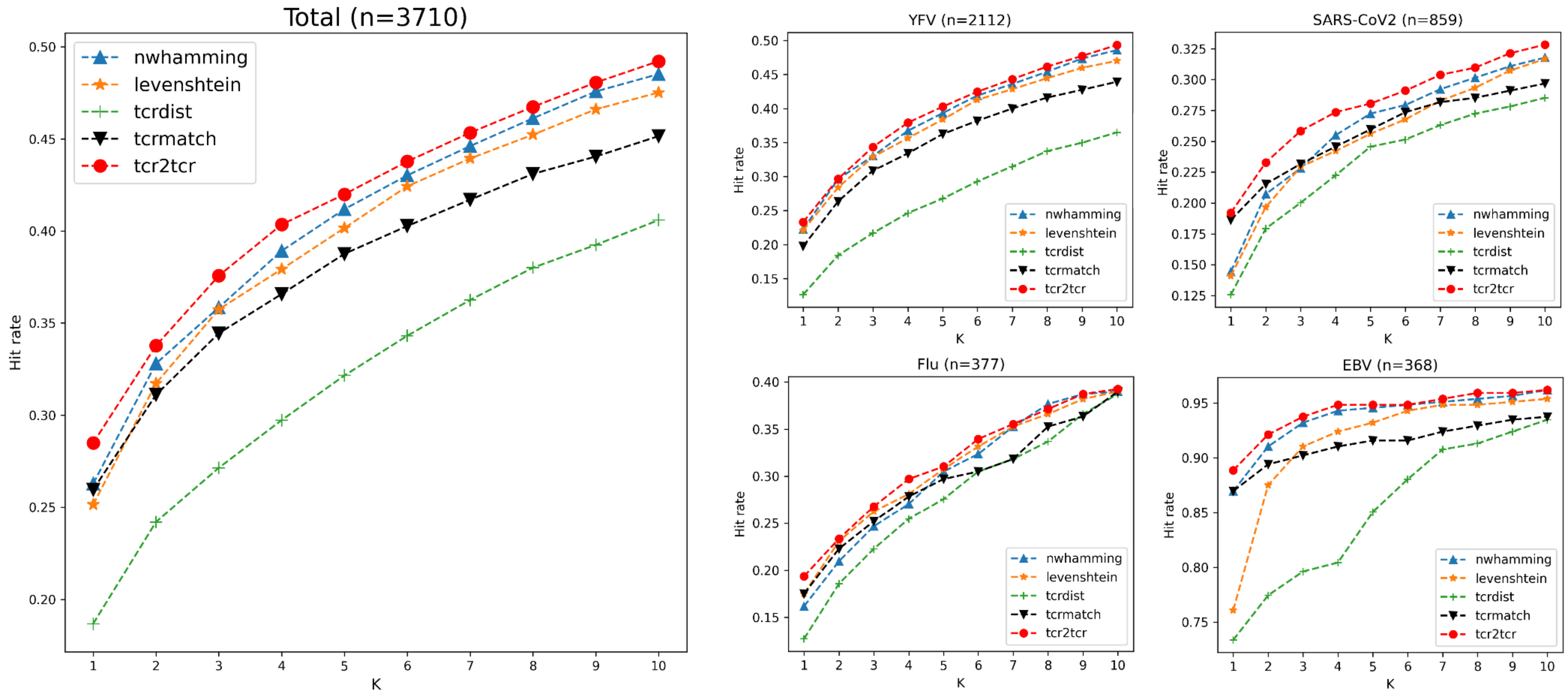

S4. Benchmark performances of TCRanno and state-of-the-art methods on four benchmark datasets.
A hit is a correct prediction of the cognate epitope (with top K choices) for the query TCR sequence. The hit rate is the number of hits dividing the total number of query TCR sequences. The left figure shows hit rates on the merged total of the four test sets. The right figures show hit rates on each test set.

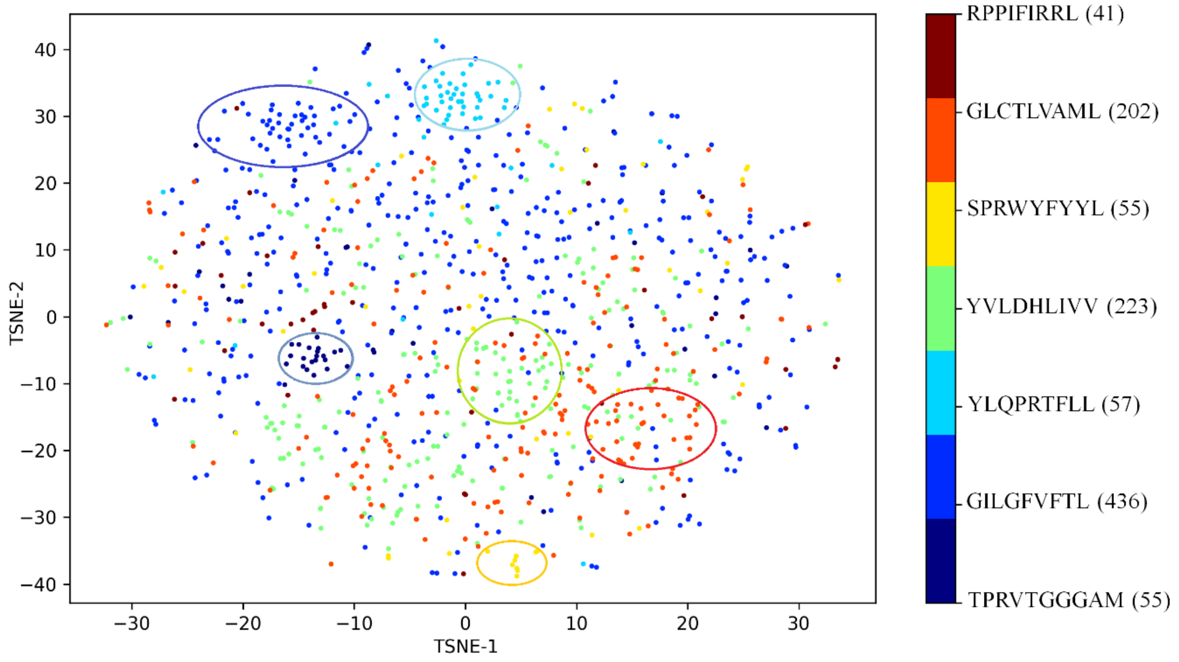

S5. Visualization of latent space class separation on benchmark data.
The merged total of the four benchmark datasets contain 3710 unique TCR sequences specific to 122 epitope classes. We filtered classes with too big or too small sizes and selected epitope classes of size between 30 and 500 (i.e., 30-500 TCR sequences specific to the epitope; resulting in 7 epitope classes) for t-SNE 2D visualization of the embeddings (32-dim vectors). The t-SNE plot showed appreciable intragroup aggregation (highlighted with ellipses) in most of the classes.

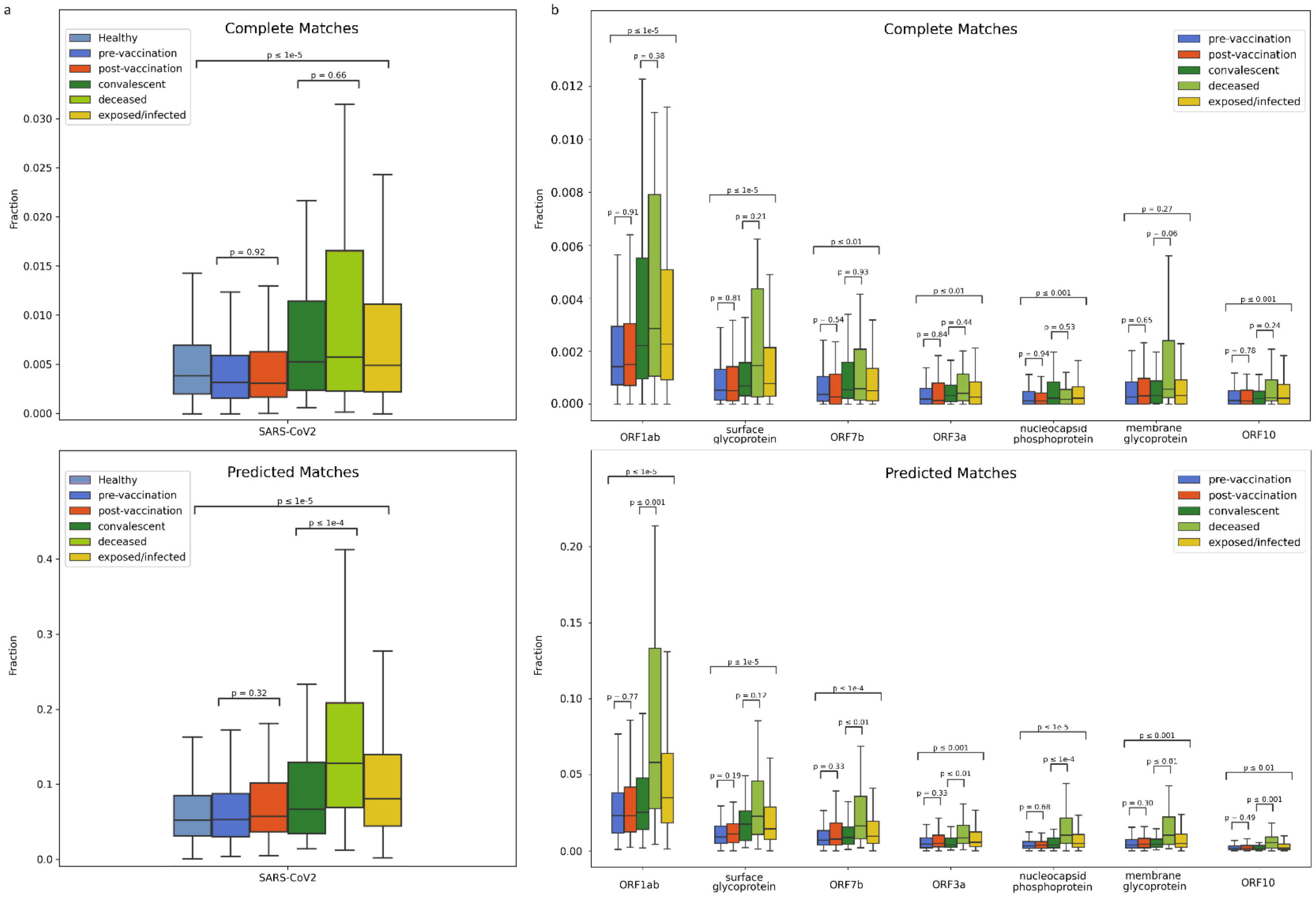

S6. Fractions of SARS-CoV2-specific TCRs in different sub-populations for CM and PM dimensions.
(a) SARS-CoV2-specific TCR fractions in healthy (n=786), pre-vaccination (n=224), post-vaccination (n=137), SARS-CoV2 convalescent (n=62), deceased (n=40) and exposed/infected (n=1485), for both complete matches (top) and predicted matches (bottom). (b) Fractions of TCRs specific to major antigens of SARS-CoV2 in pre-vaccination, post-vaccination, SARS-CoV2 convalescent, deceased and exposed/infected, for both complete matches (top) and predicted matches (bottom).

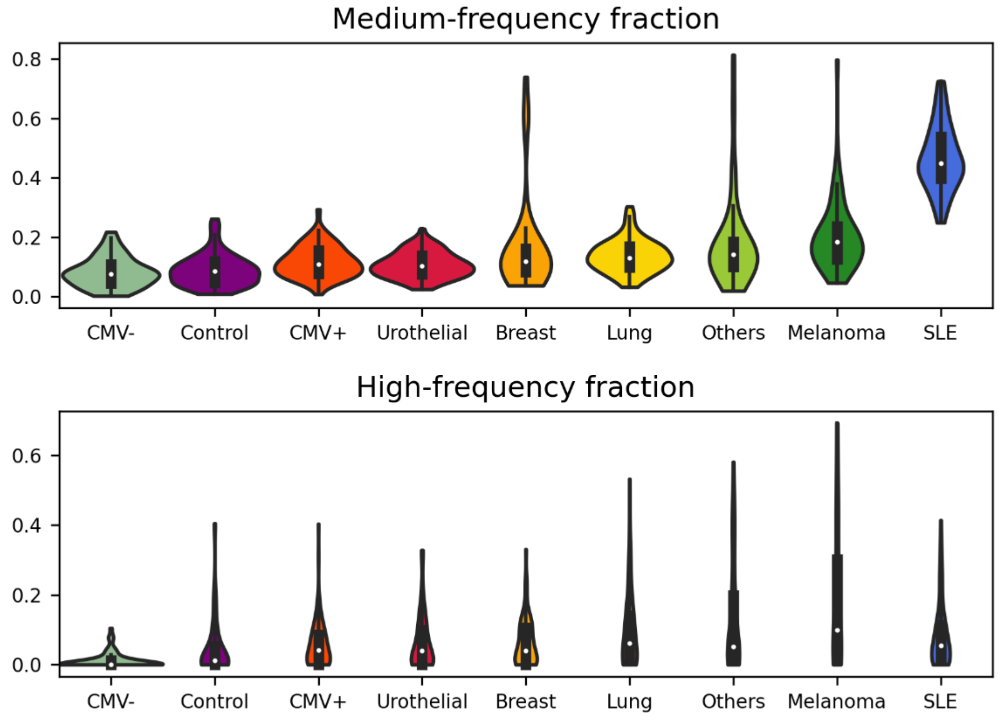

S7. Medium/High-frequency clonotype fractions in different sub-populations.
The effective fraction is consisted of medium-frequency clonotype fraction and high-frequency clonotype fraction.

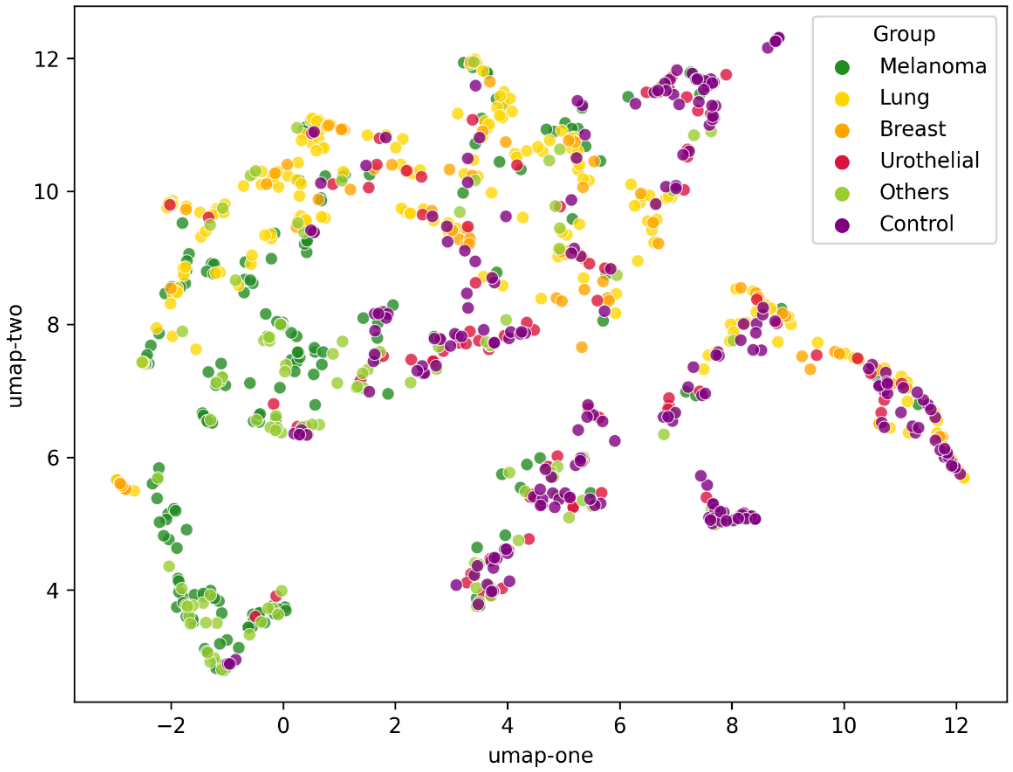

S8. UMAP plot of healthy control (n=224, pre-vaccination) vs different cancer types (n=686).

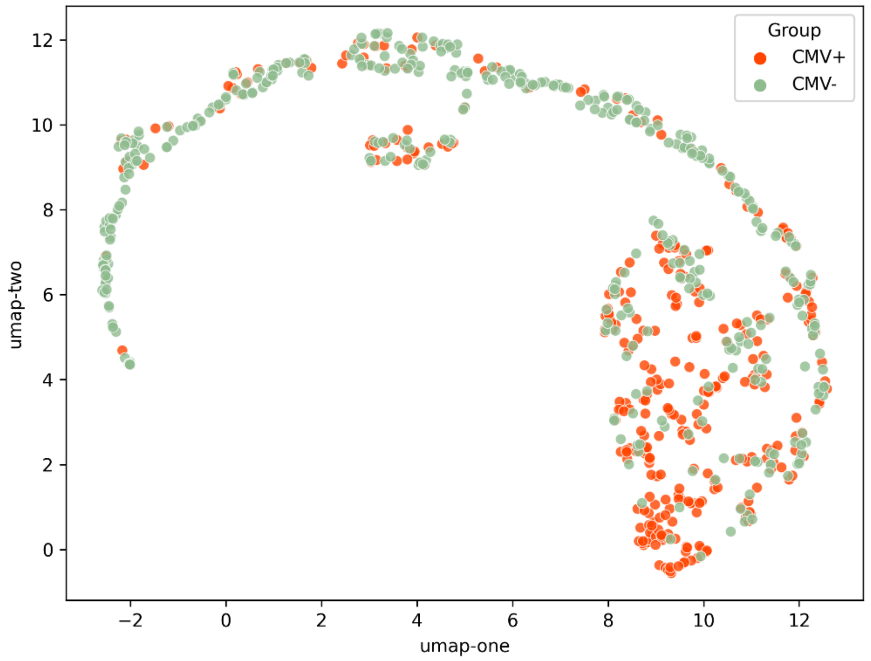

S9. UMAP plot of CMV+ (n=340) vs CMV- (n=420) individuals.

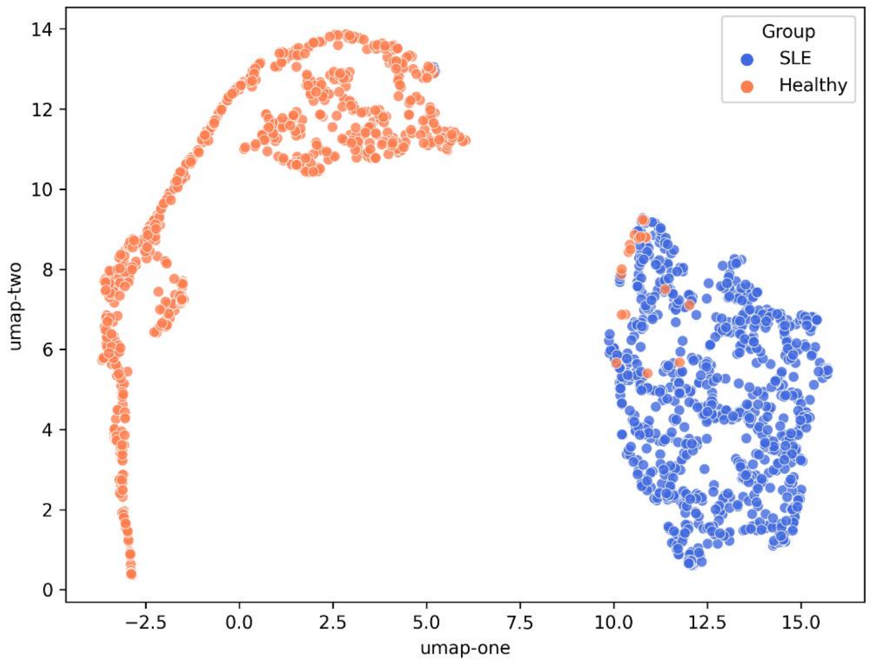

S10. UMAP plot of SLE patients (n=877) vs Healthy individuals (n=786).

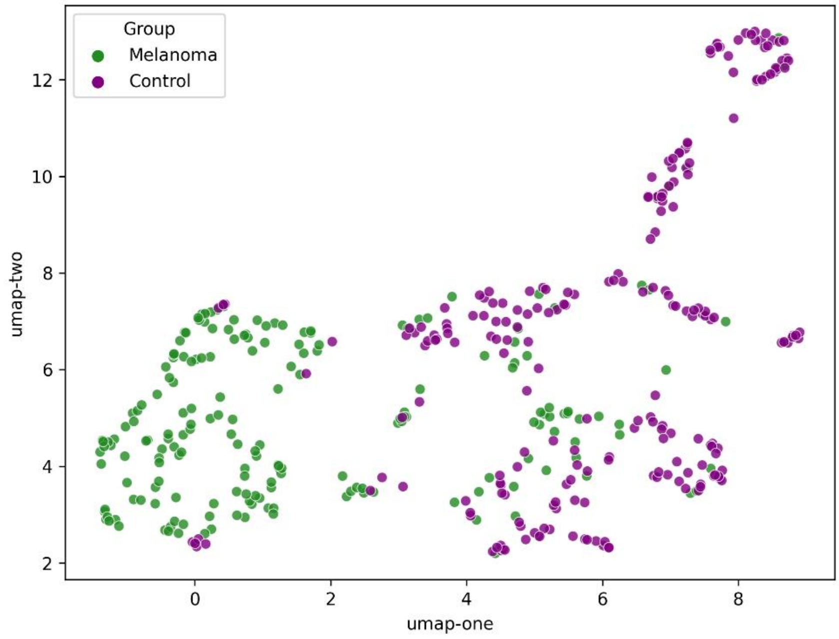

S11. UMAP plot of melanoma patients (n=196) vs healthy controls (n=224, pre-vaccination).

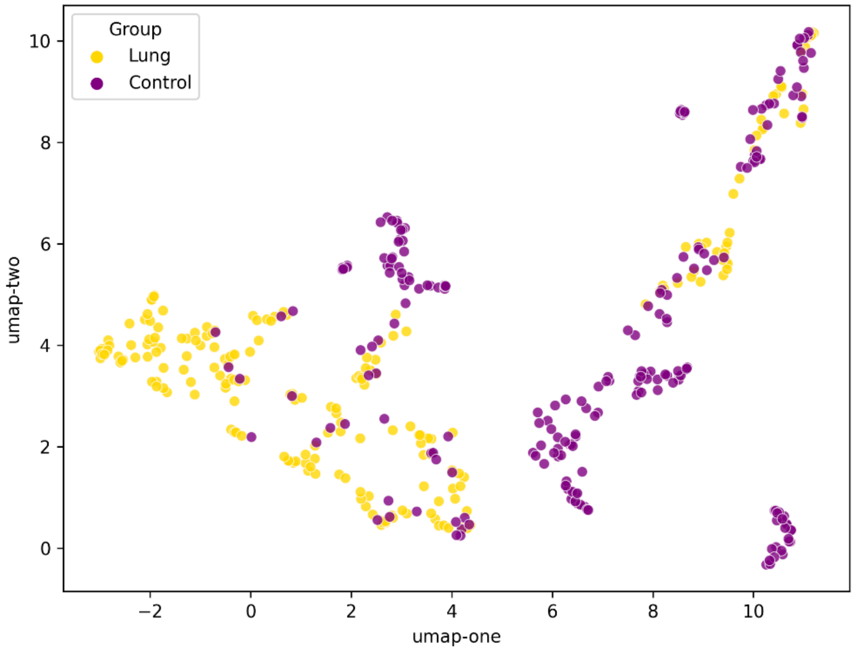

S12. UMAP plot of lung cancer patients (n=209) vs healthy controls (n=224, pre-vaccination).

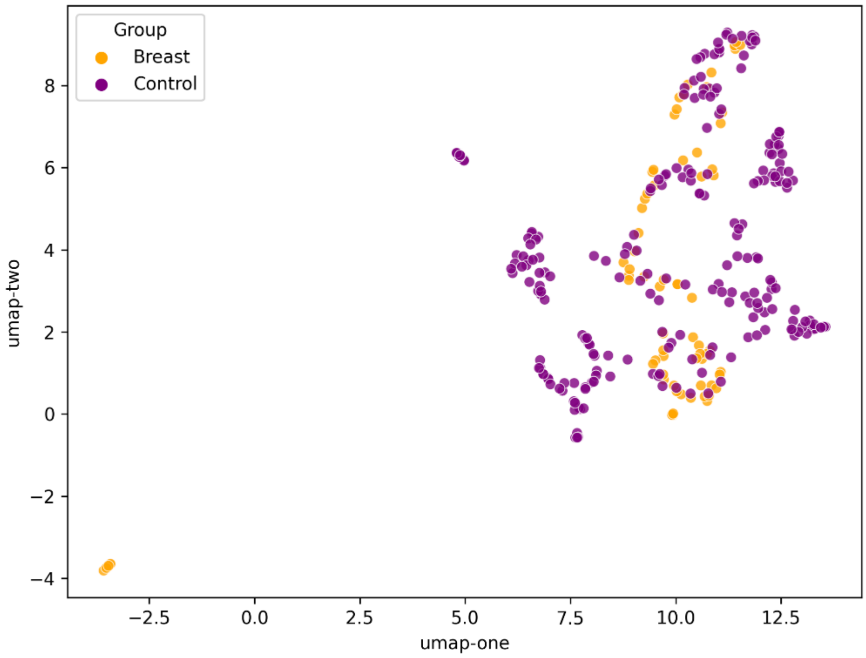

S13. UMAP plot of breast cancer patients (n=73) vs healthy controls (n=224, pre-vaccination).

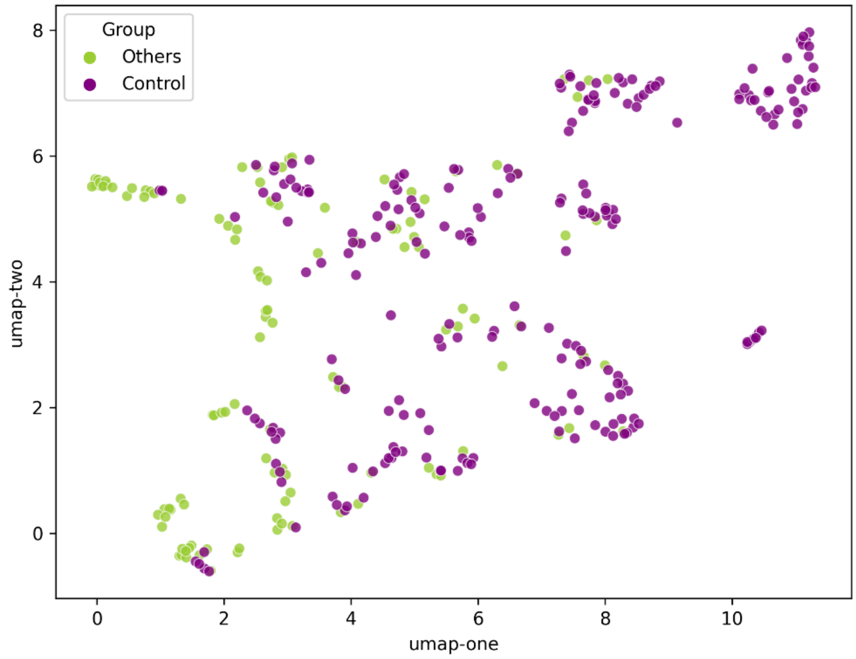

S14. UMAP plot of other cancers (n=118) vs healthy controls (n=224, pre-vaccination).

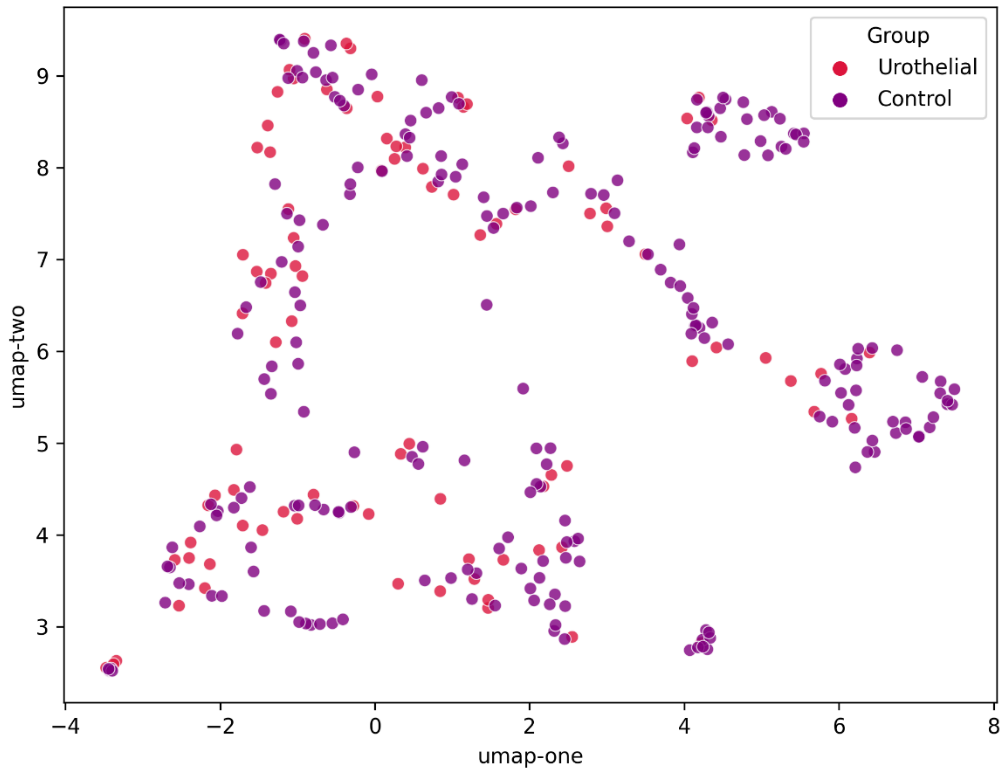

S15. UMAP plot of urothelial bladder cancer patients (n=90) vs healthy controls (n=224, pre-vaccination).

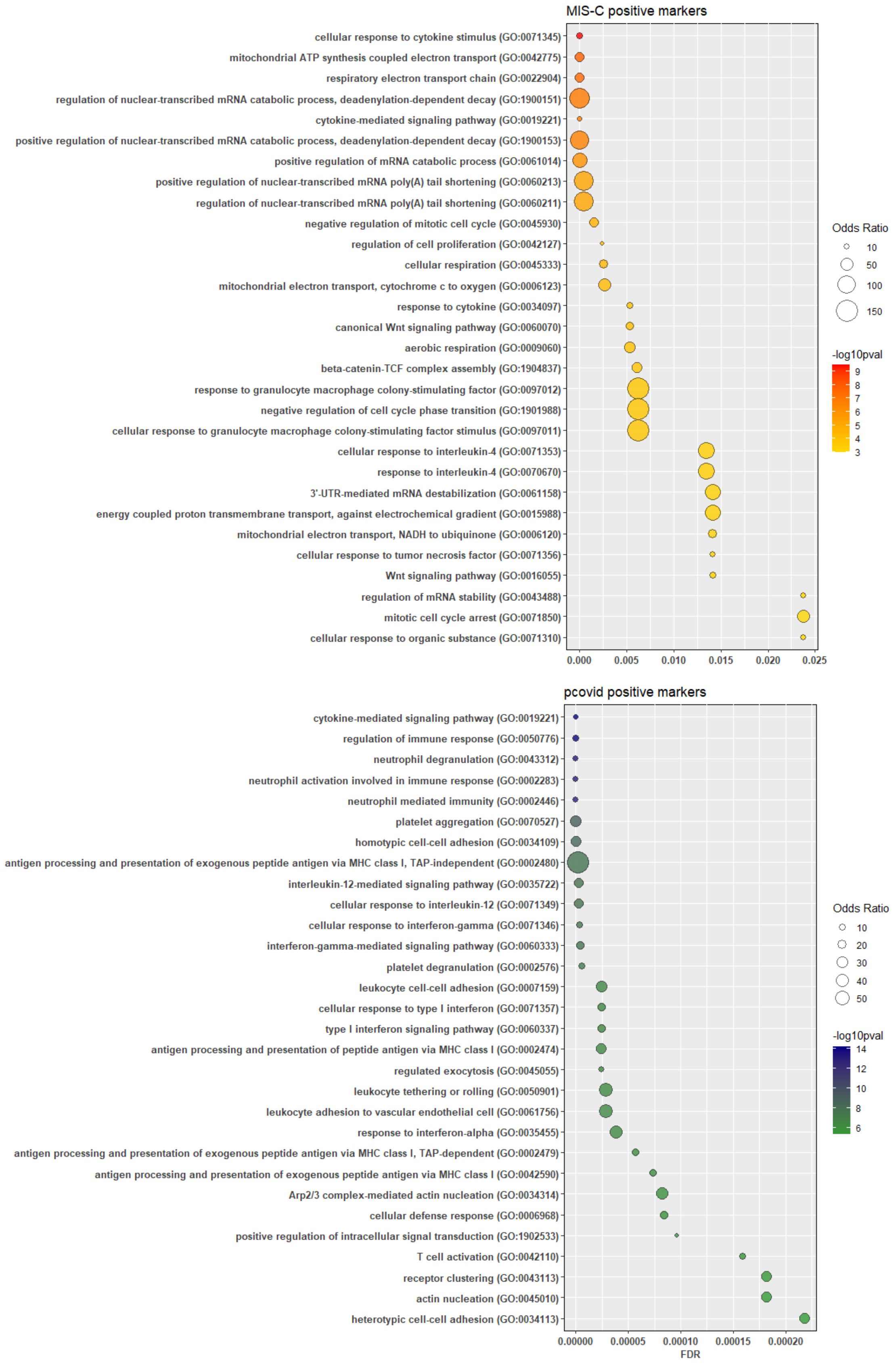

S16. Gene ontology enrichment analysis of differentially expressed genes in MIS-C vs pediatric COVID-19 (pcovid).

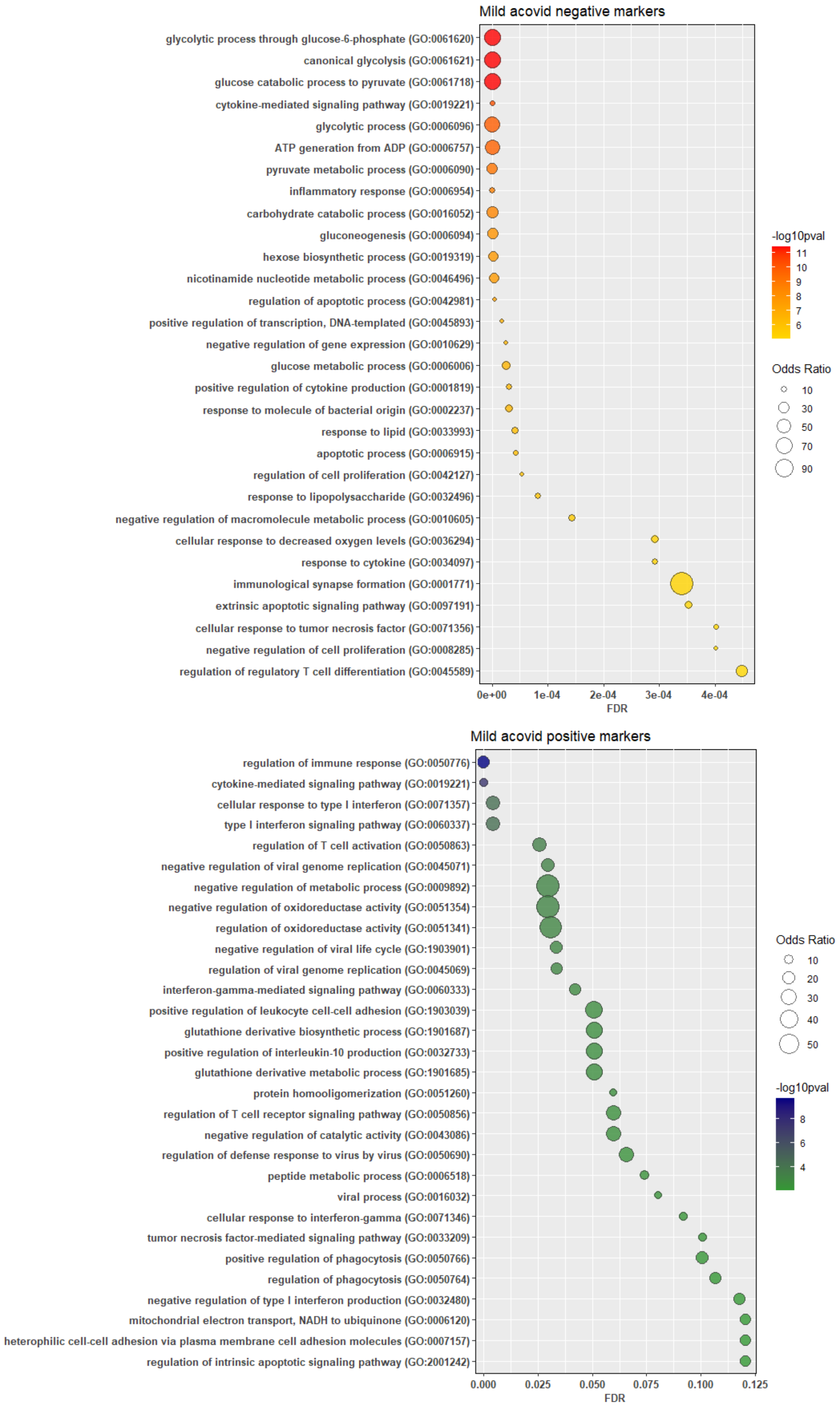

S17. Gene ontology enrichment analysis of differentially expressed genes in Severe vs Mild adult COVID-19 (acovid).

### Supplementary Tables

S1. Runtime performances of the benchmarked methods.

| Runtime (s) | Size = 100 | Size = 500 | Size = 1000 | Size = 5000 | Size = 10000 | Size = 50000 | Size = 100000 |
|---|---|---|---|---|---|---|---|
| tcr2tcr | 19.298±0.349 | 28.592±0.636 | **30.622±0.69** | **82.62±10.471** | **137.99±6.034** | **515.742±2.85** | **1032.446±20.178** |
| tcrmatch | **8.626±0.135** | **22.86±0.551** | 45.352±1.277 | 195.76±1.521 | 385.124±3.925 | 1912.728±15.631 | > 1 hour |
| nwhamming | 75.136±4.683 | 298.72±81.11 | 428.76±9.666 | 2169.406±41.059 | > 1 hour | > 5 hours | > 10 hours |
| tcrdist | 15.014±0.184 | 76.318±0.633 | 154.594±2.143 | 778.72±6.986 | 1590.206±15.067 | > 2 hours | > 4 hours |
| levenshtein | 16.046±0.092 | 82.162±0.452 | 166.142±2.027 | 835.8±11.478 | 1699.48±16.327 | > 2 hours | > 4 hours |

S2. Mean fraction (percentage) composition of TCR repertoire specificity of CMV+ individuals.

| Rank | CM:Top_Organisms (mean fraction) | PM:Top_Organisms (mean fraction) |
|---|---|---|
| 1 | SARS-CoV2(0.00955, 72.65%) | SARS-CoV2(0.10713, 66.03%) |
| 2 | **CMV(0.0045, 34.27%)** | **CMV(0.0308, 18.98%)** |
| 3 | EBV(0.00195, 14.8%) | EBV(0.0132, 8.14%) |
| 4 | Influenza(0.00169, 12.87%) | Influenza(0.01028, 6.34%) |
| 5 | Homo sapiens(0.00082, 6.21%) | HBV(0.00447, 2.75%) |
| 6 | HBV(0.00079, 5.99%) | YFV(0.00332, 2.04%) |
| 7 | YFV(0.00041, 3.14%) | Homo sapiens(0.00265, 1.64%) |
| 8 | Lymphocytic choriomeningitis virus(0.00039, 2.97%) | . |
| 9 | HIV(0.00019, 1.41%) | . |
| Total | 0.013139937 | 0.162247767 |

| Rank | CM:Top_Antigens (mean fraction) | PM:Top_Antigens (mean fraction) |
|---|---|---|
| 1 | ORF1ab->SARS-CoV2(0.00457, 34.77%) | ORF1ab->SARS-CoV2(0.0409, 25.21%) |
| 2 | surface glycoprotein->SARS-CoV2(0.00284, 21.61%) | surface glycoprotein->SARS-CoV2(0.0208, 12.82%) |
| 3 | **IE1->CMV(0.00283, 21.5%)** | **IE1->CMV(0.02012, 12.4%)** |
| 4 | ORF7b->SARS-CoV2(0.00228, 17.39%) | ORF7b->SARS-CoV2(0.01494, 9.21%) |
| 5 | **pp65->CMV(0.00171, 13.0%)** | ORF3a->SARS-CoV2(0.01166, 7.19%) |
| 6 | membrane glycoprotein->SARS-CoV2(0.00135, 10.26%) | **pp65->CMV(0.01074, 6.62%)** |
| 7 | ORF10->SARS-CoV2(0.00126, 9.62%) | membrane glycoprotein->SARS-CoV2(0.00879, 5.42%) |
| 8 | M->Influenza(0.00111, 8.48%) | nucleocapsid phosphoprotein->SARS-CoV2(0.00788, 4.86%) |
| 9 | ORF3a->SARS-CoV2(0.00107, 8.13%) | M->Influenza(0.00594, 3.66%) |
| 10 | nucleocapsid phosphoprotein->SARS-CoV2(0.00086, 6.53%) | BRLF1->EBV(0.00484, 2.98%) |
| 11 | ORF7a->SARS-CoV2(0.00083, 6.33%) | BMLF1->EBV(0.00422, 2.6%) |
| 12 | EBNA4->EBV(0.0008, 6.06%) | Nucleocapsid protein->Influenza(0.00403, 2.48%) |
| 13 | BMLF1->EBV(0.00066, 4.99%) | ORF10->SARS-CoV2(0.00366, 2.26%) |
| 14 | DNA polymerase->HBV(0.00064, 4.88%) | ORF7a->SARS-CoV2(0.00351, 2.17%) |
| 15 | Nucleocapsid protein->Influenza(0.00055, 4.22%) | NS4B->YFV(0.00332, 2.04%) |
| 16 | BRLF1->EBV(0.00045, 3.46%) | DNA polymerase->HBV(0.00321, 1.98%) |
| 17 | NS4B->YFV(0.00041, 3.14%) | EBNA4->EBV(0.00286, 1.76%) |
| 18 | MLANA->Homo sapiens(0.00041, 3.12%) | ORF8->SARS-CoV2(0.0022, 1.36%) |
| 19 | nucleoprotein->Lymphocytic choriomeningitis virus(0.00039, 2.96%) | precore/core protein->HBV(0.00174, 1.07%) |
| 20 | glycoprotein->Lymphocytic choriomeningitis virus(0.00031, 2.38%) | BZLF1->EBV(0.00098, 0.61%) |
| Total | 0.013139937 | 0.162247767 |

Mean fraction and percentage composition of repertoire specificity to organisms and antigens of CMV+ individuals (n=340). Top 10 organisms with CM/PM percentage no less than 1% is shown; top 20 antigens with CM/PM percentage no less than 0.5% is shown. The total fraction is the sum of fraction for all CM/PM sequences. Significant findings are highlighted with bold font.

S3. Mean fraction (percentage) composition of TCR repertoire specificity of CMV- individuals.

| Rank | CM:Top_Organisms (mean fraction) | PM:Top_Organisms (mean fraction) |
|---|---|---|
| 1 | SARS-CoV2(0.00522, 71.3%) | SARS-CoV2(0.05615, 66.96%) |
| 2 | **CMV(0.00222, 30.31%)** | **CMV(0.01641, 19.57%)** |
| 3 | EBV(0.00153, 20.86%) | EBV(0.00686, 8.17%) |
| 4 | Influenza(0.00082, 11.18%) | Influenza(0.0053, 6.32%) |
| 5 | Homo sapiens(0.00071, 9.63%) | HBV(0.00192, 2.29%) |
| 6 | YFV(0.00028, 3.89%) | YFV(0.00143, 1.71%) |
| 7 | HBV(0.00018, 2.46%) | Homo sapiens(0.00133, 1.59%) |
| Total | 0.007323402 | 0.083862 |

| Rank | CM:Top_Antigens (mean fraction) | PM:Top_Antigens (mean fraction) |
|---|---|---|
| 1 | ORF1ab->SARS-CoV2(0.0028, 38.28%) | ORF1ab->SARS-CoV2(0.0224, 26.7%) |
| 2 | **IE1->CMV(0.00165, 22.53%)** | **IE1->CMV(0.01105, 13.18%)** |
| 3 | surface glycoprotein->SARS-CoV2(0.00112, 15.31%) | surface glycoprotein->SARS-CoV2(0.01044, 12.45%) |
| 4 | ORF7b->SARS-CoV2(0.00098, 13.37%) | ORF7b->SARS-CoV2(0.00761, 9.08%) |
| 5 | membrane glycoprotein->SARS-CoV2(0.00074, 10.09%) | **pp65->CMV(0.00542, 6.46%)** |
| 6 | **pp65->CMV(0.00071, 9.76%)** | ORF3a->SARS-CoV2(0.00506, 6.03%) |
| 7 | BMLF1->EBV(0.00062, 8.49%) | membrane glycoprotein->SARS-CoV2(0.00497, 5.92%) |
| 8 | ORF3a->SARS-CoV2(0.00057, 7.82%) | nucleocapsid phosphoprotein->SARS-CoV2(0.00402, 4.8%) |
| 9 | nucleocapsid phosphoprotein->SARS-CoV2(0.00056, 7.66%) | M->Influenza(0.00291, 3.47%) |
| 10 | M->Influenza(0.00053, 7.27%) | BRLF1->EBV(0.00228, 2.72%) |
| 11 | ORF10->SARS-CoV2(0.00041, 5.57%) | ORF10->SARS-CoV2(0.00203, 2.42%) |
| 12 | EBNA4->EBV(0.00041, 5.55%) | Nucleocapsid protein->Influenza(0.00195, 2.32%) |
| 13 | EBNA3A->EBV(0.0004, 5.4%) | EBNA4->EBV(0.00184, 2.19%) |
| 14 | BZLF1->EBV(0.00033, 4.56%) | BMLF1->EBV(0.00183, 2.18%) |
| 15 | MLANA->Homo sapiens(0.00032, 4.36%) | ORF7a->SARS-CoV2(0.00155, 1.85%) |
| 16 | NS4B->YFV(0.00028, 3.88%) | NS4B->YFV(0.00143, 1.7%) |
| 17 | Nucleocapsid protein->Influenza(0.00027, 3.67%) | DNA polymerase->HBV(0.00128, 1.52%) |
| 18 | ORF7a->SARS-CoV2(0.0002, 2.79%) | ORF8->SARS-CoV2(0.00095, 1.13%) |
| 19 | BRLF1->EBV(0.0002, 2.72%) | BZLF1->EBV(0.00062, 0.73%) |
| 20 | PLA2G6->Homo sapiens(0.00013, 1.77%) | precore/core protein->HBV(0.0006, 0.71%) |
| Total | 0.007323402 | 0.083862414 |

Mean fraction and percentage composition of repertoire specificity to organisms and antigens of CMV- individuals (n=420). Top 10 organisms with CM/PM percentage no less than 1% is shown; top 20 antigens with CM/PM percentage no less than 0.5% is shown. The total fraction is the sum of fraction for all CM/PM sequences. Significant findings are highlighted with bold font.

S4. Mean fraction (percentage) composition of TCR repertoire specificity of healthy individuals.

| Rank | CM:Top_Organisms (mean fraction) | PM:Top_Organisms (mean fraction) |
|---|---|---|
| 1 | **SARS-CoV2(0.00715, 71.73%)** | **SARS-CoV2(0.07954, 66.42%)** |
| 2 | CMV(0.00324, 32.49%) | CMV(0.02292, 19.14%) |
| 3 | EBV(0.00171, 17.16%) | EBV(0.0097, 8.1%) |
| 4 | Influenza(0.00121, 12.11%) | Influenza(0.00756, 6.32%) |
| 5 | Homo sapiens(0.00079, 7.9%) | HBV(0.0031, 2.58%) |
| 6 | HBV(0.00044, 4.46%) | YFV(0.00233, 1.95%) |
| 7 | YFV(0.00035, 3.46%) | Homo sapiens(0.00197, 1.65%) |
| 8 | Lymphocytic choriomeningitis virus(0.00018, 1.84%) | . |
| 9 | HIV(0.00011, 1.11%) | . |
| Total | 0.009964824 | 0.119752 |

| Rank | CM:Top_Antigens (mean fraction) | PM:Top_Antigens (mean fraction) |
|---|---|---|
| 1 | ORF1ab->SARS-CoV2(0.0036, 36.17%) | ORF1ab->SARS-CoV2(0.03109, 25.96%) |
| 2 | IE1->CMV(0.00217, 21.78%) | surface glycoprotein->SARS-CoV2(0.01524, 12.73%) |
| 3 | surface glycoprotein->SARS-CoV2(0.00188, 18.89%) | IE1->CMV(0.01513, 12.63%) |
| 4 | ORF7b->SARS-CoV2(0.00156, 15.67%) | ORF7b->SARS-CoV2(0.01089, 9.09%) |
| 5 | pp65->CMV(0.00117, 11.71%) | ORF3a->SARS-CoV2(0.00797, 6.65%) |
| 6 | membrane glycoprotein->SARS-CoV2(0.001, 10.0%) | pp65->CMV(0.00783, 6.54%) |
| 7 | ORF3a->SARS-CoV2(0.00079, 7.92%) | membrane glycoprotein->SARS-CoV2(0.00667, 5.57%) |
| 8 | M->Influenza(0.00079, 7.92%) | nucleocapsid phosphoprotein->SARS-CoV2(0.00574, 4.79%) |
| 9 | ORF10->SARS-CoV2(0.00078, 7.83%) | M->Influenza(0.00428, 3.57%) |
| 10 | nucleocapsid phosphoprotein->SARS-CoV2(0.00068, 6.81%) | BRLF1->EBV(0.00341, 2.85%) |
| 11 | BMLF1->EBV(0.00062, 6.23%) | Nucleocapsid protein->Influenza(0.0029, 2.42%) |
| 12 | EBNA4->EBV(0.00058, 5.8%) | BMLF1->EBV(0.0029, 2.42%) |
| 13 | ORF7a->SARS-CoV2(0.0005, 4.99%) | ORF10->SARS-CoV2(0.0028, 2.34%) |
| 14 | Nucleocapsid protein->Influenza(0.0004, 3.98%) | ORF7a->SARS-CoV2(0.00246, 2.06%) |
| 15 | MLANA->Homo sapiens(0.00036, 3.59%) | NS4B->YFV(0.00233, 1.95%) |
| 16 | NS4B->YFV(0.00035, 3.46%) | EBNA4->EBV(0.00229, 1.91%) |
| 17 | DNA polymerase->HBV(0.00032, 3.26%) | DNA polymerase->HBV(0.00217, 1.81%) |
| 18 | BRLF1->EBV(0.00031, 3.09%) | ORF8->SARS-CoV2(0.00152, 1.27%) |
| 19 | BZLF1->EBV(0.00031, 3.07%) | precore/core protein->HBV(0.00111, 0.93%) |
| 20 | EBNA3A->EBV(0.0003, 3.05%) | BZLF1->EBV(0.00079, 0.66%) |
| Total | 0.009965 | 0.119752 |

Mean fraction and percentage composition of repertoire specificity to organisms and antigens of healthy individuals (n=786). Top 10 organisms with CM/PM percentage no less than 1% is shown; top 20 antigens with CM/PM percentage no less than 0.5% is shown. The total fraction is the sum of fraction for all CM/PM sequences. Significant findings are highlighted with bold font.

S5. Mean fraction (percentage) composition of TCR repertoire specificity of SARS-CoV2 exposed/infected individuals.

| Rank | CM:Top_Organisms (mean fraction) | PM:Top_Organisms (mean fraction) |
|---|---|---|
| 1 | **SARS-CoV2(0.0109, 71.18%)** | **SARS-CoV2(0.12614, 66.98%)** |
| 2 | CMV(0.0047, 30.7%) | CMV(0.03646, 19.36%) |
| 3 | EBV(0.00218, 14.21%) | EBV(0.01463, 7.77%) |
| 4 | Influenza(0.00152, 9.93%) | Influenza(0.01108, 5.88%) |
| 5 | Homo sapiens(0.00124, 8.07%) | HBV(0.00496, 2.63%) |
| 6 | HIV(0.00046, 2.99%) | Homo sapiens(0.00388, 2.06%) |
| 7 | YFV(0.00034, 2.25%) | YFV(0.00336, 1.78%) |
| 8 | HBV(0.00031, 2.02%) | . |
| Total | 0.015313336 | 0.188317 |

| Rank | CM:Top_Antigens (mean fraction) | PM:Top_Antigens (mean fraction) |
|---|---|---|
| 1 | ORF1ab->SARS-CoV2(0.00561, 36.66%) | ORF1ab->SARS-CoV2(0.05011, 26.61%) |
| 2 | surface glycoprotein->SARS-CoV2(0.00275, 17.94%) | surface glycoprotein->SARS-CoV2(0.02439, 12.95%) |
| 3 | pp65->CMV(0.00257, 16.78%) | IE1->CMV(0.02313, 12.28%) |
| 4 | IE1->CMV(0.00212, 13.85%) | ORF7b->SARS-CoV2(0.01656, 8.79%) |
| 5 | ORF7b->SARS-CoV2(0.00193, 12.6%) | pp65->CMV(0.01308, 6.94%) |
| 6 | ORF10->SARS-CoV2(0.00153, 9.99%) | ORF3a->SARS-CoV2(0.01118, 5.94%) |
| 7 | membrane glycoprotein->SARS-CoV2(0.00144, 9.39%) | nucleocapsid phosphoprotein->SARS-CoV2(0.00999, 5.31%) |
| 8 | ORF3a->SARS-CoV2(0.00127, 8.26%) | membrane glycoprotein->SARS-CoV2(0.00991, 5.26%) |
| 9 | nucleocapsid phosphoprotein->SARS-CoV2(0.00103, 6.75%) | M->Influenza(0.00584, 3.1%) |
| 10 | M->Influenza(0.00083, 5.42%) | BRLF1->EBV(0.00459, 2.44%) |
| 11 | BMLF1->EBV(0.00072, 4.69%) | ORF10->SARS-CoV2(0.00459, 2.43%) |
| 12 | ORF7a->SARS-CoV2(0.0007, 4.57%) | BMLF1->EBV(0.00457, 2.43%) |
| 13 | Nucleocapsid protein->Influenza(0.00064, 4.16%) | Nucleocapsid protein->Influenza(0.00456, 2.42%) |
| 14 | BRLF1->EBV(0.00052, 3.42%) | ORF7a->SARS-CoV2(0.00369, 1.96%) |
| 15 | EBNA4->EBV(0.0005, 3.24%) | EBNA4->EBV(0.00355, 1.88%) |
| 16 | MLANA->Homo sapiens(0.00045, 2.91%) | DNA polymerase->HBV(0.00336, 1.79%) |
| 17 | BZLF1->EBV(0.00042, 2.73%) | NS4B->YFV(0.00335, 1.78%) |
| 18 | EBNA3A->EBV(0.00034, 2.25%) | ORF8->SARS-CoV2(0.00271, 1.44%) |
| 19 | NS4B->YFV(0.00034, 2.25%) | precore/core protein->HBV(0.00165, 0.88%) |
| 20 | Gag->HIV(0.00034, 2.2%) | BZLF1->EBV(0.00129, 0.69%) |
| Total | 0.015313336 | 0.188317 |

Mean fraction and percentage composition of repertoire specificity to organisms and antigens of SARS-CoV2 exposed/infected individuals (n=1485). Top 10 organisms with CM/PM percentage no less than 1% is shown; top 20 antigens with CM/PM percentage no less than 0.5% is shown. The total fraction is the sum of fraction for all CM/PM sequences. Significant findings are highlighted with bold font.

S6. Mean fraction (percentage) composition of TCR repertoire specificity of COVID-19 deceased individuals.

| Rank | CM:Top_Organisms (mean fraction) | PM:Top_Organisms (mean fraction) |
|---|---|---|
| 1 | **SARS-CoV2(0.01644, 76.86%)** | **SARS-CoV2(0.18993, 67.44%)** |
| 2 | **Homo sapiens(0.00425, 19.87%)** | CMV(0.05166, 18.34%) |
| 3 | CMV(0.00375, 17.54%) | EBV(0.02811, 9.98%) |
| 4 | EBV(0.00334, 15.6%) | Influenza(0.01678, 5.96%) |
| 5 | Influenza(0.00319, 14.93%) | **Homo sapiens(0.00939, 3.33%)** |
| 6 | YFV(0.00083, 3.87%) | HBV(0.00486, 1.73%) |
| 7 | HIV(0.00055, 2.57%) | . |
| 8 | HBV(0.00032, 1.52%) | . |
| 9 | HCV(0.00027, 1.25%) | . |
| Total | 0.021383878 | 0.281627 |

| Rank | CM:Top_Antigens (mean fraction) | PM:Top_Antigens (mean fraction) |
|---|---|---|
| 1 | ORF1ab->SARS-CoV2(0.00879, 41.1%) | ORF1ab->SARS-CoV2(0.07832, 27.81%) |
| 2 | surface glycoprotein->SARS-CoV2(0.0067, 31.34%) | surface glycoprotein->SARS-CoV2(0.0319, 11.33%) |
| 3 | ORF7b->SARS-CoV2(0.00488, 22.83%) | IE1->CMV(0.02908, 10.32%) |
| 4 | membrane glycoprotein->SARS-CoV2(0.00408, 19.1%) | ORF7b->SARS-CoV2(0.0261, 9.27%) |
| 5 | ORF10->SARS-CoV2(0.00278, 12.98%) | membrane glycoprotein->SARS-CoV2(0.01959, 6.96%) |
| 6 | M->Influenza(0.00242, 11.31%) | pp65->CMV(0.01917, 6.81%) |
| 7 | pp65->CMV(0.00216, 10.11%) | nucleocapsid phosphoprotein->SARS-CoV2(0.01845, 6.55%) |
| 8 | IE1->CMV(0.00176, 8.22%) | ORF3a->SARS-CoV2(0.0128, 4.55%) |
| 9 | **AKAP13->Homo sapiens(0.00157, 7.32%)** | Nucleocapsid protein->Influenza(0.00936, 3.32%) |
| 10 | membrane glycoprotein,surface glycoprotein->SARS-CoV2(0.00138, 6.44%) | EBNA3A->EBV(0.00825, 2.93%) |
| 11 | **Kinetochore protein Nuf2->Homo sapiens(0.00136, 6.38%)** | ORF10->SARS-CoV2(0.00725, 2.57%) |
| 12 | BMLF1->EBV(0.00136, 6.37%) | BRLF1->EBV(0.00708, 2.51%) |
| 13 | ORF3a->SARS-CoV2(0.00134, 6.28%) | M->Influenza(0.00645, 2.29%) |
| 14 | BRLF1->EBV(0.00128, 5.97%) | BMLF1->EBV(0.00575, 2.04%) |
| 15 | **MLANA->Homo sapiens(0.00105, 4.93%)** | EBNA4->EBV(0.00563, 2.0%) |
| 16 | EBNA3A->EBV(0.00102, 4.78%) | ORF7a->SARS-CoV2(0.00481, 1.71%) |
| 17 | NS4B->YFV(0.00083, 3.87%) | DNA polymerase->HBV(0.00374, 1.33%) |
| 18 | EBNA6->EBV(0.00075, 3.49%) | IE2->CMV(0.00344, 1.22%) |
| 19 | **ABCD3->Homo sapiens(0.00075, 3.49%)** | **Sterol-4-alpha-carboxylate 3-dehydrogenase, decarboxylating->Homo sapiens(0.003, 1.06%)** |
| 20 | nucleocapsid phosphoprotein->SARS-CoV2(0.00073, 3.42%) | **GANAB->Homo sapiens(0.00267, 0.95%)** |
| Total | 0.021383878 | 0.281627 |

Mean fraction and percentage composition of repertoire specificity to organisms and antigens of COVID-19 deceased individuals (n=40). Top 10 organisms with CM/PM percentage no less than 1% is shown; top 20 antigens with CM/PM percentage no less than 0.5% is shown. The total fraction is the sum of fraction for all CM/PM sequences. Significant findings are highlighted with bold font.

S7. Mean fraction (percentage) composition of TCR repertoire specificity of COVID-19 convalescents.

| Rank | CM:Top_Organisms (mean fraction) | PM:Top_Organisms (mean fraction) |
|---|---|---|
| 1 | SARS-CoV2(0.00929, 74.05%) | SARS-CoV2(0.09841, 67.41%) |
| 2 | CMV(0.00383, 30.57%) | CMV(0.02677, 18.34%) |
| 3 | Influenza(0.00172, 13.71%) | EBV(0.01279, 8.76%) |
| 4 | EBV(0.00151, 12.06%) | Influenza(0.01117, 7.65%) |
| 5 | Homo sapiens(0.00053, 4.22%) | HBV(0.00324, 2.22%) |
| 6 | HBV(0.00044, 3.5%) | Homo sapiens(0.00308, 2.11%) |
| 7 | YFV(0.0002, 1.61%) | YFV(0.00227, 1.56%) |
| 8 | HIV(0.00019, 1.54%) | . |
| 9 | SARS coronavirus(0.00017, 1.34%) | . |
| Total | 0.012539543 | 0.145981 |

| Rank | CM:Top_Antigens (mean fraction) | PM:Top_Antigens (mean fraction) |
|---|---|---|
| 1 | ORF1ab->SARS-CoV2(0.00393, 31.36%) | ORF1ab->SARS-CoV2(0.03627, 24.85%) |
| 2 | surface glycoprotein->SARS-CoV2(0.00289, 23.04%) | surface glycoprotein->SARS-CoV2(0.02201, 15.08%) |
| 3 | pp65->CMV(0.00257, 20.52%) | IE1->CMV(0.01559, 10.68%) |
| 4 | ORF7b->SARS-CoV2(0.00168, 13.4%) | ORF7b->SARS-CoV2(0.01452, 9.95%) |
| 5 | IE1->CMV(0.00141, 11.27%) | pp65->CMV(0.01161, 7.95%) |
| 6 | ORF7a->SARS-CoV2(0.00126, 10.01%) | ORF3a->SARS-CoV2(0.00891, 6.11%) |
| 7 | ORF10->SARS-CoV2(0.00124, 9.88%) | membrane glycoprotein->SARS-CoV2(0.00857, 5.87%) |
| 8 | nucleocapsid phosphoprotein->SARS-CoV2(0.00102, 8.1%) | nucleocapsid phosphoprotein->SARS-CoV2(0.00727, 4.98%) |
| 9 | membrane glycoprotein->SARS-CoV2(0.00101, 8.07%) | M->Influenza(0.00682, 4.67%) |
| 10 | Nucleocapsid protein->Influenza(0.00095, 7.58%) | ORF10->SARS-CoV2(0.00452, 3.09%) |
| 11 | ORF3a->SARS-CoV2(0.00078, 6.24%) | BRLF1->EBV(0.00398, 2.73%) |
| 12 | M->Influenza(0.00076, 6.02%) | Nucleocapsid protein->Influenza(0.00387, 2.65%) |
| 13 | BMLF1->EBV(0.00058, 4.64%) | EBNA4->EBV(0.00372, 2.55%) |
| 14 | BZLF1->EBV(0.00042, 3.32%) | BMLF1->EBV(0.00372, 2.55%) |
| 15 | DNA polymerase->HBV(0.00039, 3.14%) | DNA polymerase->HBV(0.00233, 1.6%) |
| 16 | MLANA->Homo sapiens(0.00038, 3.06%) | NS4B->YFV(0.00226, 1.55%) |
| 17 | EBNA3A->EBV(0.00038, 3.0%) | ORF8->SARS-CoV2(0.00205, 1.4%) |
| 18 | EBNA4->EBV(0.00033, 2.6%) | ORF7a->SARS-CoV2(0.00195, 1.33%) |
| 19 | precore/core protein->HBV(0.00032, 2.51%) | MLANA->Homo sapiens(0.0014, 0.96%) |
| 20 | NS4B->YFV(0.0002, 1.61%) | precore/core protein->HBV(0.00107, 0.74%) |
| Total | 0.012539543 | 0.145981 |

Mean fraction and percentage composition of repertoire specificity to organisms and antigens of COVID-19 convalescent individuals (n=62). Top 10 organisms with CM/PM percentage no less than 1% is shown; top 20 antigens with CM/PM percentage no less than 0.5% is shown. The total fraction is the sum of fraction for all CM/PM sequences.

S8. Mean fraction (percentage) composition of TCR repertoire specificity of COVID-19 AZD1222 pre-vaccination individuals.

| Rank | CM:Top_Organisms (mean fraction) | PM:Top_Organisms (mean fraction) |
|---|---|---|
| 1 | SARS-CoV2(0.00594, 65.13%) | SARS-CoV2(0.0848, 68.59%) |
| 2 | CMV(0.00318, 34.94%) | CMV(0.02124, 17.18%) |
| 3 | EBV(0.00185, 20.26%) | EBV(0.01006, 8.13%) |
| 4 | Influenza(0.0014, 15.33%) | Influenza(0.00733, 5.93%) |
| 5 | Homo sapiens(0.00067, 7.33%) | Homo sapiens(0.0032, 2.59%) |
| 6 | YFV(0.00023, 2.53%) | HBV(0.00301, 2.44%) |
| 7 | HBV(0.00021, 2.29%) | YFV(0.00172, 1.39%) |
| 8 | HIV(0.00016, 1.73%) | . |
| Total | 0.009114935 | 0.123637 |

| Rank | CM:Top_Antigens (mean fraction) | PM:Top_Antigens (mean fraction) |
|---|---|---|
| 1 | ORF10->SARS-CoV2(0.00094, 10.31%) | pp65->CMV(0.007, 5.66%) |
| 2 | BMLF1->EBV(0.00092, 10.14%) | nucleocapsid phosphoprotein->SARS-CoV2(0.00636, 5.15%) |
| 3 | ORF3a->SARS-CoV2(0.00081, 8.9%) | M->Influenza(0.00376, 3.04%) |
| 4 | M->Influenza(0.00068, 7.5%) | ORF10->SARS-CoV2(0.0033, 2.67%) |
| 5 | Nucleocapsid protein->Influenza(0.00068, 7.42%) | BRLF1->EBV(0.00325, 2.63%) |
| 6 | ORF7a->SARS-CoV2(0.00056, 6.11%) | Nucleocapsid protein->Influenza(0.003, 2.43%) |
| 7 | nucleocapsid phosphoprotein->SARS-CoV2(0.00053, 5.78%) | BMLF1->EBV(0.00275, 2.23%) |
| 8 | MLANA->Homo sapiens(0.00037, 4.09%) | EBNA4->EBV(0.0024, 1.94%) |
| 9 | BRLF1->EBV(0.00036, 3.97%) | DNA polymerase->HBV(0.00224, 1.81%) |
| 10 | BZLF1->EBV(0.00035, 3.79%) | ORF7a->SARS-CoV2(0.00212, 1.71%) |
| 11 | EBNA4->EBV(0.00035, 3.78%) | ORF8->SARS-CoV2(0.00173, 1.4%) |
| 12 | EBNA3A->EBV(0.00024, 2.62%) | NS4B->YFV(0.00172, 1.39%) |
| 13 | NS4B->YFV(0.00023, 2.53%) | MLANA->Homo sapiens(0.00166, 1.34%) |
| 14 | ORF8->SARS-CoV2(0.0002, 2.16%) | BZLF1->EBV(0.00107, 0.86%) |
| 15 | ORF10->SARS-CoV2(0.00094, 10.31%) | pp65->CMV(0.007, 5.66%) |
| 16 | BMLF1->EBV(0.00092, 10.14%) | nucleocapsid phosphoprotein->SARS-CoV2(0.00636, 5.15%) |
| 17 | ORF3a->SARS-CoV2(0.00081, 8.9%) | M->Influenza(0.00376, 3.04%) |
| 18 | M->Influenza(0.00068, 7.5%) | ORF10->SARS-CoV2(0.0033, 2.67%) |
| 19 | Nucleocapsid protein->Influenza(0.00068, 7.42%) | BRLF1->EBV(0.00325, 2.63%) |
| 20 | ORF7a->SARS-CoV2(0.00056, 6.11%) | Nucleocapsid protein->Influenza(0.003, 2.43%) |
| Total | 0.009114935 | 0.123637 |

Mean fraction and percentage composition of repertoire specificity to organisms and antigens of COVID-19 AZD1222 pre-vaccination individuals (n=224). Top 10 organisms with CM/PM percentage no less than 1% is shown; top 20 antigens with CM/PM percentage no less than 0.5% is shown. The total fraction is the sum of fraction for all CM/PM sequences.

S9. Mean fraction (percentage) composition of TCR repertoire specificity of COVID-19 AZD1222 post-vaccination individuals.

| Rank | CM:Top_Organisms (mean fraction) | PM:Top_Organisms (mean fraction) |
|---|---|---|
| 1 | SARS-CoV2(0.00587, 62.64%) | SARS-CoV2(0.09802, 68.92%) |
| 2 | CMV(0.00313, 33.44%) | CMV(0.02364, 16.63%) |
| 3 | EBV(0.00183, 19.53%) | EBV(0.01199, 8.43%) |
| 4 | Influenza(0.00087, 9.3%) | Influenza(0.00865, 6.08%) |
| 5 | Homo sapiens(0.0006, 6.45%) | Homo sapiens(0.00383, 2.69%) |
| 6 | HBV(0.00037, 4.0%) | HBV(0.00326, 2.29%) |
| 7 | YFV(0.00032, 3.4%) | YFV(0.00193, 1.36%) |
| 8 | HIV(0.00014, 1.51%) | . |
| Total | 0.00936476 | 0.14222 |

| Rank | CM:Top_Antigens (mean fraction) | PM:Top_Antigens (mean fraction) |
|---|---|---|
| 1 | ORF1ab->SARS-CoV2(0.00296, 31.57%) | ORF1ab->SARS-CoV2(0.03362, 23.64%) |
| 2 | IE1->CMV(0.00207, 22.09%) | **surface glycoprotein->SARS-CoV2(0.02272, 15.97%)** |
| 3 | **surface glycoprotein->SARS-CoV2(0.00137, 14.64%)** | IE1->CMV(0.01594, 11.21%) |
| 4 | ORF7b->SARS-CoV2(0.00134, 14.31%) | ORF7b->SARS-CoV2(0.01456, 10.24%) |
| 5 | pp65->CMV(0.00115, 12.28%) | ORF3a->SARS-CoV2(0.01057, 7.43%) |
| 6 | membrane glycoprotein->SARS-CoV2(0.0009, 9.62%) | membrane glycoprotein->SARS-CoV2(0.00833, 5.85%) |
| 7 | BMLF1->EBV(0.00078, 8.29%) | pp65->CMV(0.00791, 5.56%) |
| 8 | ORF10->SARS-CoV2(0.00073, 7.76%) | nucleocapsid phosphoprotein->SARS-CoV2(0.00578, 4.06%) |
| 9 | ORF3a->SARS-CoV2(0.00062, 6.57%) | M->Influenza(0.00468, 3.29%) |
| 10 | nucleocapsid phosphoprotein->SARS-CoV2(0.00061, 6.52%) | BRLF1->EBV(0.0045, 3.17%) |
| 11 | M->Influenza(0.00057, 6.13%) | ORF10->SARS-CoV2(0.00337, 2.37%) |
| 12 | ORF7a->SARS-CoV2(0.00049, 5.26%) | Nucleocapsid protein->Influenza(0.00334, 2.35%) |
| 13 | BRLF1->EBV(0.0004, 4.28%) | BMLF1->EBV(0.00333, 2.34%) |
| 14 | BZLF1->EBV(0.00038, 4.02%) | EBNA4->EBV(0.00266, 1.87%) |
| 15 | NS4B->YFV(0.00032, 3.4%) | DNA polymerase->HBV(0.00254, 1.79%) |
| 16 | MLANA->Homo sapiens(0.00031, 3.3%) | ORF7a->SARS-CoV2(0.00238, 1.67%) |
| 17 | DNA polymerase->HBV(0.0003, 3.23%) | MLANA->Homo sapiens(0.00222, 1.56%) |
| 18 | EBNA3A->EBV(0.0003, 3.16%) | NS4B->YFV(0.00193, 1.36%) |
| 19 | Nucleocapsid protein->Influenza(0.00028, 3.03%) | ORF8->SARS-CoV2(0.00174, 1.22%) |
| 20 | ORF8->SARS-CoV2(0.00026, 2.76%) | BZLF1->EBV(0.00124, 0.87%) |
| Total | 0.00936476 | 0.14222 |

Mean fraction and percentage composition of repertoire specificity to organisms and antigens of COVID-19 AZD1222 post-vaccination individuals (n=137). Top 10 organisms with CM/PM percentage no less than 1% is shown; top 20 antigens with CM/PM percentage no less than 0.5% is shown. The total fraction is the sum of fraction for all CM/PM sequences. Significant findings are highlighted with bold font.

S10. Mean fraction (percentage) composition of TCR repertoire specificity of SLE patients.

| Rank | CM:Top_Organisms (mean fraction) | PM:Top_Organisms (mean fraction) |
|---|---|---|
| 1 | SARS-CoV2(0.01867, 67.68%) | SARS-CoV2(0.3411, 65.32%) |
| 2 | CMV(0.00842, 30.53%) | CMV(0.10501, 20.11%) |
| 3 | **HBV(0.00335, 12.13%)** | EBV(0.04061, 7.78%) |
| 4 | EBV(0.00262, 9.49%) | Influenza(0.03405, 6.52%) |
| 5 | Influenza(0.00241, 8.73%) | **HBV(0.01658, 3.18%)** |
| 6 | Homo sapiens(0.00105, 3.82%) | Homo sapiens(0.01017, 1.95%) |
| 7 | YFV(0.00071, 2.56%) | YFV(0.00883, 1.69%) |
| 8 | HIV(0.00033, 1.18%) | HCV(0.00196, 0.38%) |
| Total | **0.027590824** | **0.522186** |

| Rank | CM:Top_Antigens (mean fraction) | PM:Top_Antigens (mean fraction) |
|---|---|---|
| 1 | ORF1ab->SARS-CoV2(0.00956, 34.63%) | ORF1ab->SARS-CoV2(0.13232, 25.34%) |
| 2 | surface glycoprotein->SARS-CoV2(0.00486, 17.63%) | IE1->CMV(0.06523, 12.49%) |
| 3 | IE1->CMV(0.00447, 16.21%) | surface glycoprotein->SARS-CoV2(0.06228, 11.93%) |
| 4 | pp65->CMV(0.00391, 14.16%) | ORF7b->SARS-CoV2(0.04945, 9.47%) |
| 5 | ORF7b->SARS-CoV2(0.00389, 14.11%) | pp65->CMV(0.03939, 7.54%) |
| 6 | membrane glycoprotein->SARS-CoV2(0.00265, 9.61%) | ORF3a->SARS-CoV2(0.03154, 6.04%) |
| 7 | **precore/core protein->HBV(0.00236, 8.56%)** | nucleocapsid phosphoprotein->SARS-CoV2(0.02625, 5.03%) |
| 8 | ORF3a->SARS-CoV2(0.00236, 8.54%) | membrane glycoprotein->SARS-CoV2(0.02622, 5.02%) |
| 9 | ORF10->SARS-CoV2(0.00223, 8.09%) | M->Influenza(0.01722, 3.3%) |
| 10 | nucleocapsid phosphoprotein->SARS-CoV2(0.00202, 7.32%) | Nucleocapsid protein->Influenza(0.01596, 3.06%) |
| 11 | **DNA polymerase->HBV(0.0015, 5.44%)** | BRLF1->EBV(0.0141, 2.7%) |
| 12 | Nucleocapsid protein->Influenza(0.00129, 4.67%) | ORF10->SARS-CoV2(0.01302, 2.49%) |
| 13 | ORF7a->SARS-CoV2(0.00116, 4.21%) | **DNA polymerase->HBV(0.01176, 2.25%)** |
| 14 | M->Influenza(0.00102, 3.69%) | BMLF1->EBV(0.01111, 2.13%) |
| 15 | BMLF1->EBV(0.00085, 3.1%) | EBNA4->EBV(0.01, 1.91%) |
| 16 | EBNA4->EBV(0.00085, 3.09%) | ORF7a->SARS-CoV2(0.00961, 1.84%) |
| 17 | NS4B->YFV(0.00071, 2.56%) | NS4B->YFV(0.00883, 1.69%) |
| 18 | BRLF1->EBV(0.0007, 2.53%) | ORF8->SARS-CoV2(0.00675, 1.29%) |
| 19 | UL28->CMV(0.00059, 2.14%) | **precore/core protein->HBV(0.00645, 1.24%)** |
| 20 | ORF8->SARS-CoV2(0.00051, 1.86%) | BZLF1->EBV(0.0047, 0.9%) |
| Total | **0.027590824** | **0.522186** |

Mean fraction and percentage composition of repertoire specificity to organisms and antigens of SLE patients (n=877). Top 10 organisms with CM/PM percentage no less than 1% is shown; top 20 antigens with CM/PM percentage no less than 0.5% is shown. The total fraction is the sum of fraction for all CM/PM sequences. Significant findings are highlighted with bold font.

S11. Mean fraction (percentage) composition of TCR repertoire specificity of melanoma patients.

| Rank | CM:Top_Organisms (mean fraction) | PM:Top_Organisms (mean fraction) |
|---|---|---|
| 1 | SARS-CoV2(0.02118, 71.47%) | SARS-CoV2(0.23327, 67.52%) |
| 2 | CMV(0.01003, 33.85%) | CMV(0.06233, 18.04%) |
| 3 | Influenza(0.00536, 18.09%) | EBV(0.02646, 7.66%) |
| 4 | EBV(0.00516, 17.42%) | Influenza(0.01698, 4.91%) |
| 5 | Homo sapiens(0.00288, 9.7%) | YFV(0.00969, 2.8%) |
| 6 | HBV(0.00123, 4.16%) | Homo sapiens(0.00772, 2.23%) |
| 7 | YFV(0.00114, 3.86%) | HBV(0.00735, 2.13%) |
| Total | **0.029635652** | **0.345502** |

| Rank | CM:Top_Antigens (mean fraction) | PM:Top_Antigens (mean fraction) |
|---|---|---|
| 1 | ORF1ab->SARS-CoV2(0.01007, 33.98%) | ORF1ab->SARS-CoV2(0.09913, 28.69%) |
| 2 | pp65->CMV(0.00593, 20.0%) | surface glycoprotein->SARS-CoV2(0.05046, 14.61%) |
| 3 | surface glycoprotein->SARS-CoV2(0.00554, 18.7%) | IE1->CMV(0.03686, 10.67%) |
| 4 | IE1->CMV(0.00483, 16.29%) | ORF7b->SARS-CoV2(0.03476, 10.06%) |
| 5 | membrane glycoprotein->SARS-CoV2(0.00482, 16.26%) | pp65->CMV(0.02536, 7.34%) |
| 6 | ORF7a->SARS-CoV2(0.00415, 14.02%) | membrane glycoprotein->SARS-CoV2(0.02116, 6.12%) |
| 7 | ORF7b->SARS-CoV2(0.00347, 11.71%) | ORF3a->SARS-CoV2(0.01952, 5.65%) |
| 8 | BMLF1->EBV(0.00302, 10.18%) | nucleocapsid phosphoprotein->SARS-CoV2(0.01198, 3.47%) |
| 9 | M->Influenza(0.003, 10.13%) | BMLF1->EBV(0.0105, 3.04%) |
| 10 | **MLANA->Homo sapiens(0.0026, 8.76%)** | NS4B->YFV(0.00969, 2.8%) |
| 11 | ORF3a->SARS-CoV2(0.00255, 8.62%) | M->Influenza(0.0093, 2.69%) |
| 12 | nucleocapsid phosphoprotein->SARS-CoV2(0.00215, 7.27%) | BRLF1->EBV(0.00755, 2.18%) |
| 13 | ORF10->SARS-CoV2(0.00212, 7.14%) | Nucleocapsid protein->Influenza(0.00687, 1.99%) |
| 14 | Nucleocapsid protein->Influenza(0.00211, 7.1%) | ORF7a->SARS-CoV2(0.00667, 1.93%) |
| 15 | EBNA3A->EBV(0.00145, 4.9%) | ORF10->SARS-CoV2(0.00647, 1.87%) |
| 16 | NS4B->YFV(0.00114, 3.86%) | EBNA4->EBV(0.00516, 1.49%) |
| 17 | DNA polymerase->HBV(0.00108, 3.64%) | DNA polymerase->HBV(0.00479, 1.39%) |
| 18 | EBNA4->EBV(0.00088, 2.99%) | precore/core protein->HBV(0.00388, 1.12%) |
| 19 | EBNA6->EBV(0.0008, 2.72%) | IGF2BP2->Homo sapiens(0.00316, 0.92%) |
| 20 | **ABCD3->Homo sapiens(0.0008, 2.71%)** | ORF8->SARS-CoV2(0.00298, 0.86%) |
| Total | 0.029635652 | 0.345502 |

Mean fraction and percentage composition of repertoire specificity to organisms and antigens of melanoma patients (n=196). Top 10 organisms with CM/PM percentage no less than 1% is shown; top 20 antigens with CM/PM percentage no less than 0.5% is shown. The total fraction is the sum of fraction for all CM/PM sequences. Significant findings are highlighted with bold font.

S12. Mean fraction (percentage) composition of TCR repertoire specificity of lung cancer patients.

| Rank | CM:Top_Organisms (mean fraction) | PM:Top_Organisms (mean fraction) |
|---|---|---|
| 1 | SARS-CoV2(0.01442, 71.12%) | SARS-CoV2(0.14712, 67.75%) |
| 2 | CMV(0.00936, 46.16%) | CMV(0.03949, 18.19%) |
| 3 | EBV(0.00374, 18.45%) | EBV(0.01432, 6.59%) |
| 4 | Influenza(0.00211, 10.39%) | Influenza(0.01373, 6.32%) |
| 5 | Homo sapiens(0.00094, 4.64%) | HBV(0.00483, 2.23%) |
| 6 | HIV(0.00078, 3.85%) | Homo sapiens(0.00429, 1.98%) |
| 7 | HBV(0.0005, 2.49%) | YFV(0.00346, 1.59%) |
| 8 | YFV(0.00032, 1.59%) | HCV(0.00295, 1.36%) |
| 9 | Lymphocytic choriomeningitis virus(0.00029, 1.41%) | . |
| Total | 0.020272246 | 0.21716 |

| Rank | CM:Top_Antigens (mean fraction) | PM:Top_Antigens (mean fraction) |
|---|---|---|
| 1 | pp65->CMV(0.00616, 30.38%) | ORF1ab->SARS-CoV2(0.06005, 27.65%) |
| 2 | ORF1ab->SARS-CoV2(0.00503, 24.79%) | surface glycoprotein->SARS-CoV2(0.02506, 11.54%) |
| 3 | membrane glycoprotein->SARS-CoV2(0.00479, 23.64%) | IE1->CMV(0.02442, 11.25%) |
| 4 | IE1->CMV(0.00333, 16.42%) | ORF7b->SARS-CoV2(0.0206, 9.49%) |
| 5 | surface glycoprotein->SARS-CoV2(0.00282, 13.91%) | pp65->CMV(0.01518, 6.99%) |
| 6 | BMLF1->EBV(0.00222, 10.95%) | ORF3a->SARS-CoV2(0.01167, 5.37%) |
| 7 | ORF7b->SARS-CoV2(0.0022, 10.83%) | nucleocapsid phosphoprotein->SARS-CoV2(0.01161, 5.35%) |
| 8 | ORF3a->SARS-CoV2(0.00169, 8.34%) | membrane glycoprotein->SARS-CoV2(0.01103, 5.08%) |
| 9 | ORF10->SARS-CoV2(0.00134, 6.62%) | M->Influenza(0.00933, 4.3%) |
| 10 | nucleocapsid phosphoprotein->SARS-CoV2(0.00106, 5.25%) | ORF10->SARS-CoV2(0.00681, 3.14%) |
| 11 | M->Influenza(0.00106, 5.23%) | BRLF1->EBV(0.00529, 2.44%) |
| 12 | Nucleocapsid protein->Influenza(0.00098, 4.85%) | ORF7a->SARS-CoV2(0.00456, 2.1%) |
| 13 | MLANA->Homo sapiens(0.00069, 3.4%) | Nucleocapsid protein->Influenza(0.00384, 1.77%) |
| 14 | ORF7a->SARS-CoV2(0.00064, 3.15%) | BMLF1->EBV(0.00383, 1.76%) |
| 15 | EBNA4->EBV(0.00062, 3.05%) | NS4B->YFV(0.00346, 1.59%) |
| 16 | EBNA3A->EBV(0.00052, 2.54%) | DNA polymerase->HBV(0.00333, 1.53%) |
| 17 | BRLF1->EBV(0.00051, 2.52%) | EBNA4->EBV(0.00333, 1.53%) |
| 18 | Nef->HIV(0.00041, 2.0%) | ORF8->SARS-CoV2(0.00292, 1.34%) |
| 19 | DNA polymerase->HBV(0.00037, 1.8%) | NS3->HCV(0.00284, 1.31%) |
| 20 | ORF8->SARS-CoV2(0.00036, 1.77%) | MLANA->Homo sapiens(0.00158, 0.73%) |
| Total | 0.020272246 | 0.21716 |

Mean fraction and percentage composition of repertoire specificity to organisms and antigens of lung cancer patients (n=209). Top 10 organisms with CM/PM percentage no less than 1% is shown; top 20 antigens with CM/PM percentage no less than 0.5% is shown. The total fraction is the sum of fraction for all CM/PM sequences.

S13. Mean fraction (percentage) composition of TCR repertoire specificity of breast cancer patients.

| Rank | CM:Top_Organisms (mean fraction) | PM:Top_Organisms (mean fraction) |
|---|---|---|
| 1 | **Influenza(0.00928, 45.65%)** | SARS-CoV2(0.14779, 70.53%) |
| 2 | SARS-CoV2(0.00897, 44.15%) | CMV(0.03781, 18.04%) |
| 3 | CMV(0.0054, 26.56%) | EBV(0.0155, 7.4%) |
| 4 | EBV(0.00158, 7.79%) | Influenza(0.01424, 6.8%) |
| 5 | Homo sapiens(0.00059, 2.9%) | HBV(0.00413, 1.97%) |
| 6 | YFV(0.00037, 1.82%) | YFV(0.0031, 1.48%) |
| 7 | HBV(0.00025, 1.22%) | Homo sapiens(0.00229, 1.09%) |
| 8 | . | HCV(0.00209, 1.0%) |
| Total | 0.020317835 | 0.209561 |

| Rank | CM:Top_Antigens (mean fraction) | PM:Top_Antigens (mean fraction) |
|---|---|---|
| 1 | M->Influenza(0.00713, 35.11%) | ORF1ab->SARS-CoV2(0.04942, 23.58%) |
| 2 | ORF1ab->SARS-CoV2(0.00423, 20.84%) | surface glycoprotein->SARS-CoV2(0.0378, 18.04%) |
| 3 | pp65->CMV(0.00285, 14.03%) | IE1->CMV(0.02137, 10.2%) |
| 4 | surface glycoprotein->SARS-CoV2(0.00276, 13.58%) | ORF7b->SARS-CoV2(0.0194, 9.26%) |
| 5 | IE1->CMV(0.00269, 13.22%) | pp65->CMV(0.01677, 8.0%) |
| 6 | Nucleocapsid protein->Influenza(0.0021, 10.32%) | membrane glycoprotein->SARS-CoV2(0.01496, 7.14%) |
| 7 | ORF10->SARS-CoV2(0.00152, 7.48%) | ORF3a->SARS-CoV2(0.01154, 5.51%) |
| 8 | ORF7b->SARS-CoV2(0.00148, 7.28%) | nucleocapsid phosphoprotein->SARS-CoV2(0.00926, 4.42%) |
| 9 | membrane glycoprotein->SARS-CoV2(0.00148, 7.28%) | Nucleocapsid protein->Influenza(0.00709, 3.38%) |
| 10 | ORF3a->SARS-CoV2(0.00068, 3.33%) | BRLF1->EBV(0.00673, 3.21%) |
| 11 | BMLF1->EBV(0.00062, 3.05%) | M->Influenza(0.00673, 3.21%) |
| 12 | nucleocapsid phosphoprotein->SARS-CoV2(0.00056, 2.74%) | ORF10->SARS-CoV2(0.00635, 3.03%) |
| 13 | EBNA4->EBV(0.00048, 2.36%) | BMLF1->EBV(0.0047, 2.24%) |
| 14 | ORF7a->SARS-CoV2(0.00039, 1.92%) | ORF7a->SARS-CoV2(0.00351, 1.68%) |
| 15 | BZLF1->EBV(0.00039, 1.91%) | DNA polymerase->HBV(0.00316, 1.51%) |
| 16 | NS4B->YFV(0.00037, 1.82%) | NS4B->YFV(0.0031, 1.48%) |
| 17 | MLANA->Homo sapiens(0.00029, 1.42%) | EBNA4->EBV(0.00294, 1.41%) |
| 18 | BRLF1->EBV(0.00023, 1.15%) | NS3->HCV(0.00206, 0.99%) |
| 19 | ORF6->SARS-CoV2(0.00022, 1.09%) | ORF8->SARS-CoV2(0.0018, 0.86%) |
| 20 | EBNA3A->EBV(0.00022, 1.06%) | ORF6->SARS-CoV2(0.00128, 0.61%) |
| Total | 0.020317835 | 0.209561 |

Mean fraction and percentage composition of repertoire specificity to organisms and antigens of breast cancer patients (n=73). Top 10 organisms with CM/PM percentage no less than 1% is shown; top 20 antigens with CM/PM percentage no less than 0.5% is shown. The total fraction is the sum of fraction for all CM/PM sequences. Significant findings are highlighted with bold font.

S14. Mean fraction (percentage) composition of TCR repertoire specificity of urothelial bladder cancer patients.

| Rank | CM:Top_Organisms (mean fraction) | PM:Top_Organisms (mean fraction) |
|---|---|---|
| 1 | SARS-CoV2(0.01621, 87.59%) | SARS-CoV2(0.09953, 65.67%) |
| 2 | CMV(0.00293, 15.86%) | CMV(0.03036, 20.03%) |
| 3 | Influenza(0.00151, 8.15%) | EBV(0.01454, 9.59%) |
| 4 | EBV(0.00128, 6.93%) | Influenza(0.00573, 3.78%) |
| 5 | Homo sapiens(0.00044, 2.4%) | YFV(0.00413, 2.73%) |
| 6 | HBV(0.00023, 1.24%) | HBV(0.00352, 2.32%) |
| 7 | YFV(0.00012, 0.62%) | Homo sapiens(0.00315, 2.08%) |
| Total | 0.018502 | 0.151555 |

| Rank | CM:Top_Antigens (mean fraction) | PM:Top_Antigens (mean fraction) |
|---|---|---|
| 1 | surface glycoprotein->SARS-CoV2(0.01043, 56.39%) | ORF1ab->SARS-CoV2(0.03692, 24.36%) |
| 2 | ORF1ab->SARS-CoV2(0.00907, 49.03%) | surface glycoprotein->SARS-CoV2(0.02088, 13.78%) |
| 3 | membrane glycoprotein->SARS-CoV2(0.00279, 15.05%) | IE1->CMV(0.01877, 12.38%) |
| 4 | pp65->CMV(0.00192, 10.39%) | ORF7b->SARS-CoV2(0.01682, 11.1%) |
| 5 | ORF10->SARS-CoV2(0.00171, 9.26%) | pp65->CMV(0.01147, 7.57%) |
| 6 | ORF7b->SARS-CoV2(0.00136, 7.35%) | ORF3a->SARS-CoV2(0.00971, 6.41%) |
| 7 | IE1->CMV(0.00136, 7.33%) | BRLF1->EBV(0.00729, 4.81%) |
| 8 | ORF3a->SARS-CoV2(0.00123, 6.64%) | membrane glycoprotein->SARS-CoV2(0.00659, 4.35%) |
| 9 | M->Influenza(0.00116, 6.28%) | nucleocapsid phosphoprotein->SARS-CoV2(0.00635, 4.19%) |
| 10 | BMLF1->EBV(0.00072, 3.89%) | ORF10->SARS-CoV2(0.00456, 3.01%) |
| 11 | nucleocapsid phosphoprotein->SARS-CoV2(0.00059, 3.17%) | NS4B->YFV(0.00413, 2.73%) |
| 12 | ORF6->SARS-CoV2(0.00053, 2.87%) | M->Influenza(0.00384, 2.53%) |
| 13 | Nucleocapsid protein->Influenza(0.00031, 1.69%) | EBNA4->EBV(0.0033, 2.17%) |
| 14 | EBNA4->EBV(0.00027, 1.46%) | ORF7a->SARS-CoV2(0.00288, 1.9%) |
| 15 | MLANA->Homo sapiens(0.00026, 1.4%) | DNA polymerase->HBV(0.00272, 1.8%) |
| 16 | ORF7a->SARS-CoV2(0.00025, 1.35%) | BMLF1->EBV(0.00263, 1.73%) |
| 17 | ORF8->SARS-CoV2(0.00023, 1.25%) | Nucleocapsid protein->Influenza(0.00186, 1.23%) |
| 18 | BZLF1->EBV(0.0002, 1.1%) | MLANA->Homo sapiens(0.00161, 1.06%) |
| 19 | precore/core protein->HBV(0.00014, 0.76%) | BZLF1->EBV(0.00118, 0.78%) |
| 20 | NY-ESO-1->Homo sapiens(0.00012, 0.64%) | ORF8->SARS-CoV2(0.00117, 0.77%) |
| Total | 0.018502 | 0.151555 |

Mean fraction and percentage composition of repertoire specificity to organisms and antigens of urothelial bladder cancer patients (n=90). Top 10 organisms with CM/PM percentage no less than 1% is shown; top 20 antigens with CM/PM percentage no less than 0.5% is shown. The total fraction is the sum of fraction for all CM/PM sequences.

S15. Mean fraction (percentage) composition of TCR repertoire specificity of all cancer patients.

| Rank | CM:Top_Organisms (mean fraction) | PM:Top_Organisms (mean fraction) |
|---|---|---|
| 1 | SARS-CoV2(0.01574, 71.5%) | SARS-CoV2(0.17247, 67.79%) |
| 2 | CMV(0.00752, 34.17%) | CMV(0.04719, 18.55%) |
| 3 | Influenza(0.00363, 16.48%) | EBV(0.01897, 7.46%) |
| 4 | EBV(0.00341, 15.48%) | Influenza(0.01422, 5.59%) |
| 5 | Homo sapiens(0.00135, 6.12%) | YFV(0.0055, 2.16%) |
| 6 | HBV(0.00073, 3.31%) | HBV(0.00544, 2.14%) |
| 7 | YFV(0.00051, 2.34%) | Homo sapiens(0.0047, 1.85%) |
| 8 | HIV(0.00039, 1.78%) | HCV(0.0018, 0.71%) |
| Total | 0.022010816 | 0.254415 |

| Rank | CM:Top_Antigens (mean fraction) | PM:Top_Antigens (mean fraction) |
|---|---|---|
| 1 | ORF1ab->SARS-CoV2(0.00692, 31.43%) | ORF1ab->SARS-CoV2(0.06842, 26.89%) |
| 2 | surface glycoprotein->SARS-CoV2(0.00466, 21.19%) | surface glycoprotein->SARS-CoV2(0.03431, 13.48%) |
| 3 | pp65->CMV(0.00455, 20.67%) | IE1->CMV(0.02868, 11.27%) |
| 4 | membrane glycoprotein->SARS-CoV2(0.00413, 18.76%) | ORF7b->SARS-CoV2(0.02463, 9.68%) |
| 5 | IE1->CMV(0.0033, 14.99%) | pp65->CMV(0.01857, 7.3%) |
| 6 | ORF7b->SARS-CoV2(0.00244, 11.08%) | ORF3a->SARS-CoV2(0.01594, 6.27%) |
| 7 | M->Influenza(0.00228, 10.38%) | membrane glycoprotein->SARS-CoV2(0.01416, 5.57%) |
| 8 | BMLF1->EBV(0.00176, 8.01%) | nucleocapsid phosphoprotein->SARS-CoV2(0.01156, 4.54%) |
| 9 | ORF7a->SARS-CoV2(0.00175, 7.93%) | M->Influenza(0.00872, 3.43%) |
| 10 | ORF10->SARS-CoV2(0.00173, 7.87%) | ORF10->SARS-CoV2(0.00682, 2.68%) |
| 11 | ORF3a->SARS-CoV2(0.00168, 7.62%) | BRLF1->EBV(0.00638, 2.51%) |
| 12 | nucleocapsid phosphoprotein->SARS-CoV2(0.00152, 6.9%) | BMLF1->EBV(0.00593, 2.33%) |
| 13 | Nucleocapsid protein->Influenza(0.00124, 5.62%) | NS4B->YFV(0.0055, 2.16%) |
| 14 | MLANA->Homo sapiens(0.00105, 4.79%) | Nucleocapsid protein->Influenza(0.00502, 1.97%) |
| 15 | EBNA3A->EBV(0.00065, 2.95%) | ORF7a->SARS-CoV2(0.00478, 1.88%) |
| 16 | DNA polymerase->HBV(0.00061, 2.77%) | EBNA4->EBV(0.00428, 1.68%) |
| 17 | EBNA4->EBV(0.0006, 2.71%) | DNA polymerase->HBV(0.0038, 1.49%) |
| 18 | BRLF1->EBV(0.00058, 2.66%) | ORF8->SARS-CoV2(0.00273, 1.07%) |
| 19 | ORF8->SARS-CoV2(0.00052, 2.37%) | BZLF1->EBV(0.00203, 0.8%) |
| 20 | NS4B->YFV(0.00051, 2.34%) | precore/core protein->HBV(0.002, 0.79%) |
| Total | 0.022010816 | 0.254415 |

Mean fraction and percentage composition of repertoire specificity to organisms and antigens of all cancer patients (n=686). Top 10 organisms with CM/PM percentage no less than 1% is shown; top 20 antigens with CM/PM percentage no less than 0.5% is shown. The total fraction is the sum of fraction for all CM/PM sequences.

S16. ANOVA test p-values for the 7 numeric features on different comparisons of sub-populations.

| Comparison | SARS-CoV2 | CMV | Influenza | EBV | HBV | Med | High |
|---|---|---|---|---|---|---|---|
| CMV+ vs CMV- | 2.25E-32 | 1.36E-22 | 1.76E-09 | 1.16E-11 | 0.000002 | 5.33E-21 | 4.81E-33 |
| SLE vs Healthy | 0 | 0 | 1.23E-146 | 8.88E-197 | 7.12E-86 | 0 | 9.94E-49 |
| Melanoma vs Control | 1.42E-35 | 1.23E-26 | 1.18E-11 | 7.33E-10 | 0.000002 | 4.44E-36 | 3.25E-20 |
| Lung cancer vs Control | 3.45E-15 | 8.69E-12 | 0.000704 | 0.000045 | 0.001242 | 2.70E-14 | 1.35E-09 |
| Breast cancer vs Control | 2.84E-07 | 3.04E-07 | 9.29E-09 | 0.008046 | 0.109701 | 2.96E-09 | 0.024077 |
| Urothelial bladder cancer vs Control | 0.007669 | 0.005549 | 0.23278 | 0.02039 | 0.47932 | 0.007451 | 0.019181 |
| Other cancers vs Control | 5.02E-17 | 1.73E-12 | 0.000002 | 5.44E-09 | 0.000116 | 1.77E-12 | 4.22E-12 |
| Melanoma vs Lung cancer vs Breast cancer vs Urothelial cancer vs Other cancers vs Control | 8.13E-40 | 8.54E-25 | 1.14E-10 | 2.16E-13 | 6.74E-07 | 1.22E-32 | 8.39E-25 |

Med = medium-frequency clonotype fraction; High = high-frequency clonotype fraction.
